# Monogenic and polygenic inheritance become instruments for clonal selection

**DOI:** 10.1101/653691

**Authors:** Po-Ru Loh, Giulio Genovese, Steven A McCarroll

**Affiliations:** Division of Genetics, Department of Medicine, Brigham and Women’s Hospital and Harvard Medical School; Program in Medical and Population Genetics, Broad Institute of MIT and Harvard; Stanley Center for Psychiatric Research, Broad Institute of MIT and Harvard; Department of Genetics, Harvard Medical School

## Abstract

Clonally expanded blood cells with somatic mutations (clonal hematopoiesis, CH) are commonly acquired with age and increase risk of later blood cancer. To identify genes and mutations that give selective advantage to mutant clones, we identified among 482,789 UK Biobank participants some 19,632 autosomal mosaic chromosomal alterations (mCAs), including deletions, duplications, and copy number-neutral loss of heterozygosity (CNN-LOH). Analysis of these acquired mutations, along with inherited genetic variation, revealed 52 inherited, rare, large-effect coding or splice variants (in seven genes) that greatly (odds ratios of 11 to 758) increased vulnerability to CH with specific acquired CNN-LOH mutations. Acquired mutations systematically replaced the inherited risk alleles (at *MPL*) or duplicated them to the homologous chromosome (at *FH, NBN, MRE11, ATM, SH2B3*, and *TM2D3*). Three of the seven genes (*MRE11, NBN*, and *ATM*) encode components of the MRN-ATM pathway, which limits cell division after DNA damage and telomere attrition; another two (*MPL, SH2B3*) encode proteins that regulate stem cell self-renewal. In addition to these monogenic inherited forms of CH, we found a common and surprisingly polygenic form: CNN-LOH mutations across the genome tended to cause chromosomal segments with alleles that promote hematopoietic cell proliferation to replace their homologous (allelic) counterparts, increasing polygenic drive for blood-cell proliferation traits. This dynamic reveals a challenge for lifelong cytopoiesis in any genetically diverse species: individuals inherit unequal proliferative genetic potentials on paternally and maternally derived chromosome-pairs, and readily-acquired mutations that replace chromosomal segments with their homologous counterparts give selective advantage to mutant cells.

---

Clonal expansion of blood cells with acquired mutations (“clonal hematopoiesis,” CH) often arises as humans age [1–10]. Clones increase risk of future blood cancers, which can arise from subclones that acquire additional mutations [2–5, 9, 10]. The identities and functions of the mutations in expanded clones could help reveal the early selective pressures that lead to clonality and underlie pre-cancerous states. The blood clones identified to date contain many different largescale mosaic chromosomal alterations (mCAs: deletions, duplications, and copy-number neutral loss of heterozygosity [CNN-LOH]) on all chromosomes, but it remains unknown which genes and variants drive selection of most clones.

To address this question using data from a large number of people with CH, we identified 17,111 CH cases involving 19,632 autosomal mCAs (Fig. 1, Supplementary Figures 1–22, and Supplementary Tables 1–3) by analyzing SNP-array intensity data from 482,789 UK Biobank (UKB) participants 40–70 years of age [11, 12]. To identify these cases, we applied a method we recently described and applied to a smaller interim UKB release [10]; our approach finds imbalances in the abundances of homologous chromosomal segments, which are identified by combining allele-specific intensity data with long-range chromosomal phase information [13, 14]. We classified 73% of the detected mCAs as either loss (3,718 events), gain (2,389 events), or CNNLOH (8,185 events), the replacement of one chromosomal segment by its homologous (allelic) counterpart (Supplementary Table 1). (Another 5,340 mCAs could not be confidently classified, as power to detect imbalances exceeded power to distinguish copy-neutral from copy-number-altering mCAs [10]; Supplementary Fig. 23.) Of the 19,632 detected mCAs, 12,683 were present at cell fractions from 0.5 to 5%, and 6,949 were present at cell fractions >5% (Supplementary Fig. 24). Consistent with previous work [2, 3, 6, 7, 10], mCAs on different chromosomes exhibited different recurrence rates (Fig. 1 and Supplementary Fig. 25) and a range of tendencies to be more common among males and the elderly (Supplementary Fig. 26 and Supplementary Table 4). Clones also tended to be found in individuals with anomalous counts for specific populations of blood cells (Supplementary Fig. 27 and Supplementary Table 5).

**Table 1.** Associations of mosaic CNN-LOH mutations with inherited rare coding or splice variants in *cis*.

| Arm | Gene | Position<sup>a</sup> | Variant | Effect<sup>b</sup> | Alleles<sup>c</sup> | AF<sup>d</sup> | GWAS |  | Allelic shift in hets |  |  |
| --- | --- | --- | --- | --- | --- | --- | --- | --- | --- | --- | --- |
|  |  |  |  |  |  |  | *P* | OR (95% CI) | *N*<sub>REF</sub><sup>e</sup> | *N*<sub>ALT</sub> | *P* |
| Novel loci at which rare variants associate with CNN-LOH events in *cis* |  |  |  |  |  |  |  |  |  |  |  |
| 1q | *FH* | 241675301 | rs199822819 | missense | G/C | 0.0003 | 4.9×10<sup>−11</sup> | 28 (14–55) | 1 | 8 | 0.039 |
| 8q | *NBN* | 90983441 | rs1187082186 | frameshift | ATTTGT/A | 0.0002 | 4.8×10<sup>−13</sup> | 210 (92–484) | 0 | 6 | 0.031 |
| 11q | *MRE11* | 94189489 | rs587781384 | stop gained | C/A | 4×10<sup>−5</sup> | 5.6×10<sup>−10</sup> | 130 (50–338) | 0 | 5 | 0.062 |
| 12q | *SH2B3* | 111885310 | rs72650673 | missense | G/A | 0.002 | 3.1×10<sup>−8</sup> | 11 (5.8–20) | 1 | 8 | 0.039 |
| Previously reported loci at which rare variants associate with CNN-LOH events in *cis* |  |  |  |  |  |  |  |  |  |  |  |
| 1p | *MPL* | 43804305 | rs28928907 | missense | G/C | 0.0006 | 1.9×10<sup>−130</sup> | 142 (111–184) | 70 | 0 | 1.7×10<sup>−21</sup> |
| 11q | *ATM* | 108172425 | rs587779844 | missense | C/T | 0.0001 | 3.5×10<sup>−20</sup> | 96 (52–177) | 0 | 12 | 0.00049 |
| 15q | *TM2D3* | 102151467 | 70kb del<sup>f</sup> | gene deletion | ref/del | 0.0003 | 9.8×10<sup>−224</sup> | 555 (425–724) | 2 | 110 | 2.4×10<sup>−30</sup> |
| Additional independently associated likely causal coding or splice variants |  |  |  |  |  |  |  |  |  |  |  |
| 1p | *MPL* | 43803600 | rs146249964 | splice donor | T/A | 0.0001 | 2.8×10<sup>−23</sup> | 97 (55–171) | 12 | 0 | 0.00049 |
|  |  | 43803817 | rs148434485 | stop gained | C/T | 2×10<sup>−5</sup> | 1.6×10<sup>−6</sup> | 128 (37–446) | 2 | 0 | 0.5 |
|  |  | 43803824 | rs145714475 | missense | T/C | 2×10<sup>−5</sup> | 1.9×10<sup>−6</sup> | 120 (35–414) | 3 | 0 | 0.25 |
|  |  | 43803903 | rs142565191 | splice donor | G/A | 4×10<sup>−5</sup> | 7.5×10<sup>−6</sup> | 72 (22–238) | 3 | 0 | 0.25 |
|  |  | 43804234 | rs587778514 | frameshift | CCT/C | 1×10<sup>−5</sup> | 3.9×10<sup>−5</sup> | 199 (40–987) | 2 | 0 | 0.5 |
|  |  | 43804375 | rs587778515 | frameshift | CT/C | 0.0002 | 7.0×10<sup>−41</sup> | 105 (68–161) | 24 | 0 | 1.2×10<sup>−7</sup> |
|  |  | 43804396 | rs752453717 | splice modifier | G/C | 0.0003 | 5.8×10<sup>−36</sup> | 74 (48–113) | 24 | 0 | 1.2×10<sup>−7</sup> |
|  |  | 43804957 | rs764904424 | missense | C/G | 0.0001 | 2.1×10<sup>−8</sup> | 35 (15–79) | 6 | 0 | 0.031 |
|  |  | 43805052 | rs6088 | missense | G/A | 9×10<sup>−5</sup> | 8.3×10<sup>−10</sup> | 61 (26–141) | 6 | 0 | 0.031 |
|  |  | 43805656 | rs144210383 | missense | G/T | 0.0001 | 5.3×10<sup>−9</sup> | 44 (19–101) | 6 | 0 | 0.031 |
|  |  | 43805713 | rs121913611 | missense | C/T | 0.0002 | 3.3×10<sup>−28</sup> | 102 (61–171) | 17 | 0 | 1.5×10<sup>−5</sup> |
|  |  | 43812115 | rs769297582 | splice acceptor | G/C | 2×10<sup>−5</sup> | 5.1×10<sup>−7</sup> | 199 (54–737) | 3 | 0 | 0.25 |
|  |  | 43814627 | rs754859909 | stop gained | G/A | 7×10<sup>−5</sup> | 1.7×10<sup>−16</sup> | 126 (61–258) | 9 | 0 | 0.0039 |
|  |  | 43814729 | 454bp del<sup>g</sup> | exon 10 deletion | ref/del | 0.0002 | 3.6×10<sup>−58</sup> | 153 (104–225) | 31 | 0 | 9.3×10<sup>−10</sup> |
|  |  | 43817942 | rs369156948 | stop gained | C/T | 3×10<sup>−5</sup> | 4.8×10<sup>−8</sup> | 114 (39–333) | 4 | 0 | 0.12 |
|  |  | 43817973 | rs971379181 | frameshift | CG/C | 3×10<sup>−5</sup> | 5.8×10<sup>−13</sup> | 240 (93–618) | 6 | 0 | 0.031 |
| 8q | *NBN* | 90983420 | rs777460725 | missense | A/C | 0.0001 | 8.1×10<sup>−5</sup> | 114 (28–465) | 0 | 2 | 0.5 |
| 11q | *ATM* | 108127067 | rs1137887 | splice modifier | G/A | 4×10<sup>−5</sup> | 9.6×10<sup>−6</sup> | 65 (20–214) | 0 | 2 | 0.5 |
|  |  | 108141801 | rs786203054 | missense | T/G | 7×10<sup>−6</sup> | 1.2×10<sup>−5</sup> | 437 (73–2618) | 0 | 2 | 0.5 |
|  |  | 108155007 | rs781357995 | frameshift | AG/A | 0.0001 | 3.0×10<sup>−9</sup> | 48 (21–111) | 0 | 6 | 0.031 |
|  |  | 108175528 | rs376603775 | stop gained | C/T | 6×10<sup>−5</sup> | 2.8×10<sup>−5</sup> | 44 (14–143) | 0 | 4 | 0.12 |
|  |  | 108179837 | rs774925473 | splice modifier | A/G | 8×10<sup>−5</sup> | 6.8×10<sup>−5</sup> | 33 (10–104) | 0 | 3 | 0.25 |
|  |  | 108181006 | rs56399311 | missense | A/G | 8×10<sup>−5</sup> | 1.7×10<sup>−6</sup> | 44 (16–120) | 0 | 4 | 0.12 |
|  |  | 108201108 | rs56399857 | missense | T/G | 0.0002 | 4.9×10<sup>−5</sup> | 18 (6.6–48) | 0 | 4 | 0.12 |
|  |  | 108202611 | rs587776547<sup>h</sup> | inframe deletion | C...T/C | 7×10<sup>−5</sup> | 8.5×10<sup>−9</sup> | 73 (29–183) | 0 | 5 | 0.062 |
|  |  | 108216545 | rs587779872 | missense | C/T | 2×10<sup>−5</sup> | 3.6×10<sup>−11</sup> | 251 (89–706) | 0 | 5 | 0.062 |
| 12q | *SH2B3* | 111885295 | rs148636776 | missense | G/A | 0.0004 | 4.0×10<sup>−5</sup> | 19 (7–50) | 0 | 5 | 0.062 |
| 15q | *TM2D3* | 102182739 | rs113189685 | missense | G/T | 3×10<sup>−5</sup> | 2.8×10<sup>−8</sup> | 132 (45–389) | 1 | 3 | 0.62 |
|  |  | 102182749 | rs754640606 | missense | G/C | 5×10<sup>−5</sup> | 1.2×10<sup>−40</sup> | 544 (289–1025) | 0 | 19 | 3.8×10<sup>−6</sup> |
|  |  | 102182761 | rs976377433 | missense | A/G | 3×10<sup>−5</sup> | 2.3×10<sup>−8</sup> | 140 (47–413) | 0 | 4 | 0.12 |
|  |  | 102190214 | rs768556490 | frameshift | G/GT | 3×10<sup>−5</sup> | 8.2×10<sup>−29</sup> | 758 (327–1759) | 1 | 11 | 0.0063 |

Results of two independent statistical tests are reported: (i) a Fisher test treating individuals with a mosaic CNN-LOH mutation in *cis* as cases; and (ii) a binomial test for biased allelic imbalance in heterozygous cases. Loci reaching genome-wide significance in the first test are reported. At these loci, additional independently associated coding or splice variants reaching Bonferroni significance are also reported. For full details of statistical tests, see Methods. (See next page for table footnotes.)

**Figure 1.**
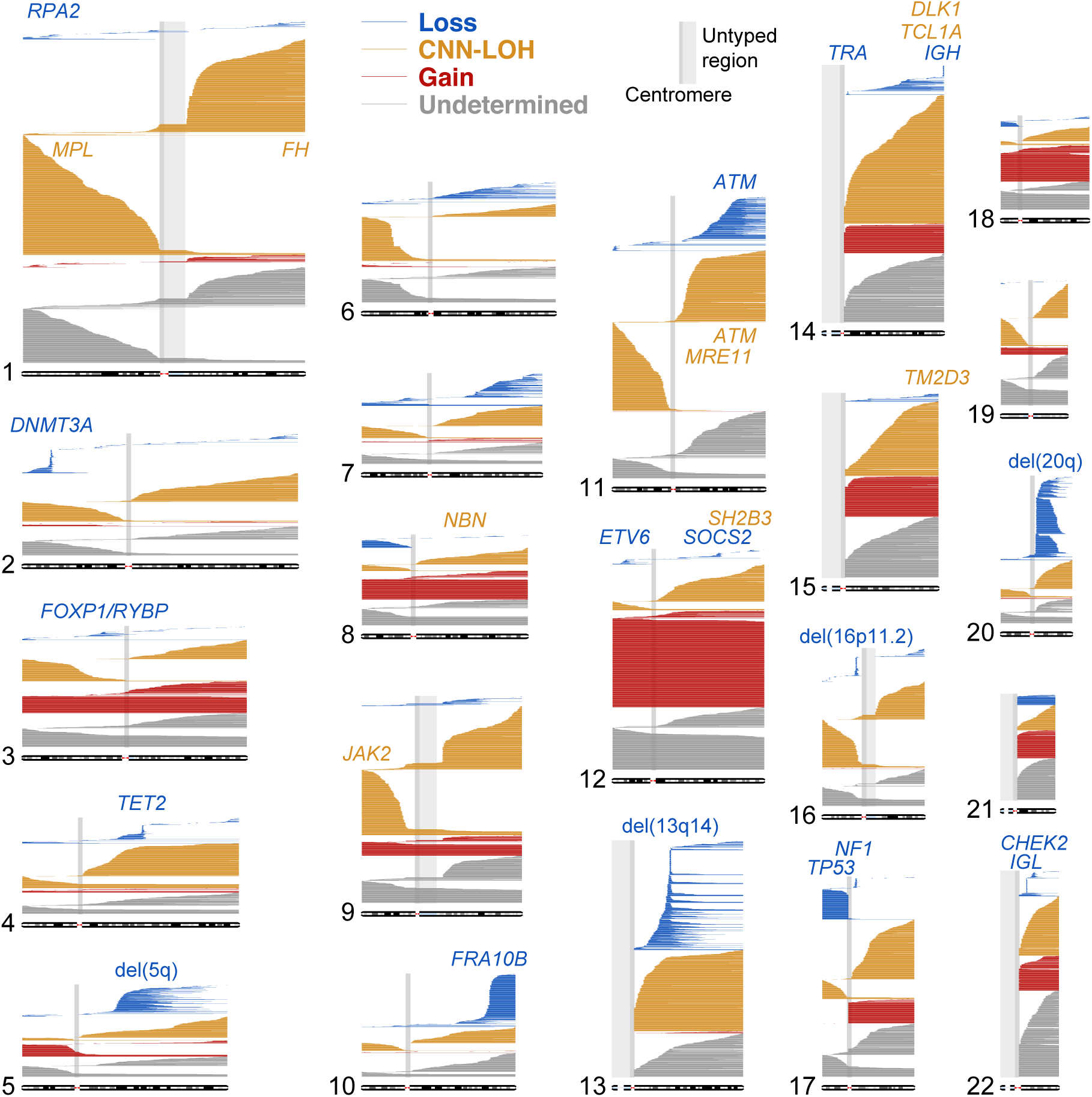
Mosaic chromosomal alterations detected among 482,789 UK Biobank participants. Each horizontal line corresponds to an mCA; a total of 19,632 autosomal events in 17,111 unique individuals are displayed. Detected events are color-coded by copy number of the affected chromosome or segment (orange, LOH; blue, loss/deletion; red, gain/duplication). Focal deletions are labeled in blue with the names of putative target genes. Loci containing inherited variants influencing somatic events in *cis* are labeled in the same color as the corresponding mCA (orange for CNN-LOH-associated loci, blue for losses). Enlarged per-chromosome plots are provided in Supplementary Figures 1–22.

We next sought to identify specific genes and variants that propel the selection of clones. We recently identified three loci (*MPL, ATM*, and *TM2D3–TARSL2*) at which inherited rare variants increase the risk of developing clones with acquired CNN-LOH mutations that affect the rare inherited risk allele in a predictable way [10]. To detect loci targeted by CNN-LOH mutations in this manner, and to identify causal inherited variants at these loci, we searched the genome for associations between inherited variants and CNN-LOH mutations acquired in *cis*.

Inherited rare variants at seven loci (*MPL, ATM, TM2D3, FH, NBN, MRE11*, and *SH2B3*) associated with the development of blood clones in which an acquired CNN-LOH mutation had affected the inherited risk allele in a consistent way (Table 1 and Fig. 2). At six loci (all loci other than *MPL*), the inherited rare alleles were in almost every case made homozygous by somatic CNN-LOH mutations (149 of 153 cases; binomial *P* =3.9×10^−39^). All the associations were driven by rare coding variants with large effect sizes (ORs 11–555; 95% CIs 5.8–724): the lead associated variants at six of the seven loci were coding mutations, and the lead variant at the remaining locus, *MRE11* (rs762019591; Fisher’s exact *P* =3.0×10^−11^), tagged a nonsense SNP in *MRE11* (rs587781384; Table 1).

**Figure 2.**
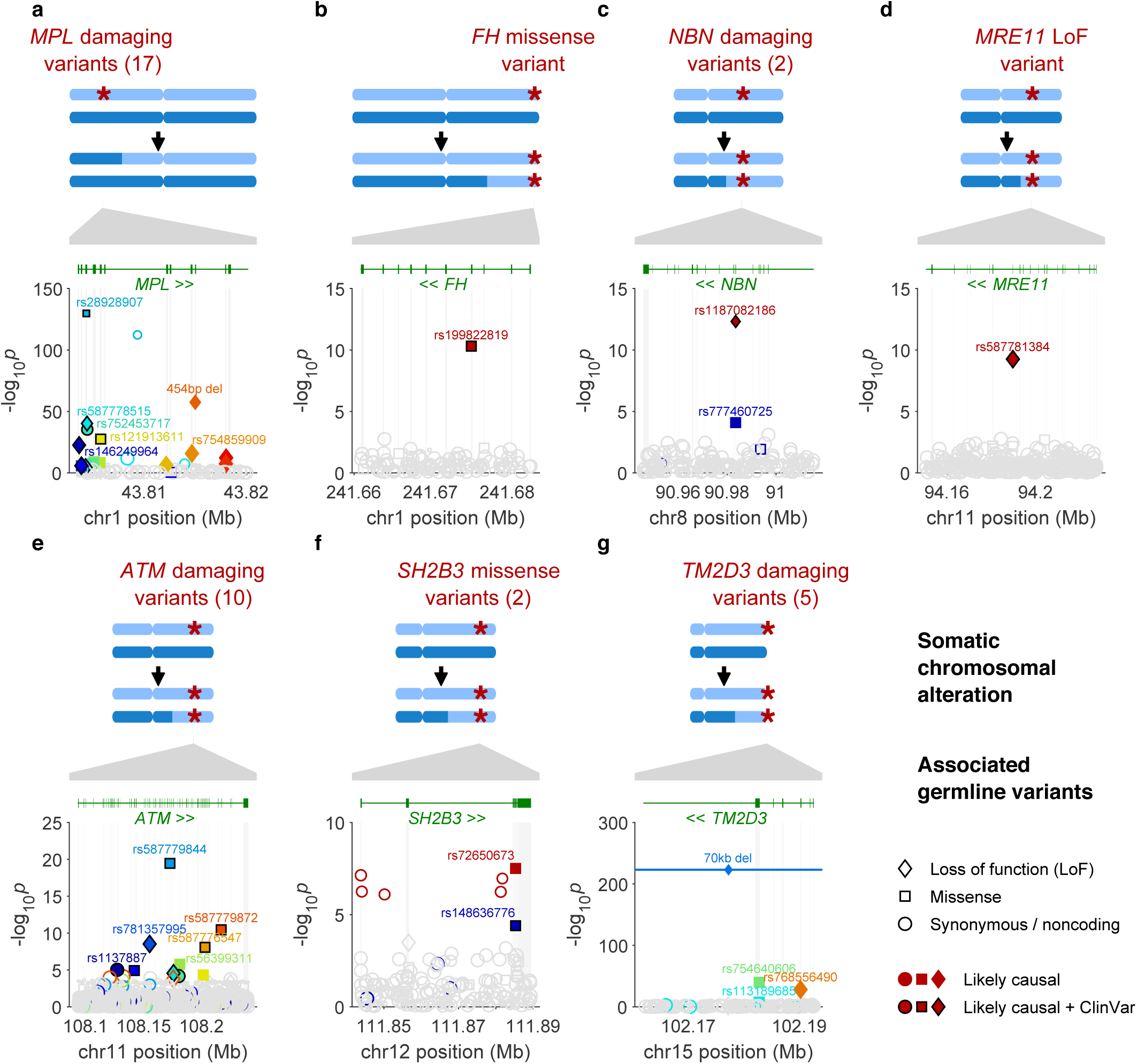
Fine-mapped inherited sequence alleles associated with the acquisition/selection of CNN-LOH mutations in *cis*. (**a**) *MPL*, (**b**) *FH*, (**c**) *NBN*, (**d**) *MRE11*, (**e**) *ATM*, (**f**) *SH2B3*, (**g**) *TM2D3*. At each locus, the CNN-LOH mutations acquired by expanded clones tend to have deleted (**a**) or duplicated (**b**–**g**) the inherited alleles in a predictable manner as shown. Each panel is organized in the following way: top, genomic modifications observed in clones; bottom, association *P*-values (Fisher’s exact test) vs. chromosomal position. All variants with filled symbols are likely causal coding or splice variants (Table 1); black marker edges indicate evidence of pathogenicity in ClinVar [21]. Distinct colors are used to indicate the statistical independence of variants; any variants in linkage disequilibrium with likely causal variants (*R*^2^>0.2 in cases) are indicated with open symbols with a border color matching that of the likely causal variant. Symbol shapes indicate the effects of the indicated variant on encoded protein (LoF, missense, etc.); symbol sizes scale inversely with minor allele frequency.

The functions of five of the seven implicated genes converged upon two likely mechanisms of clonal advantage. Three implicated genes (*MRE11, NBN, ATM*) encode proteins that act together to limit cell growth after DNA damage and telomere attrition [15]. Specifically, *MRE11* and *NBN* encode two of the three proteins in the MRN complex, which recognizes double-strand breaks and activates the checkpoint kinase encoded by *ATM* [16,17]. Thus, strong-effect mutations at *MRE11, NBN*, and *ATM* (made homozygous by CNN-LOH; Fig. 2c,d,e) disrupt a key pathway that limits proliferation in cells that have experienced DNA damage or telomere shortening.

Two other genes underlying monogenic-inherited vulnerability to CH encode proteins that regulate stem cell self-renewal: *MPL*, encoding the thrombopoietin receptor; and *SH2B3*, encoding a signaling protein (LNK) that negatively regulates hematopoietic signaling through MPL [18]. The primary *SH2B3* risk allele (rs72650673:A) increases platelet counts in carriers [19], consistent with impaired inhibition of hematopoietic signaling; this same missense allele had been made homozygous by clonally selected CNN-LOH mutations at *SH2B3* (Fig. 2f). Clonally selected CNN-LOH mutations at *MPL* acted in the opposite direction, eliminating rare inherited variants that reduce *MPL* function (Fig. 2a) [10].

To identify additional variants at all seven loci for which CNN-LOH mutations led to subsequent clonal proliferation, we performed fine-mapping analyses comprehensively examining coding and splice variants in these genes by integrating information from SNP arrays, imputation, and exome sequencing (Methods, Supplementary Figures 28–30, and Supplementary Note). Among 616 missense, loss-of-function (LoF) [20], or likely pathogenic [21] variants tested (Methods), 52 variants associated independently with CNN-LOH mosaicism in *cis* (FDR<0.05 per locus; ORs 11–758, 95% CIs 4–2,618), including multiple variants in *MPL* (28 variants), *ATM* (13), *TM2D3*(5), *NBN* (2), and *SH2B3* (2); 38 of the 52 variants reached Bonferroni significance (Fisher’s exact *P*<8.1×10^−5^ for 616 variants tested; Table 1, Fig. 2, and Supplementary Tables 6 and 7). All 52 variants were rare (population allele frequency <0.2%), and 23 of the 52 variants had been reported as clinically significant [21] in hereditary blood disorders (eight *MPL* variants and one *SH2B3* variant) or cancer (11 *ATM* variants and one variant each in *MRE11, NBN*, and *FH*). All 28 *MPL* variants were consistently removed from the genomes of expanded clones by CNN-LOH mutations (244 of 244 cases, binomial *P* =7.1×10^−74^), consistent with the idea that the inherited alleles (with reduced MPL function) have a hypo-proliferative effect that is rescued by acquired CNN-LOH [10]. The 24 variants at the other six loci were systematically duplicated by CNN-LOH mutations (233 of 239 cases, binomial *P* =5.6×10^−61^), consistent with pro-proliferative effects of reduced ATM, MRE11, NBN, SH2B3, TM2D3, and FH function (Table 1, Fig. 2 and Supplementary Table 6). Identity-by-descent (IBD)-sharing analyses of individuals with 1p CNN-LOH mutations spanning *MPL* and individuals with 11q CNN-LOH mutations spanning *ATM* indicated that while the risk variants we identified (Table 1 and Supplementary Table 6) are the primary drivers of heritable CH risk at these loci, the full allelic series likely include many more risk variants (Supplementary Figures 31 and 32).

To detect additional potential risk variants and to estimate the fraction of CNN-LOH clones attributable to inherited protein-altering variants at each locus, we examined exome sequence data available for 49,960 of the UK Biobank participants [22]. Among 271 exome-sequenced individuals with unexplained mosaic CNN-LOH events spanning the seven loci (i.e., not carrying any of the 52 variants already identified), 22 individuals carried 21 distinct ultra-rare coding or splice variants that altered the encoded proteins (vs. 1.28 individuals expected by chance, *P* =2.8×10^−20^; “ultra-rare” refers to population allele frequency less than 0.0001; Methods and Supplementary Tables 8–10). Collectively, *MPL* variants identified by these association and burden analyses were present in 39 of 71 exome-sequenced individuals with 1p CNN-LOH events spanning *MPL* (vs. 0.5 expected), indicating that ∼54% of acquired 1p CNN-LOH events are driven by inherited coding or splice variants at *MPL* (Supplementary Table 10). Similarly, inherited variants at *ATM, NBN, SH2B3*, and *TM2D3* appeared to drive ∼17–33% of CNN-LOH events spanning these loci (Supplementary Table 10). Altogether we estimate that about 5% of clones with CNN-LOH arose from one of these seven monogenic inherited vulnerabilities.

Common inherited variants at five loci conferred more modest CH risk. Common variants at *TCL1A* and *DLK1* on 14q associated with acquired 14q CNN-LOH mutations (Supplementary Table 11 and Supplementary Note), whereas common variants at *TERC, SP140*, and the previously-implicated *TERT* locus [8] broadly increased the risk of CH involving any autosomal mCA (Supplementary Table 12 and Supplementary Note). Notably, *TERC* and *TERT* both encode proteins with key roles in the maintenance and elongation of telomeres.

Some CNN-LOH events provided “second hits” to acquired point mutations. At the frequently-mutated *DNMT3A, TET2*, and *JAK2* loci [4, 5], ∼24–60% of CNN-LOH mutations appeared to provide second hits to somatic point mutations detectable from exome sequencing reads (Supplementary Fig. 33 and Supplementary Table 10; additional CNN-LOH events spanning these loci might be explained by point mutations present at lower cell fractions we could not detect; Methods). Among 33 exome-sequenced individuals with 9p CNN-LOH events, 20 individuals had at least one read suggesting *JAK2* V617F mutation; conversely, 18 of 46 individuals with *JAK2* V617F calls had a detectable mCA on 9p (15 CNN-LOH events and three chromosome 9 duplications). Together, the putative “second-hit” clones at these loci accounted for about 0.3% of all detected CNN-LOH clones.

The great majority of the 17,111 hematopoietic clones we observed in UKB still had unknown causes; most clones had CNN-LOH mutations, which were numerous on every chromosome arm (Fig. 1). Recent work in human and agricultural genetics has revealed that many phenotypes are shaped by polygenic effects from alleles of modest effect at hundreds of loci on all chromosomes [23–25]. We wondered whether polygenic inheritance—even along a single chromosome arm—might be sufficiently potent to drive clonal advantage and evolution among an individual’s cells.

We hypothesized that inherited haplotypes of many common alleles along a chromosome arm can themselves be instruments for clonal selection (Fig. 3a). To evaluate this possibility, we tested whether the haplotypes duplicated and deleted by likely-CNN-LOH mutations (Methods) tended to differ systematically in “proliferative potential” as estimated from combinations of many inherited alleles. We estimated this proliferative potential by building polygenic statistical models [26] for blood-cell abundance traits (using data on blood-cell counts from UKB participants) and for clonal Y chromosome loss, a frequent marker of hematopoietic clones [27, 28]. Based on these models, we estimated “proliferative polygenic risk scores” (PPRS) for the combinations of common alleles along the haplotypes gained and lost by CNN-LOH mutations in expanded clones (Methods).

**Figure 3.**
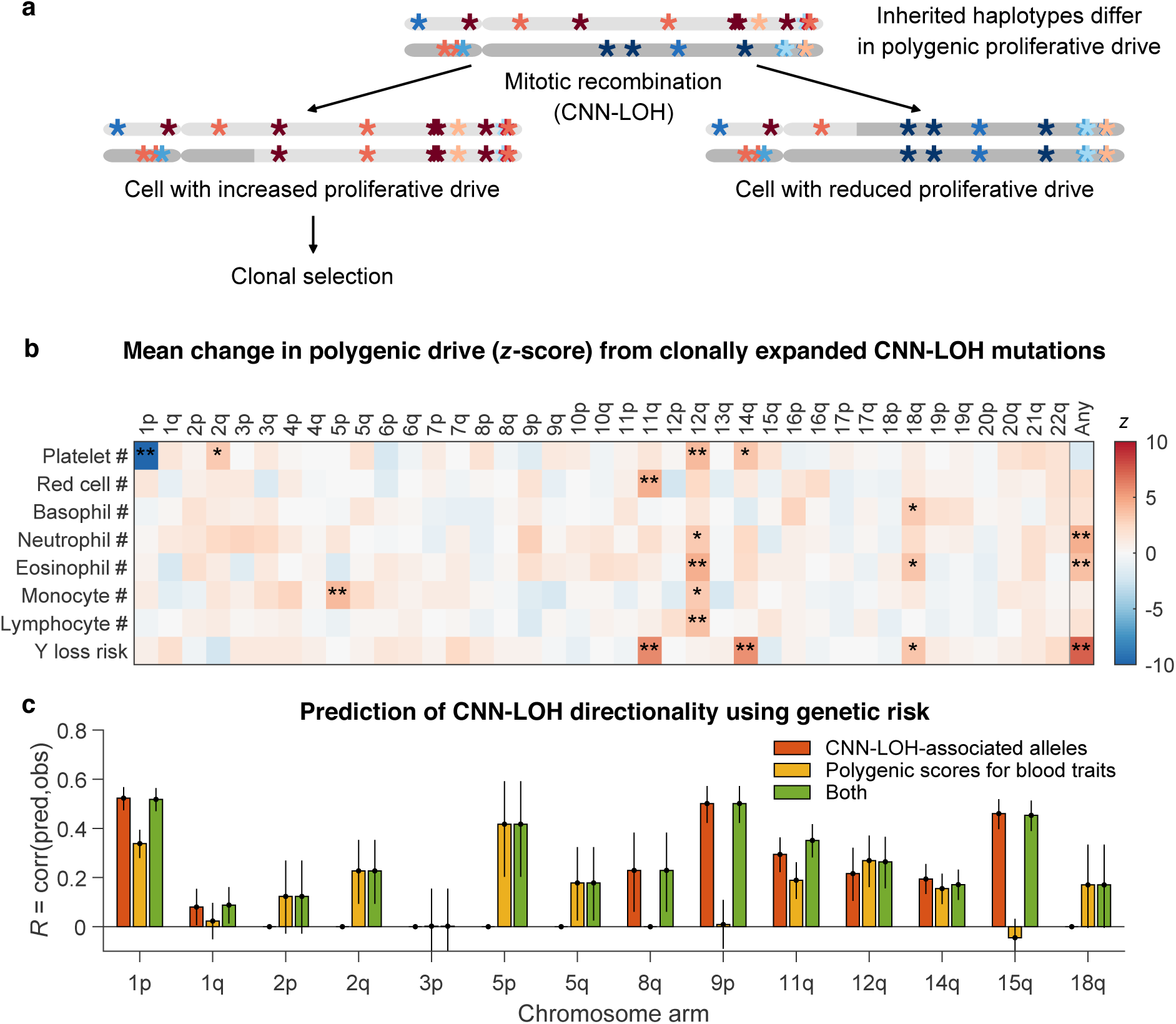
Polygenic and monogenic influences on clonal proliferation of cells with CNN-LOH mutations. (**a**) Two cellular outcomes of a CNN-LOH mutation (mitotic recombination) involving homologous chromosome arms that bear inherited alleles with differing proliferative potentials. In one cell, the CNN-LOH mutation has duplicated the chromosomal arm that has alleles that more strongly promote proliferation; proliferative polygenic drive increases, potentially resulting in clonal selection of the mutant cell. By contrast, the cell with the complementary CNN-LOH mutation may have reduced tendency to proliferate. (**b**) CNN-LOH mutations in expanded clones broadly increase polygenic risk scores for increased blood-cell counts and risk of mosaic Y chromosome loss (a marker for clonal hematopoiesis [27, 28]). The heatmap displays changes in polygenic scores for each trait, averaged across all ascertained (expanded) CNN-LOH mutations observed on each chromosome arm (color bar, *z*-score; *, significant at FDR<0.05; *,*,Bonferroni-corrected *P*<0.05). (**c**) Prediction of the direction of CNN-LOH mutations (in expanded clones) from inherited alleles on the affected chromosome arms. Prediction accuracy (the correlation between predicted and observed CNN-LOH direction) is plotted for predictions made using: only CNN-LOH-associated alleles (Table 1 and Supplementary Table 6) (red); polygenic score differentials on affected chromosomal segments (orange); or both sources of information (green). Error bars, 95% CIs. Results are plotted for 14 chromosome arms for which at least one predictor was available. Numeric data are provided in Supplementary Tables 13 and 15. Analyses of polygenic scores for control traits such as height and BMI are provided in Supplementary Table 14.

CNN-LOH mutations in hematopoietic clones tended to have caused chromosomal segments with higher PPRS to replace homologous (allelic) counterparts with lower PPRS (Fig. 3b). Averaging across all autosomal CNN-LOH events, the allelic substitutions produced by CNN-LOH mutations significantly increased PPRS for clonality with Y chromosome loss (*P* =1.2×10^−13^) and also tended to increase PPRS for the individual blood-cell proliferation traits (most significant: neutrophil counts, *P* =7.5×10^−6^; eosinophil counts, *P* =1.4×10^−4^). This effect was observed throughout the genome: 14 distinct combinations of chromosome arms and cell-abundance traits exhibited significant upward shifts in PPRS (at an FDR of 0.05), and 209 of all 312 combinations exhibited a positive mean increase (*P* =2.0×10^−9^, sign test; Fig. 3b and Supplementary Table 13). These effects were specific to polygenic risk for blood cell traits: clonally selected CNN-LOH mutations did not tend to affect polygenic scores for control traits such as height and BMI (Supplementary Table 14).

These results raised the intriguing possibility that the direction of mosaic CNN-LOH mutations— i.e., which haplotype is to be deleted, and which is to be duplicated, in clonally selected cells—can be predicted from inherited variation. To test this idea, we performed cross-validated prediction using logistic regression on either (i) the CNN-LOH-associated alleles we had found (Table 1); polygenic score (PPRS) differentials on chromosomal segments affected by CNN-LOH; or both CNN-LOH-associated alleles and PPRS differentials (Methods). Polygenic scores and specific inherited CNN-LOH-associated alleles each helped predict CNN-LOH directions; combining both sources of information yielded the most predictive information, reaching significance (FDR<0.05) for 12 of 14 chromosome arms tested (Fig. 3c and Supplementary Table 15; we tested the 14 arms for which the prediction algorithm nominated at least one predictor for testing in a nonoverlapping data set; Methods). The directions of CNN-LOH mutations were correctly predicted for 59% (*P* =4.8×10^−44^) of 5,582 CNN-LOH events on these 14 arms (range 50-–70%). Stronger inherited imbalances correlated with greater predictability: upon restricting to events involving larger imbalances in PPRS (top quintile), prediction accuracy increased to 72% (*P* =1.1×10^−82^).

Clonal hematopoiesis substantially increases risk of adverse health outcomes, including blood cancers, cardiovascular disease, and mortality [2–5, 9, 29]. Previously, we observed that mosaic chromosomal alterations on different chromosomes make a range of contributions to blood cancer risk [10]. The size of the full UK Biobank data set allowed us to further appreciate the extent to which different mCAs associate with distinct health outcomes (Methods). Thirteen specific mCAs significantly associated (FDR<0.05) with subsequent hematological cancer diagnoses during 4–9 years of follow-up, with +12, 13q–, and 14q– events conferring >100-fold higher risk of chronic lymphocytic leukemia, and *JAK2*-related 9p CNN-LOH events conferring 260-fold (89– 631-fold) higher risk of myeloproliferative neoplasms (Fig. 4a and Supplementary Table 16). The far-more-common CNN-LOH events on other chromosome arms also significantly increased blood cancer risk (aggregate hazard ratio=2.84 (2.14–3.78), excluding the 9p events). (We corrected these analyses for age and sex and restricted to individuals with normal blood counts at assessment, no previous cancer diagnoses, and no cancer diagnoses within one year of assessment.) We did not find a significant increase in cardiovascular risk among individuals with most categories of clones—with the notable exception of *JAK2*-related 9p CNN-LOH events (Fig. 4b and Supplementary Table 17)—suggesting that the relationship between clonal hematopoiesis and cardiovascular disease [5, 29] arises from clones that harbor specific mutations.

**Figure 4.**
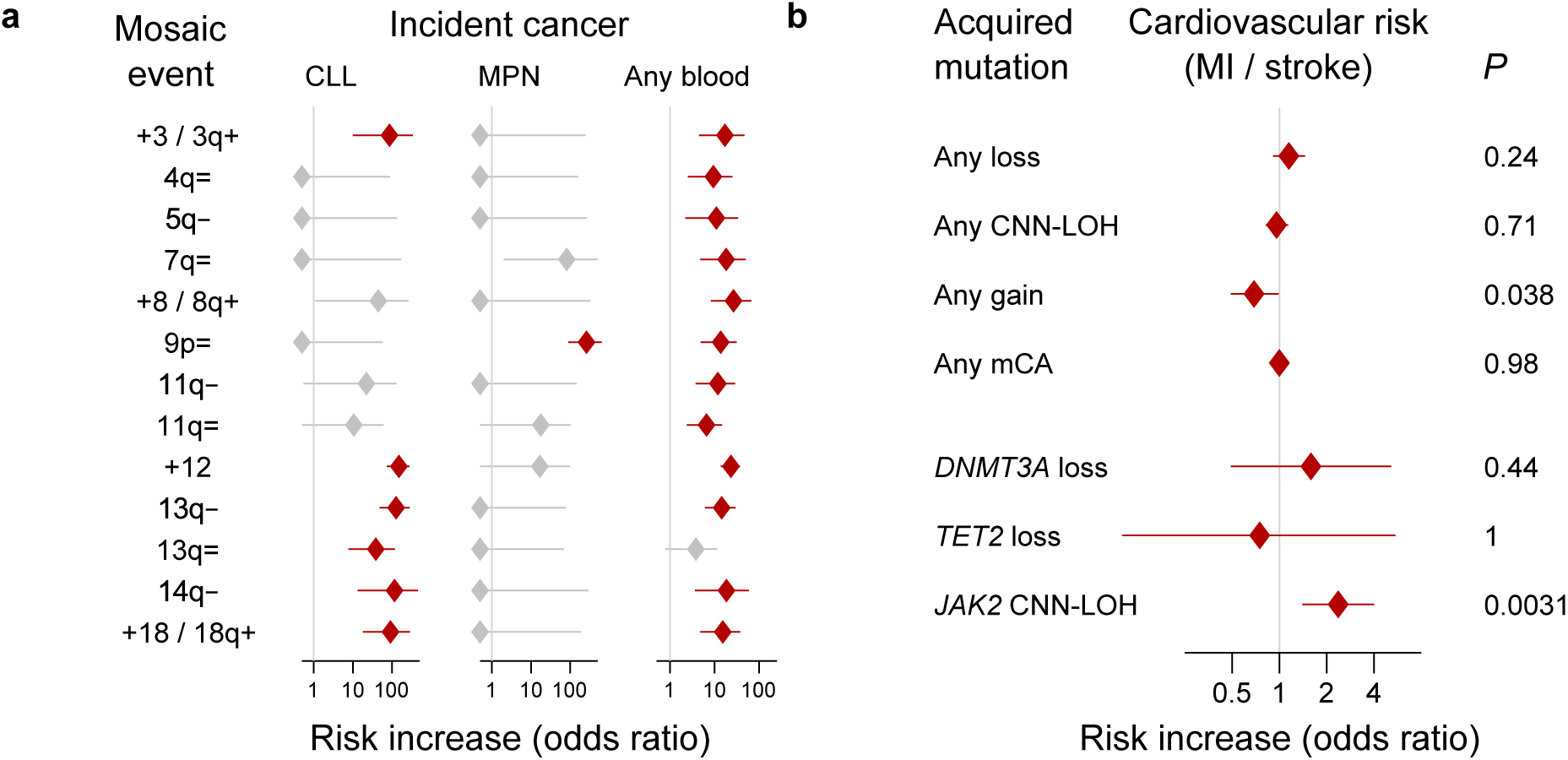
Associations of mCAs with incident cancers and cardiovascular disease. (**a**) Clones with specific mCAs confer increased risk of incident blood cancers diagnosed >1 year after DNA collection in individuals with normal blood counts at assessment (Cochran-Mantel-Haenszel test adjusting for age and sex; error bars, 95% CIs). (**b**) Loss, CNN-LOH, and gain events (on any autosome) do not broadly increase risk for incident myocardial infarction or stroke, but CNN-LOH events on 9p (containing *JAK2*) do increase cardiovascular risk [29] (Fisher’s exact test on cases and controls matched for assessment year, age, sex, smoking, hypertension, BMI, and type 2 diabetes status; error bars, 95% CIs). Statistical tests are detailed in Methods. Numeric data are provided in Supplementary Tables 16 and 17.

These results illuminate the clonal advantages conferred by CNN-LOH, the common substitution of one chromosome arm for its homologous counterpart, which was present in most of the clones ascertained by mCAs (Fig. 1). Although the first-order gene-dosage effects of deletions and duplications are clear [2, 3, 6, 7, 30], clonal expansions of copy-number neutral mutations are both more common (Fig. 1) and more mysterious: the substitution of one chromosome arm for its inherited homolog does not modify gene dosage, so why would a cell that has undergone such a mutation gain a proliferative advantage? Our results, obtained from many genomic loci, point to a single core principle: clonally expanded CNN-LOH events routinely replace inherited chromosomal segments with homologous segments that more strongly promote proliferation. Examples of potent CNN-LOH events have previously been observed in disease studies at a few loci where CNN-LOH events provide second hits to acquired mutations [31], disrupt imprinting [32], or revert pathogenic mutations in rare monogenic disorders of the skin and blood [33, 34]. We recently observed that CNN-LOH mutations can also lead to clonal selection in healthy blood by modifying allelic dosage of inherited rare variants at three loci [10]. The analyses described here demonstrate that this proliferative mechanism is in fact at work throughout the genome: we identified six more loci (*FH, NBN, MRE11, SH2B3, TCL1A*, and *DLK1*) at which CNN-LOH mutations gain advantage from at least 50 inherited alleles, and we observed a pervasive polygenic effect driven by combinations of inherited alleles along chromosome arms. Our finding that the direction of 5,582 CNN-LOH mutations (across 14 chromosome arms) could be predicted with 59% accuracy—based only on the alleles inherited on each arm—implies that a substantial fraction of clonal expansions with CNN-LOH (at least 59 – 41 = 18%) are influenced by inherited alleles that cause maternal and paternal haplotypes to differ in their tendency to promote proliferation. Furthermore, this underestimates the strength and prevalence of polygenic selective pressure; as polygenic risk scores are informed by larger samples and lower-frequency alleles, their predictive accuracy tends to greatly increase [24, 25, 35].

We were initially surprised that even a modest fraction of an individual’s polygenic risk— arising from a single chromosome arm—could create substantial clonal advantage. We believe that this results from an important aspect of clonal evolution: mutated cells compete with nearly isogenic cells in a common, shared environment. Estimates of the effects of common alleles and polygenic risk—which are usually made in the context of diverse genetic backgrounds and abundant environmental variation—are likely to underestimate the potential of such alleles to become instruments for clonal selection.

Because human populations harbor abundant heterozygosity, and mitotic recombination events occur frequently over the course of an individual’s lifetime [10, 33], imbalances in the proliferative potential of the homologous chromosome arms inherited from one’s two parents provide a context in which clonal selection is almost inevitable. Managing this dynamic presents challenges for cytopoiesis throughout the lifespan in any genetically diverse species.

## Acknowledgments

We thank S. Bakhoum, S. Raychaudhuri, M. Sherman, S. Elledge, and C. Terao for helpful discussions. This research was conducted using the UK Biobank Resource under Application #19808. P.-R.L. was supported by US NIH grant DP2 ES030554, a Burroughs Wellcome Fund Career Award at the Scientific Interfaces, the Next Generation Fund at the Broad Institute of MIT and Harvard, and a Glenn Foundation for Medical Research and AFAR Grants for Junior Faculty award. G.G. and S.A.M. were supported by US NIH grant R01 HG006855. G.G. was supported by US Department of Defense Breast Cancer Research Break-through Award W81XWH-16-1-0316. Computational analyses were performed on the O2 High Performance Compute Cluster, supported by the Research Computing Group, at Harvard Medical School (http://rc.hms.harvard.edu), and on the Genetic Cluster Computer (http://www.geneticcluster.org) hosted by SURFsara and financially supported by the Netherlands Scientific Organization (NWO 480-05-003 PI: Posthuma) along with a supplement from the Dutch Brain Foundation and the VU University Amsterdam.

## Author Contributions

All authors designed the study. P.-R.L. performed computational analyses. All authors wrote the paper.

## Author Information

The authors declare competing interests: patent application PCT/WO2019/ 079493 has been filed on the mCA detection method used in this work. Correspondence should be addressed to P.-R.L., G.G., or S.A.M.

## Methods

### UK Biobank cohort and genotyping

The UK Biobank is a very large prospective study of individuals aged 40–70 years at assessment [11]. Participants attended assessment centers between 2006 and 2010, where they contributed blood samples for genotyping and blood analysis and answered questionnaires about medical history and environmental exposures. In the years since assessment, health outcome data for these individuals (e.g., diagnoses of cancer and cardiovascular disease) have been accruing via UK national registries and hospital records managed by the NHS.

We analyzed genetic data from the full UK Biobank cohort, which consists of 488,377 individuals genotyped on the Affymetrix UK BiLEVE and UK Biobank Axiom arrays. The BiLEVE and Biobank arrays have >95% overlap and contain a total of 784,256 unique autosomal variants; 49,950 individuals were genotyped on the BiLEVE array [36] and the remaining individuals on the Biobank array. We restricted our analyses to 487,409 individuals passing previous genotyping QC and previously imputed to ∼93 million autosomal variants [12]; we re-phased these individuals using Eagle2 [14] to improve phasing accuracy and imputed them to the union of the BiLEVE and Biobank arrays using Minimac3 [37] (Supplementary Note). We further removed 427 individuals with low genotyping quality (B-allele frequency s.d.>0.11 at heterozygous sites), 4,111 individuals with evidence of possible sample contamination (Supplementary Note), and 82 individuals who had withdrawn consent, leaving 482,789 individuals for analysis. We performed data processing using `plink` [38].

We additionally analyzed exome sequencing data available for 49,960 individuals [22]. To extend our rare variant association analyses to include variants identified in exome-sequenced individuals, we phased these individuals using Eagle2 and imputed into the full cohort using Minimac4 (Supplementary Note).

### Detection of mCAs using genotyping intensities and long-range haplotype phase

As in our previous work [10], we detected mCAs in genotyping intensity data from blood DNA samples using an approach that leverages the chromosome-scale accuracy of statistical phasing in the UK Biobank cohort [13, 14] (Supplementary Note). In brief, our approach harnesses long-range phase information to search for local imbalances between maternal and paternal allelic fractions in a cell population, enabling considerable gains in sensitivity for detection of large events at low cell fractions [10]. As before, we applied our approach to genotyping intensities that we transformed to log_2_ R ratio (LRR) and B-allele frequency (BAF) values [39] (which measure total and relative allelic intensities) after affine-normalization and GC wave-correction [2, 10, 40].

In analyzing the full cohort, we made two minor modifications to our original approach. First, we halved the switch error rate parameter of our hidden Markov model (HMM) for BAF deviations, reflecting improved phasing accuracy in the full cohort. Second, we perfomed a few additional QC steps on the event calls to filter potential technical artifacts that became visible in the full data set; these filters affected <1% of the call set (Supplementary Note).

Our detection procedure produced a final call set of 19,632 autosomal mCAs at a nominal FDR of 0.05 (based on our phase randomization approach to estimate statistic significance) [10]. We verified that our FDR was well-controlled using an independent FDR estimation procedure based on the age distribution of event carriers [10]; this approach produced a concordant FDR estimate of 6.6% (4.5–8.6%) (Supplementary Fig. 34 and Supplementary Note). As before, we observed that lower-confidence events tended to have uncertain copy number (because our power to detect allelic imbalances exceeds our power to distinguish CNN-LOH from copy-number alterations) and less precise event boundaries [10]; we provide information on the uncertainty of each event call in Supplementary Data.

### Identifying variants associated with CNN-LOH mutations in *cis*

We performed two types of association tests to identify inherited variants that influence mosaic CNN-LOH mutations in *cis*. First, for each variant, we performed a Fisher test for association with a case-control phenotype specific to that variant: we considered samples to be cases if they carried a likely CNN-LOH event containing the variant or within 4Mb (to allow for uncertainty in event boundaries). We considered an event to be a likely CNN-LOH event if it either (i) was called as a CNN-LOH event or (ii) had undetermined copy number, extended to a telomere, and had *|*LRR*|*<0.02. We performed this test on all typed and imputed variants and applied a genome-wide significance threshold of 5×10^−8^ for coding variants and 1×10^−9^ for all other variants.

Second, we searched for variants for which CNN-LOH mutations in individuals heterozygous for the variant tended to preferentially duplicate one allele and remove the other allele from the genome. For each variant, we examined heterozygous individuals with a likely CNN-LOH eventm overlapping the variant, and then performed a binomial test to check whether the CNN-LOH direction tended to favor one allele versus the other. We restricted the binomial test to individuals in which the variant was confidently phased relative to the mosaic event, i.e., there was no disagreement in five random resamples from the HMM used to call the event).

Given that the two association tests described above are independent, the second test provided a means of validating associations identified by the first test, as any spurious associations from the first test would have no correlation with CNN-LOH direction, whereas variants truly associated with CNN-LOH mutations in *cis* typically have strong associations with CNN-LOH direction (Table 1). We also performed a combined test to identify common variants that did not reach genome-wide significance in the first test alone (which was underpowered for common variants due to small case counts) but reached significance using both tests together (Fisher’s combined *P*<1×10^−8^).

We restricted our association analyses to 455,009 individuals who reported European ancestry. Among these individuals, 96,590 pairs had previously been identified to be third-degree or closer relatives [12,41]. For each chromosome, we pruned the samples to an unrelated subset by removing one individual from each related pair, preferentially keeping (i) individuals with a likely CNN-LOH on the chromosome and (ii) older controls. This pruning decreased total sample sizes to slightly less than 380,000 individuals (Supplementary Table 18). We verified that filtering on ancestry and relatedness in this way produced well-calibrated association test statistics (Supplementary Fig. 35).

### Fine-mapping loci associated with CNN-LOH mutations in *cis*

Given that our association analyses identified rare, large-effect coding variants in seven genes (*FH, NBN, MRE11, SH2B3, MPL, ATM*, and *TM2D3*), we undertook fine-mapping analyses at these loci to uncover additional coding or splice variants in these genes likely to be objects of clonal selection (upon modification of allelic load via CNN-LOH mutation). We tested variants in these genes in three categories: (i) missense variants with CADD v1.3 score >20 (ref. [42]); (ii) predicted LoF variants (i.e., stop gained, frameshift, splice acceptor, or splice donor sites in any transcript annotated by VEP [20]); and (iii) likely pathogenic variants (according to ClinVar [21], downloaded Mar 25, 2019). We restricted these analyses to rare variants with MAF between 5×10^−6^ and 0.01. For directly genotyped variants, we required missingness <0.01; for imputed variants, we required INFO>0.2 (for variants imputed by UK Biobank using IMPUTE4 [12]) or Minimac *R*^2^>0.4 (for variants we imputed; Supplementary Note). In addition to variants available from genotyping and imputation, we also tested two structural variants: a 454bp deletion that we discovered in *MPL* (Supplementary Figures 28–30 and Supplementary Note) and a ∼70kb deletion of *TM2D3* that we previously identified [10]. In total, 616 variants across the seven loci satisfied these criteria.

Of these 616 variants, 38 variants reached Bonferroni significance (*P*<8.1×10^−5^; Table 1) and 52 variants reached FDR<0.05 significance (assessed per gene; Supplementary Table 6). We determined that all 52 FDR-significant variants were likely to causally drive independent associations with CNN-LOH events in *cis*, based on the following lines of evidence. First, CNN-LOH events acted on all 52 variants in the expected direction (consistently removing rare variants in *MPL* and duplicating rare variants in the other six genes; Supplementary Table 6); in contrast, variants associated by chance would have random phase relative to CNN-LOH events. Second, none of the 52 variants tagged other nearby variants with stronger associations. On the contrary, variants in linkage disequilibrium with the 52 variants had weaker associations explained by tagging of the 52 variants (Fig. 2); e.g., the variants near *MPL* and *ATM* that we previously reported [10] each tagged one of the 52 variants (Supplementary Table 7). Third, none of the 52 variants tagged each other. The association signals at the 52 variants were driven by almost entirely non-overlapping sets of carriers who also had CNN-LOH events in *cis*; the only overlap occurred between 11q CNN-LOH individuals carrying the rs587779872 *ATM* missense variant (6 carriers with 11q CNN-LOH) and the rs786204751 *ATM* stop gain variant (2 carriers with 11q CNN-LOH, both also carrying rs587779872; Supplementary Fig. 32). The rs587779872 association remained significant in non-carriers of rs786204751, while the rs786204751 stop gain mutation nullified the effect of the rs587779872 missense mutation (occurring later in *ATM*), leading us to conclude that these associations were likely to be independent.

### Burden analyses to detect ultra-rare variants targeted by CNN-LOH events

To identify CNN-LOH events potentially explained by variants too rare to reach significance in single-variant association analyses, we analyzed variant calls from exome sequencing of 49,960 UK Biobank participants [22] for a burden of ultra-rare coding and splice variants in individuals with CNN-LOH events. Because these variant calls potentially contained a small fraction of somatic variants that had risen to cell fractions higher than ∼20%, we included *DNMT3A, TET2*, and *JAK2* in these analyses in addition to the seven genes at which we found inherited variants influencing CH. Beyond being frequently mutated in CH [4, 5], *DNMT3A, TET2*, and *JAK2* are also frequently overlapped by CNN-LOH events (Fig. 1), suggesting that some CNN-LOH events act on previously-acquired point mutations in these genes.

As in our fine-mapping analyses, we considered variants annotated as (i) missense with CADD score >20, (ii) predicted LoF, or (iii) likely pathogenic in ClinVar. We restricted to ultra-rare variants (MAF<1×10^−4^), with the exception of *JAK2* V617F, which was called in 46 exome-sequenced individuals (MAF=4.6×10^−4^). (For *JAK2* and *ATM*, we used exome variant calls generated by UK Biobank using the “functionally equivalent” (FE) pipeline [43], which we found provided slightly better power at these loci; for all other analyses, we used variant calls from Regeneron’s Seal Point Balinese (SPB) pipeline [22].) For each gene, we examined individuals with CNN-LOH events spanning the gene (not already explained by any of the 52 variants identified in our association analyses) and tabulated the number of such individuals who carried any of the rare variants under consideration (Supplementary Table 9). We then computed a burden *P*-value using a one-sided binomial test comparing the observed count to expectation (based on variant frequencies among 46,633 exome-sequenced individuals who reported European ancestry).

For each variant call potentially targeted by a CNN-LOH event, we further examined allelic read depths from the exome sequencing data to assess whether the variant was likely to be of inherited or acquired origin. While read depths were generally insufficient to make a confident assessment on a per-variant level (and making this determination is complicated by mapping bias toward the reference allele [4]), the allelic depths broadly indicated that all or most variants implicated at our seven inherited risk loci were indeed inherited, while all or most variants implicated at *DNMT3A, TET2*, and *JAK2* had been acquired somatically (Supplementary Fig. 33).

### GWAS for *trans* associations with any autosomal mCA

We tested common variants for *trans* associations with the presence of any detectable autosomal mCA. We computed association test statistics using BOLT-LMM [26, 44] on 452,469 individuals (of which 16,366 were cases) who reported European ancestry and had imputation data available on autosomes and the X chromosome [12]. We included 20 principal components, age, age squared, sex, smoking status, genotyping array, and assessment center as covariates in the linear mixed model.

### Polygenic scores for blood cell traits

We analyzed 29 blood count traits: counts and percentages of basophils, eosinophils, lymphocytes, monocytes, neutrophils, platelets, red cells, reticulocytes, and high light scatter reticulocytes; white cell count, platelet and red cell distribution widths, immature reticulocyte fraction, hemoglobin concentration, mean corpuscular hemoglobin (MCH), MCH concentration, mean corpuscular volume, mean platelet volume, mean reticulocyte volume, and mean sphered cell volume. (These traits constituted all available blood count traits except nucleated red blood cell indices, which were mostly zero.) We performed basic QC and normalization on these traits using the following steps: (i) remove outliers (>7 times farther from median than nearest quartile); (ii) stratify into males, pre-menopausal females, and post-menopausal females; (iii) within each stratum: (a) inverse-normal transform; (b) regress out age, age^2^, height, height^2^, BMI, BMI^2^, ethnic group, alcohol use, and smoking status; (c) inverse-normal transform again.

We computed polygenic score coefficients (i.e., “betas” in a linear predictor) for the traits listed above using the --`predBetasFile` option of BOLT-LMM [26, 44], which estimates polygenic score coefficients using a Bayesian linear mixed model that accounts for linkage disequilbrium among variants. We computed coefficients for 709,999 autosomal and X chromosome variants in the intersection of the Biobank and BiLEVE arrays that passed QC filters (allele frequency deviation <0.02 between the arrays, missingness <0.05, failed QC in at most one genotyping batch [12]). For each blood count phenotype, we restricted the sample set to individuals of selfreported European ancestry with non-missing phenotype (437,009–445,438 individuals depending on the phenotype). We ran BOLT-LMM using the same set of covariates we used in our *trans* GWAS. We computed polygenic risk coefficients for Y loss in blood cells using an analogous analysis restricted to males [28].

### Polygenic score differentials for CNN-LOH events

The polygenic score coefficients we computed for blood cell traits allowed us to estimate the extent to which CNN-LOH mutations modified the genetic components of these traits. For each CNN-LOH mutation, we computed the difference in polygenic score carried by the haplotype that was duplicated versus the haplotype that was removed. (This quantity is equal to the difference between the polygenic load of the mutant CNN-LOH genome versus the original genome.) We determined which haplotype was duplicated and which was deleted using our hidden Markov model of phased BAF deviations [10], averaging across five posterior samples from the HMM. To identify chromosome arms in which CNN-LOH events tended to increase polygenic load for specific blood cell traits, we averaged polygenic score differentials across all CNN-LOH events on each arm and computed means and *z*-scores (indepedently for each blood cell trait; Fig. 3b and Supplementary Table 13). To maximize power, we included all “likely-CNN-LOH” events in these analyses (i.e., events called as CNN-LOH as well as events with undetermined copy number that extended to a telomere and had *|*LRR*|*<0.02, as in our *cis* association analyses), comprising a total of 11,667 likely-CNN-LOH events.

### Prediction of CNN-LOH directions using CNN-LOH-associated alleles and polygenic scores

To assess the extent to which the direction of a CNN-LOH event (i.e., which affected haplotype is duplicated and which one is deleted) can be predicted based on the alleles inherited on each haplotype, we fit logistic models on the CNN-LOH events on each chromosome arm using 10-fold cross-validation. For each fold, we performed logistic regression using stepwise forward selection on three possible sets of predictors: (i) a single variable containing the difference in the number of CNN-LOH-associated alleles (Table 1 and Supplementary Table 6) carried by the two affected haplotypes; (ii) 30 variables containing the polygenic score differentials (for the 29 blood count indices and the Y loss trait) between the two affected haplotypes; and (iii) all 31 variables together. We started forward selection using the “number of CNN-LOH-associated alleles” variable in analyses (i) and (iii) and an empty set of variables in analysis (ii). We stopped forward selection when model improvement was no longer significant at a 0.01 level. We restricted our prediction analyses to chromosome arms for which at least one variable was selected (on average across folds).

For each chromosome arm, we merged prediction results across the 10 held-out folds and then assessed accuracy in two ways. First, we computed the Pearson correlation (*R*) between observed and predicted CNN-LOH directions (using continuous-valued prediction probabilities from logistic regression). Second, we computed raw prediction accuracy (using binary, hard-called predictions). As in our analyses of polygenic score differentials, we included all likely-CNN-LOH events (as defined above) to maximize power in these analyses.

### Enrichment of mCA types in specific blood lineages

To identify classes of mCAs linked to different blood cell types [10], we first classified mCAs based on chromosomal location and copy number. For each autosome, we defined five disjoint categories of mCAs that comprised the majority of detected events: loss on p-arm, loss on q-arm, CNN-LOH on p-arm, CNN-LOH on q-arm, and gain. We subdivided loss and CNN-LOH events by arm but did not subdivide gain events because most gain events are whole-chromosome trisomies (Fig. 1). (We excluded the chr17 gain category because nearly all of these events arise from i(17q) isochromosomes already counted as 17p– events; Supplementary Fig. 17.)

For each mCA type, we computed enrichment among individuals with anomalous (top 1%) values of each of 14 normalized blood indices (counts and percentages of lymphocytes, basophils, monocytes, neutrophils, red cells, and platelets, as well as distribution widths of red cells and platelets) using Fisher’s exact test (two-sided; *P*-values reported throughout this manuscript are from two-sided statistical tests unless explicitly stated otherwise). We restricted these analyses to individuals who reported European ancestry, and reported significant enrichments passing an FDR threshold of 0.05 (Supplementary Fig. 27 and Supplementary Table 5).

### UK Biobank cancer phenotypes

We analyzed UK cancer registry data provided by UK Biobank for 81,401 individuals in our sample set who had one or more prevalent or incident cancer diagnoses. Cancer registry data included date of diagnosis and ICD-O-3 histology and behavior codes, which we used to identify individuals with diagnoses of CLL, MPN, or any blood cancer [45, 46]. Because our focus was on the prognostic power of mCAs to predict diagnoses of incident cancers >1 year after DNA collection, we excluded all individuals with cancers reported prior to this time (either from cancer registry data or self-report of prevalent cancers). We also restricted our attention to the first diagnosis of cancer in each individual, and censored diagnoses after September 30, 2014, as suggested by UK Biobank (resulting in a median follow-up time of 5.7 years, s.d. 0.8 years, range 4–9 years). Finally, we restricted analyses to individuals who reported European ancestry. These exclusions reduced the total counts of incident cases to 199 (CLL), 138 (MPN), and 1,383 (any blood cancer). In our primary analyses, we further eliminated individuals with any evidence of potential undiagnosed blood cancer based on anomalous blood counts (lymphocyte count outside the normal range of 1–3.5×10^9^/L, red cell count >6.1×10^12^/L for males or >5.4×10^12^/L for females, platelet count >450×10^9^/L, red cell distribution width >15%), leaving incident case counts of 107 (CLL), 67 (MPN), and 1,055 (any blood cancer).

### Estimation of cancer risk conferred by mCAs

To identify classes of mCAs associated with incident cancer diagnoses, we classified mCAs based on chromosomal location and copy number as described above. We then restricted our attention to the 78 classes with at least 30 carriers (to reduce our multiple hypothesis burden, given that we would be underpowered to detect associations with the rarer events). For each mCA class, we considered a sample to be a case if it contained only the mCA or if the mCA had highest cell fraction among all mCAs detected in the sample (i.e., we did not count carriers of subclonal events as cases). We computed odds ratios and *P*-values for association between mCA classes and incident cancers using Cochran-Mantel-Haenszel (CMH) tests to stratify by sex and by age in six 5-year bins. We used the CMH test to compute odds ratios (for incident cancer any time during follow-up) rather than using a Cox proportional hazards model to compute hazard ratios because both the mCA phenotypes and the incident cancer phenotypes were rare, violating assumptions of normality underlying regression. We reported significant associations passing an FDR threshold of 0.05 (Fig. 4a and Supplementary Table 16).

### UK Biobank cardiovascular disease phenotypes

We analyzed algorithmically-defined cardiovascular events (myocardial infarction and stroke) identified by UK Biobank for 26,873 individuals in our sample set. Events had been identified based on information from baseline questionnaires and/or nurse-led interviews and from linked hospital admission and death registry data sets. We restricted our analyses to individuals with no missing cardiovascular covariates, self-reported European ancestry, and no prevalent cardiovascular disease, and we further pruned to a subset of individuals unrelated at the third degree [12, 41], leaving 360,513 individuals, of which 6,998 had incident cardiovascular events during 5–10 years of follow-up.

### Estimation of cardiovascular risk conferred by mCAs

To increase statistical power and limit the multiple hypothesis testing burden, we grouped all incident cardiovascular events into a single case-control phenotype and tested this phenotype for association with detectable mCAs. We considered mosaicism phenotypes defined by grouping all autosomal mCAs into one phenotype or by grouping mCAs by copy number (loss, CNN-LOH, or gain), and we also examined specific mCAs related to common mosaic point mutations [4, 5, 29]: focal deletions at *DNMT3A*, focal deletions at *TET2*, and CNN-LOH mutations on 9p (which often duplicate a *JAK2* V617F mutation [47–50]) (Fig. 1). For each category of mCAs, we created a subsample of mCA carriers and noncarriers matched on assessment year, age (in 1-year bins), sex, smoking status (current/ever/never), hypertension status, BMI (<25, 25–30, >30), and type 2 diabetes status, selecting carrier-noncarrier ratios to maximize power. We estimated cardiovascular risk conferred by each category of mCAs by performing Fisher’s exact test on the matched sample sets.

## Code availability

Code used to perform the analyses in this study is available from the authors upon request. To facilitate similar analyses of other data sets, a standalone software implementation of the algorithm used to call mCAs (MoChA) is available at https://github.com/freeseek/ mocha.

## Data availability

Mosaic event calls are available in Supplementary Data. Access to the UK Biobank Resource is available by application (http://www.ukbiobank.ac.uk/).

