## Supplementary Information for "Monogenic and polygenic inheritance become instruments for clonal selection"

**4 Common variants influencing mCAs in *cis* 7**

**5 Inherited variants associated with mCAs in *trans* 8**

**6 Shared genetic risk of CLL and mosaic +12 and 13q LOH 9**

**7 Estimation of mortality risk conferred by mCAs 10**

#### References 11

#### Supplementary Figures 16

#### Supplementary Tables 54

### Supplementary Note

### 1 Phasing and imputation of the UK Biobank cohort

#### 1.1 Phasing

The UK Biobank cohort was previously phased using SHAPEIT3 [12, 51]; however, to improve phasing accuracy [13, 14] and to expand the set of phased variants, we rephased the data set using Eagle2 [14], employing a multiple-run voting strategy [52] to optimize accuracy. The SHAPEIT3-phasing performed by UK Biobank included 670,739 autosomal markers present on both the BiLEVE and Biobank arrays that passed the following filters: (a) failed QC in at most 1 genotyping batch, (b) missingness $<$0.05, (c) MAF$<$0.0001 [12]. We performed five runs of phasing using Eagle2 on five distinct marker sets:

1. The same set of autosomal markers previously phased using SHAPEIT3.

2. A subset of 650,084 autosomal markers obtained by further excluding MAF$<$0.001 variants.

3. A separate set of 706,877 autosomal markers present on the Biobank array (but not necessarily the BiLEVE array) passing the following four filters: (i) allele frequency deviation $<$0.02 between the Biobank and BiLEVE arrays (for markers present on both the Biobank and BiLEVE array); (ii) missingness $<$0.15; (iii) either (iii-a) MAF$>$0.001 and passing filters used in SHAPEIT3 phasing, or (iii-b) MAF$<$0.001; (iv) Hardy-Weinberg disequilibrium $P > 10^{-100}$ (according to `plink` [38]).

4. A larger set of 712,138 autosomal markers obtained using the same four filters as above, but relaxing the Hardy-Weinberg threshold to $P > 10^{-200}$.

5. A larger set of 714,468 autosomal markers obtained using the same four filters as above, but relaxing the Hardy-Weinberg threshold to $P > 4.9 \times 10^{-324}$ (the smallest representable double-precision floating point value).

In each phasing run, we ran Eagle2 (v2.3.5) on 105 overlapping chunks of $\sim$10,000 markers (with overlaps of at least 2,000 markers between consecutive chunks on the same autosome); on very large data sets, this partition-ligation approach improves Eagle2's accuracy because Eagle2 conditions on a fixed set of `--Kpbwt` haplotypes per individual within an input region. We set `--Kpbwt` to 100,000 for the first two runs and 80,000 for the remaining three runs, and we used the `--pbwtOnly` option to only use PBWT iterations [14].

We combined phasing results from the five Eagle2 runs (covering a total of 716,197 unique markers) using a voting approach [52], reasoning that phase switch errors incurred during different

Eagle2 runs were likely to be partially independent. Specifically, we scanned through the phased haplotypes in order of genomic position, and at each successive variant, we set the phase of each heterozygous genotype by giving 7 votes to each phasing run containing the variant. These 7 votes were distributed among the most recently processed 7 hets from the run: each het voted according to the relative phase (estimated by Eagle2) between that het and the het currently under consideration. We implemented this approach to improve robustness against short 1–2 SNP "blips" that constitute a large fraction of phase switch errors in large data sets [13]; our goal was to maximize long-range phasing accuracy.

#### 1.2 Benchmarking phasing accuracy using mosaic chromosomal alterations

We benchmarked the long-range phasing accuracy of our Eagle2-phased haplotypes as well as the SHAPEIT3-phased haplotypes provided by UK Biobank using a novel benchmarking method based on mosaic chromosomal alterations. The standard approach to benchmarking phasing accuracy is to analyze a data set that contains trios, comparing gold-standard trio phase to statistical phase estimated on trio children after removing trio parents. However, in this case, we were interested in comparing our accuracy to the accuracy of the existing SHAPEIT3 phasing [12], which had been already been performed on all individuals together (such that trio phase accuracy should be very high [53] and uninformative of phasing accuracy on unrelated individuals).

To overcome this challenge, we instead obtained gold standard phasing information from individuals who carried mosaic chr12 trisomies (the most common whole-chromosome mosaic event). Mosaic events produce allelic imbalances that provide information about phase within genotyping intensity data (i.e., BAF deviations) that is invisible within the genotype calls available to phasing algorithms. To apply this approach, we ran our hidden Markov model-based mCA detection algorithm using either Eagle2-estimated phase or SHAPEIT3-estimated phase, and we compared the numbers of phase switch errors in each data set detected by the HMM. We restricted our attention to 79 individuals with chr12 trisomies with $|\Delta\text{BAF}|$ in the range 0.02–0.05 (high enough that switch errors are easily detectable, but low enough for the mosaicism not to compromise genotype calling accuracy).

Across these benchmark chromosomes, we observed that the SHAPEIT3 phasing achieved a long-range switch error rate of 0.060% (s.e.m. 0.005%), whereas our Eagle2 phasing (voting across five runs) achieved a long-range switch error rate of 0.027% (s.e.m. 0.004%), an improvement of >2x. Applying the same approach to benchmark the individual Eagle2 phasing runs showed that the voting approach achieved a $\sim$20% improvement in accuracy compared to each of the individual runs. We note that these benchmarks ignore small-scale phase switch errors (e.g., 1–2 SNP blips) that will be attributed by the HMM to measurement noise in BAF.

#### 1.3 Imputation

The UK Biobank genetic data was previously imputed to ∼93 million autosomal variants [12] using the Haplotype Reference Consortium (HRC) panel [54] and a merge of the the UK10K and 1000 Genomes Phase 3 reference panels [55]. We augmented this imputed data set by further imputing very rare coding or splice variants contained on the BiLEVE array (used to genotyped 49,950 individuals [36]) but not on the Biobank array (used to genotype the remaining ∼90% of the cohort). To impute these variants to the remainder of the cohort, we first pre-phased the BiLEVE cohort using Eagle2 with `--Kpbwt=20000` using the same partition-ligation scheme we applied to the full cohort (105 overlapping chunks of ∼10,000 autosomal markers; Sec. 1.1). We imputed the BiLEVE-only variants into the full cohort using Minimac3 v2.0.1 [37].

We adopted a similar strategy to impute very rare variants from genic regions captured in exome sequencing of 49,960 UK Biobank participants [22] into the full cohort. We performed phasing and imputation on genotype calls from the SPB exome sequencing pipeline. We dropped singleton variants, phased the remaining exome-sequencing-derived variants together with variants genotyped on the UK Biobank array (using Eagle2 with `--Kpbwt=20000`), and imputed into the full cohort (using Minimac4 v1.0.1, with noncoding variants from the UK Biobank array used as an imputation scaffold).

We also re-imputed chromosomes with detected mCAs in order to obtain phase information at imputed variants, as the imputed data supplied in the UK Biobank imputation v3 release did not contain phase information. We performed this imputation using Minimac3 with the merged UK10K and 1000 Genomes Phase 3 reference panels.

### 2 Detection and calling of an inherited deletion variant in *MPL*

We discovered the inherited 454bp deletion variant associated with 1p CNN-LOH by exploring genotyping intensities of carriers of the rs144279563 tag SNP [10]. We observed an unusual deviation in the total allelic intensities (LRR) for these carriers at four sites typed on the UK BiLEVE chip (used to genotype ∼10% of the cohort): an increase in LRR for a probe at chromosome 1 base position 43,814,653 (hg19) and a decrease in LRR at 43,814,938, 43,814,963, and 43,814,979 (Supplementary Fig. 28). Based on LRR at these four probes, we called 27 likely carriers of the structural variant among the 49,950 UK BiLEVE participants.

Only the first probe (at base position 43,814,653) was included on the Biobank chip (used to genotype the remaining ∼90% of the cohort), so to call the structural variant in the remaining samples, we employed a hybrid approach using both LRR at 43,814,653 and imputation. First, we phased the structural variant in the UK BiLEVE cohort using Eagle2 [14] and imputed it into the remainder of the cohort using Minimac3 [37]. We then re-weighted the imputed allele probabilities for each individual according to the odds of observing the measured LRR at 43,814,653 assuming the individual was a carrier vs. non-carrier of the structural variant. We estimated these odds based on the empirical distribution of LRR among high-confidence carriers (imputed probability >0.99) vs. high-confidence non-carriers (imputed probability <0.01). This re-weighting modified the calls of 16 individuals and produced a final call set of 203 likely carriers of the structural variant in the full cohort.

Upon the release of exome sequencing data for 49,960 UK Biobank participants [22], we examined exome sequencing reads aligning to the region of *MPL* affected by the structural variant in individuals we had predicted to be carriers of the variant. We observed clear read support for a 454bp deletion removing base pairs 43,814,729 through 43,815,182 (spanning *MPL* exon 10): read pairs spanning this region exhibited unusually long insert sizes and clipped alignments (Supplementary Fig. 29), and read depth in exon 10 was unusually low in all 32 predicted carriers of the structural variant who had been exome-sequenced (Supplementary Fig. 30). Interestingly, we found no evidence of a duplication around 43,814,653 in exon 9 (Supplementary Figures 29 and 30, despite our earlier observation that carriers of the structural variant exhibited consistently high genotyping intensities (LRR) at this site (Supplementary Fig. 28). Based on the lack of read support for a duplication in exon 9, we believe the structural variant consists only of the 454bp deletion; the increase in LRR from genotyping at 43,814,653 could be a technical artifact arising from the probe being only 76bp away from the deletion.

### 3 QC of mosaic chromosomal alteration calls

#### 3.1 Identification of samples with possible DNA contamination

In our previous analysis of the UK Biobank $N$=150K interim release, we observed that a small fraction of samples (<1%) exhibited evidence of possible DNA contamination based on apparent short interstitial CNN-LOH calls in specific genomic regions of long-range linkage disequilibrium; we therefore used these likely-artifactual calls to flag samples for exclusion [10]. We applied the same QC approach to the full UK Biobank data set, identifying a total of 4,074 individuals to exclude based on short interstitial CNN-LOH calls in the five regions we previously identified (chr3:∼45Mb, chr6:∼30Mb, chr8:∼45Mb, chr10:∼80Mb, chr17:∼40Mb). As in our previous analysis, we also excluded individuals with three or more interstitial CNN-LOH calls (a mostly-overlapping set of 534 individuals, bringing the total for exclusion to 4,100), and we excluded an additional 7 individuals with three or more calls with high implied switch error rates [10]. Finally, we identified an additional 4 individuals for exclusion based on having eight or more calls all within a very narrow BAF and LRR range (max $|\Delta\text{BAF}|<0.03$, LRR range <0.04), again indicating possible DNA contamination [7]. Together, these criteria resulted in 4,111 exclusions.

#### 3.2 Additional filtering of mosaic event calls

Beyond the sample exclusions described above, we also performed QC on our mosaic event call set to filter calls that were unlikely to be true mosaic events (but did not suggest sample contamination, and hence did not require excluding samples from analysis). As in our previous analysis of the $N$=150K interim data set, our main post-processing step excluded events that might be constitutional (rather than mosaic) duplications [10]. As before, we filtered subchromosomal events of length >10Mb with LRR>0.35 or with LRR>0.2 and $|\Delta\text{BAF}|>0.16$, and we filtered events of length <10Mb with LRR>0.2 or with LRR>0.1 and $|\Delta\text{BAF}|>0.1$. We chose these thresholds conservatively based on visual inspection of LRR and BAF distributions, in which likely constitutional duplications formed well-defined clusters (Supplementary Fig. 36). (Most constitutional duplications were already masked in a pre-processing step involving a separate HMM [10].) We also filtered possible constitutional deletions based on LRR<–0.5 and heterozygosity rate <1/3 the expected rate within called event regions.

Additionally, we filtered 29 event calls with LRR>0.2 and het rate >1.2x expected, LRR>0.1 and het rate >1.5x expected, or LRR>–0.05 and het rate >2x expected. Such event calls with elevated heterozygosity can arise from segmental duplications involving three distinct haploytpes or from genotype calling errors in which rare homozygotes are called as heterozygotes. Finally, we filtered 7 short interstitial events called at the *SNRPN* locus on chr15 between 24–25.5Mb. This locus is imprinted and exhibits differential replication timing between the paternal and the maternal haplotype [56]. Because a blood sample contains a fraction of replicating cells, an imbalance between maternal and paternal allelic fractions at this locus can sometimes be observed in genotype array data (without actual mosaicism).

#### 3.3 Estimation of true false discovery rate

Our procedure for calling the existence of a mosaic event involved identifying significant autocorrelation in phased BAF deviations using a likelihood ratio test statistic [10]. We calibrated these test statistics empirically using a permutation-based procedure (phase randomization) to obtain a nominal 5% false discovery rate (FDR) threshold. However, this permutation-based 5% FDR threshold assumed that the only source of autocorrelation in phased BAF is a true mosaic event. In reality, other sources of autocorrelation exist; in particular, we found that sample contamination produced autocorrelation in regions of long-range LD (resulting in unusual false positive calls that we subsequently filtered). While we believe that our filtering eliminated most samples affected by spurious autocorrelation, our true FDR is likely to be slightly larger than 5% due to residual artifacts.

Fortunately, we can estimate our true FDR by leveraging the fact that true-positive events should be observed more frequently in the genomes of older people, while false-positive calls (which have no relation to age) should be observed in individuals whose age distribution matches that of the study population. This observation allows us to estimate FDR by comparing the age distributions of the highest-confidence calls (17,061 calls passing a permutation-based FDR of 1%) vs. medium-confidence calls (2,571 additional calls passing a permutation-based FDR of 5% when combined with the high-confidence calls, but failing the 1% threshold). The medium-confidence call set is expected to have a false positive rate of ≈32% based on the permutation-based FDRs—meaning that its age distribution is expected to be an 68:32 mixture of (i) the age distribution of high-confidence calls and (ii) the age distribution of the study population. That is, the age distribution of medium-confidence calls should relax toward the age distribution of the overall study due to the inclusion of false positives—which is precisely what we see (Supplementary Fig. 34). (The figure also includes low-confidence calls at FDR 10% for additional context, although we did not analyze these calls.)

Upon fitting the age distribution of medium-confidence calls as a mixture of the age distribution of high-confidence calls and the overall study distribution, the regression fit gives mixture proportions of ≈56:44 rather than 68:32, implying a true FDR of 6.6% (4.5–8.6%, 95% CI) when combined with the high-confidence calls—slightly higher than the permutation-based FDR of 5%, as expected. We note that this estimate is contingent on two assumptions: (i) the high-confidence call set predominantly contains true positives (which is supported by the observation that changing the high-confidence FDR threshold from 1% to 0.1% results in a near-identical "gold standard" age distribution; and (ii) the true positives in the high-confidence and medium-confidence call set have the same age distribution. While we acknowledge that these assumptions are imperfect, this analysis gives good evidence that our FDR is well-controlled. (We also note that while we cannot completely rule out the possibility that our FDR is higher than we estimated, the key results of our paper are robust to higher FDRs than estimated; e.g., we would only expect a higher-than-estimated FDR to weaken GWAS associations and decrease effect sizes.)

### 4 Common variants influencing mCAs in *cis*

Our initial genotype–phenotype association analyses were well-powered to detect rare variants with large effects on *cis* CNN-LOH mosaicism but were underpowered to detect weaker effects of more-common variants. To maximize power to detect common variants associated with CNN-LOH mosaicism in *cis*, we performed a second genome-wide association analysis using a combined test for (i) association with CNN-LOH events and (ii) allelic bias of CNN-LOH directionality (i.e., tendency of CNN-LOH events in hets to consistently duplicate vs. delete the risk allele; Methods). For common variants, test (ii) can provide greater signal than (i): while a small fraction of individuals have CNN-LOH events on a given chromosome arm (limiting the contribution of (i)), a large fraction of cases are heterozygous (allowing substantial signal from (ii)).

This test revealed two novel associations between common variants at the *TCL1A* and *DLK1* loci on 14q and acquired 14q CNN-LOH mutations (Fisher's combined $P$=4.2×10$^{-9}$ and 3.6×10$^{-9}$; Supplementary Table 11). Intriguingly, the reference alleles at both loci were recently observed to increase risk of mosaic Y chromosome loss in elderly males, a trait related to cell proliferation and cell cycle regulation [27, 28]. Here, 14q CNN-LOH mutations in heterozygous carriers of these variants preferentially duplicate the same alleles (OR=1.91 (1.50–2.43) and 1.56 (1.28–1.90)), corroborating a pro-proliferative effect. (The *DLK1* locus also lies within an imprinted region that has previously been observed to be the target of parental bias in 14q CNN-LOH mutations [32], raising the possibility of interaction between allelic and parental effects; however, familial data will be needed to investigate further.)

We also searched for associations between inherited variants and copy-number-altering mCAs (i.e., loss and gain events) in *cis* but did not find any associations aside from the *FRA10B* locus, at which we previously observed that fragile alleles confer risk of mosaic 10q deletions [10].

### 5 Inherited variants associated with mCAs in *trans*

A common haplotype in *TERT* broadly increases risk of clonal hematopoiesis involving any mosaic point mutation [8], and common variants also exert *trans* effects on the likelihood of mosaic *JAK2* V617F mutation [57] and sex chromosome loss [10, 27, 28, 58]. To identify more inherited haplotypes that similarly modify risk of autosomal mosaic chromosomal alterations in general, we conducted a genome-wide association analysis between common variants and presence of any detectable autosomal mCA (Methods). Three loci reached significance ($P$<5×10$^{-8}$): *TERT* ($P$=6.9×10$^{-18}$, OR=1.11 (1.08–1.14) for rs7705526), *TERC* ($P$=2.9×10$^{-8}$, OR=0.93 (0.91–0.96) for rs12638862), and *SP140* ($P$=9.4×10$^{-9}$, OR=1.08 (1.05–1.10) for rs62191195) (Supplementary Table 12).

The associations at *TERT* and *TERC* suggest that genetic differences in telomere maintenance make some individuals more susceptible to clonal expansions than others. Consistent with this hypothesis, we further observed that two additional telomere-length-associated SNPs [59] at *OBFC1* and *RTEL1* also associated with mCA susceptibility at nominal $P$<0.05 significance (Supplementary Table 19). For 6 out of 7 SNPs previously associated with telomere length [59], the telomere length-increasing allele exhibited a risk-increasing effect sign for mCAs (Supplementary Table 19).

The lead associated variant in *SP140* matches the second-strongest association for chronic lymphocytic leukemia (CLL) [60], and we further observed that most CLL risk alleles increase risk of clonal hematopoiesis involving CLL-related +12 or 13q LOH events (see below). These results demonstrate that common variants play a modest, quantitative role in altering processes that facilitate clonal expansion (in contrast to and in addition to the larger *cis* effects of variants on which mosaic CNN-LOH mutations directly act by changing allele dosage).

### 6 Shared genetic risk of CLL and mosaic +12 and 13q LOH

Chronic lymphocytic leukemia (CLL) is a highly heritable hematological malignancy, with 42 risk loci identified to date by GWAS on up to 6,200 cases [60–68]. Given the relatively large number of carriers of CLL-associated mosaic events in the UK Biobank data set ($\sim$2,000), we sought to investigate the extent to which CLL risk alleles also influence risk of clonal hematopoiesis involving CLL-associated chromosomal alterations. We examined the two types of mCAs most strongly associated with CLL: mosaic trisomy 12 ("+12") and mosaic 13q LOH spanning *DLEU7* (including both del(13q) and 13q CNN-LOH to maximize power).

For each of 46 independent lead variants at the 42 previously identified CLL risk loci [60], we performed association tests with three case-control phenotypes: mosaic +12, mosaic 13q LOH, and (as a check) CLL in UK Biobank. We restricted each association test to individuals who reported European ancestry, and we pruned to unrelated subsets of samples as in our *cis* GWAS analyses (Methods). For mosaic +12, we tested 634 cases, defined as individuals with a whole-chromosome 12 mosaic event (including unclassified event as well as events confidently classified as gains), and 378,107 controls, defined as individuals with no chr12 mosaic event. For 13q LOH, we tested 914 cases, defined as individuals with loss or CNN-LOH events spanning *DLEU7*, and 378,048 controls, defined as individuals with no chr13 mosaic event. For CLL, we tested 656 cases, defined as individuals with any reported CLL (prevalent or incident), and 378,606 controls. These case sets contained modest overlap: of the 634 mosaic +12 cases, 78 had prevalent or incident CLL, and of the 914 mosaic 13q LOH cases, 243 had prevalent or incident CLL. Thirty-five individuals had both mosaic +12 and mosaic 13q LOH; 14 of the 35 also had prevalent or incident CLL. We performed association tests on imputed genotypes using logistic regression in `plink` [38] adjusting for age and sex as covariates. We verified that previously reported CLL risk variants replicated well in UK Biobank, with a regression coefficient of 0.85 (0.75–0.96) for observed vs. expected log(OR) and consistent effect directions for 43 of 46 variants (Supplementary Fig. 37a and Supplementary Table 20).

We observed that most CLL risk variants conferred risk for either mosaic +12, mosaic 13q LOH, or both (Supplementary Fig. 37b,c and Supplementary Table 20). Of the 46 CLL risk alleles tested, 40 had risk-increasing effect directions on mosaic +12, with 21 reaching nominal significance ($P$<0.05). For mosaic 13q LOH, 42 of 46 CLL risk alleles had risk-increasing effects, with 29 reaching nominal significance (Supplementary Table 20). Broadly, the estimated effects of these 46 risk alleles on mosaic +12 and 13q LOH risk were moderately smaller than their reported effects on CLL (log(OR) regression coefficients of 0.52 (0.40–0.64) and 0.63 (0.53–0.73), respectively).

Interestingly, we observed heterogeneity in the effects of variants on mosaic +12 vs. mosaic 13q LOH risk: some variants appeared to influence one form of mosaicism much more than the other (Supplementary Fig. 37d). We quantified this genetic heterogeneity by performing bivariate BOLT-REML analysis [69] and estimated a genetic correlation of 0.79 (0.09)—significantly less than 1—between mosaic +12 and mosaic 13q LOH risk. These results suggest that different genetic mechanisms may lead to different subtypes of CLL that manifest different chromosomal alterations.

### 7 Estimation of mortality risk conferred by mCAs

UK death registry data provided by UK Biobank reported 10,498 deaths on or before December 31, 2015 (the censoring date suggested by UK Biobank) from among 415,867 individuals of self-reported European ancestry with no evidence of potential undiagnosed blood cancer based on anomalous blood counts (Methods). This censoring date corresponded to a median follow-up time of 6.9 years (range 5–10 years).

While the relatively small number of deaths in the cohort limited our power to identify links between mCAs and mortality, we observed that clones with gain of chromosome 8 (which contains the *MYC* oncogene) associated with increased mortality even in individuals with normal blood cell counts ($P$=$3.5\times10^{-5}$), reaching Bonferroni significance among the 78 events tested, presumably due to large effect size (OR=5.10 (2.41–9.85)). In this analysis we applied statistical tests analogous to our cancer outcome analyses, using Cochran-Mantel-Haenszel (CMH) tests to adjust for sex and age while accounting for case-control imbalance.

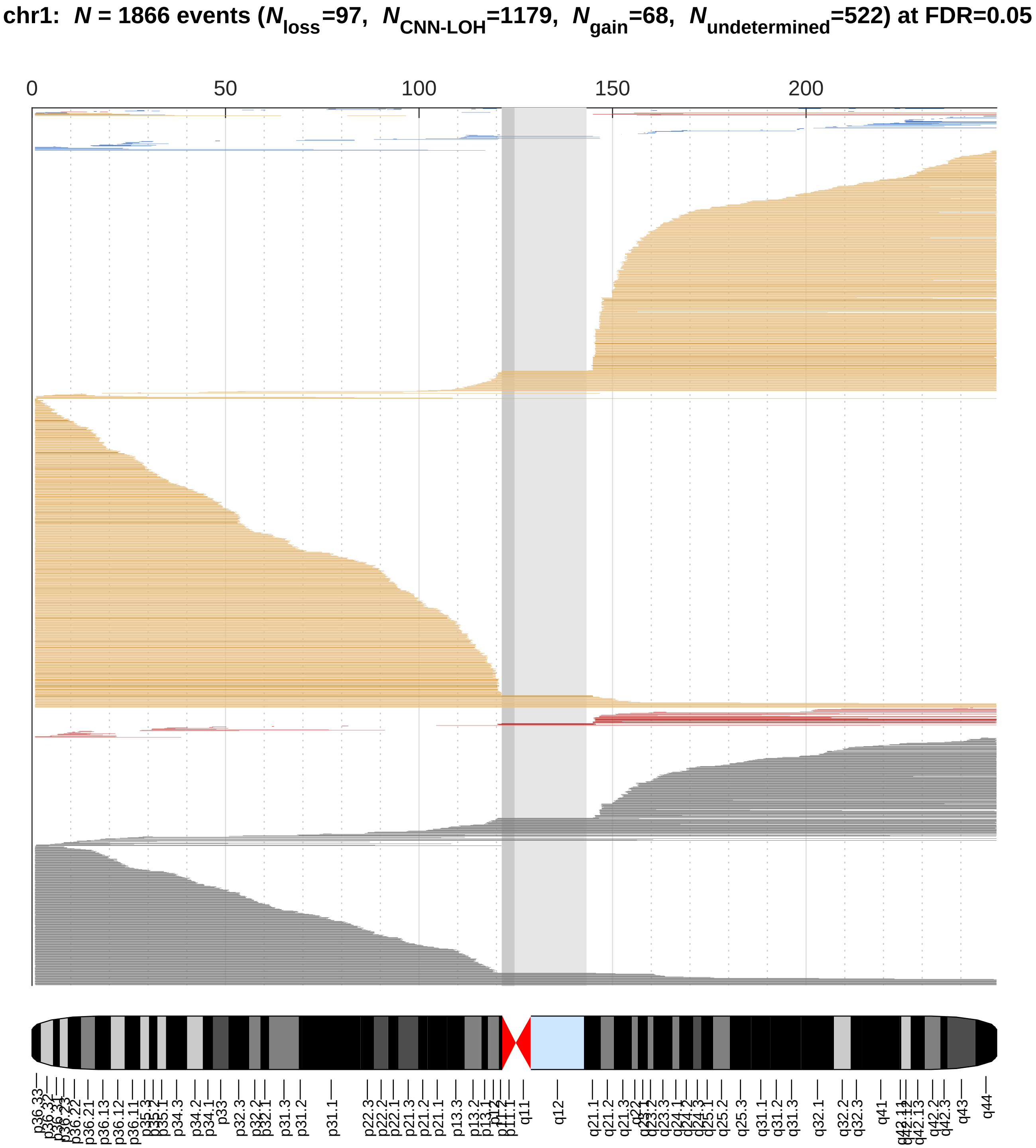

**Supplementary Figure 1. Detected mCAs on chromosome 1.** Events are color-coded by copy-number: loss (blue), CNN-LOH (orange), gain (red), undetermined (grey). Darker coloring indicates higher allelic fraction. Multiple events within a single individual are plotted with the same y-coordinate (at the top of the plot). Note that events with unknown copy number also generally have greater uncertainty in their boundaries due to low allelic fraction.

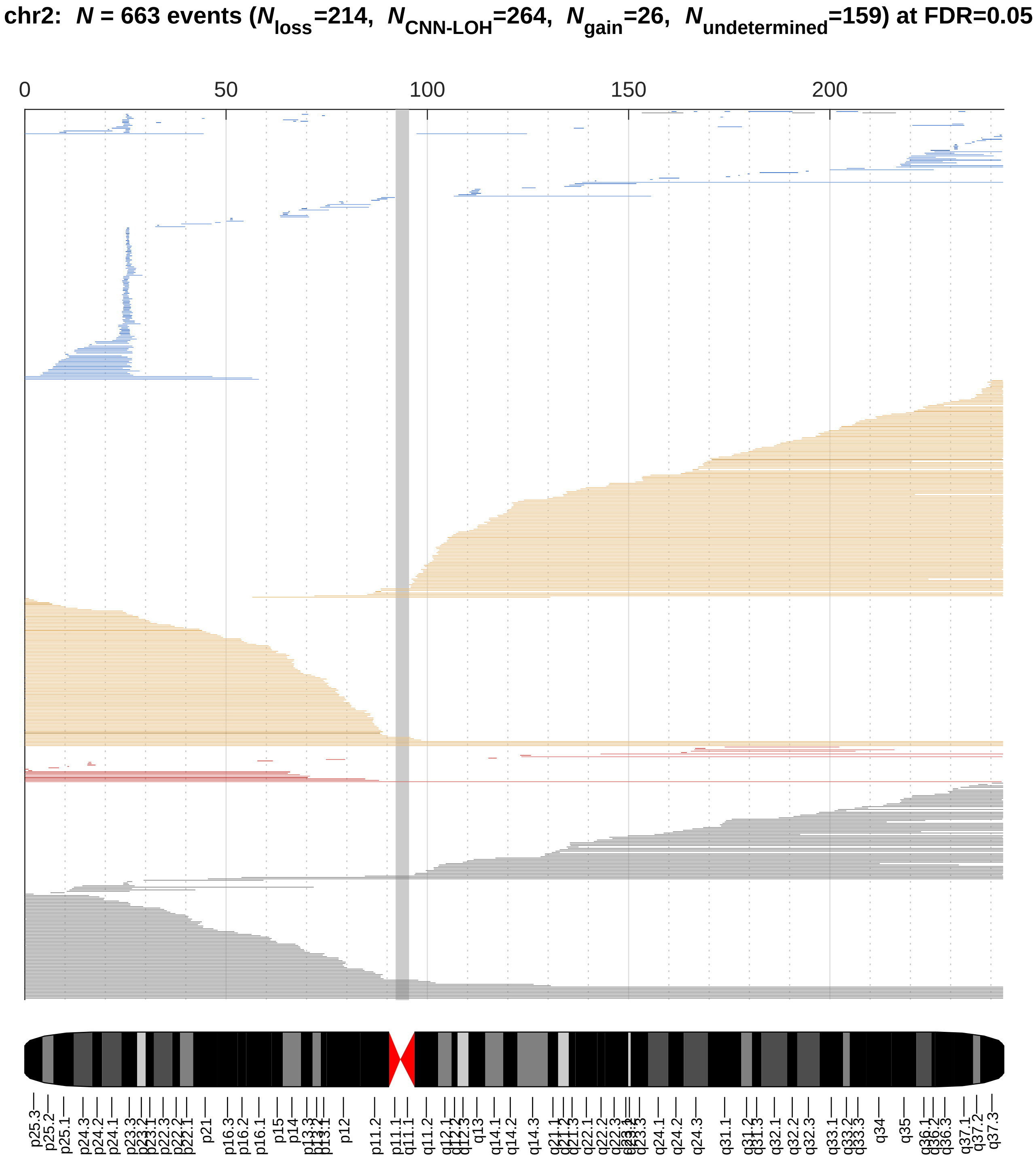

**Supplementary Figure 2. Detected mCAs on chromosome 2.** Events are color-coded by copy-number: loss (blue), CNN-LOH (orange), gain (red), undetermined (grey). Darker coloring indicates higher allelic fraction. Multiple events within a single individual are plotted with the same y-coordinate (at the top of the plot). Note that events with unknown copy number also generally have greater uncertainty in their boundaries due to low allelic fraction.

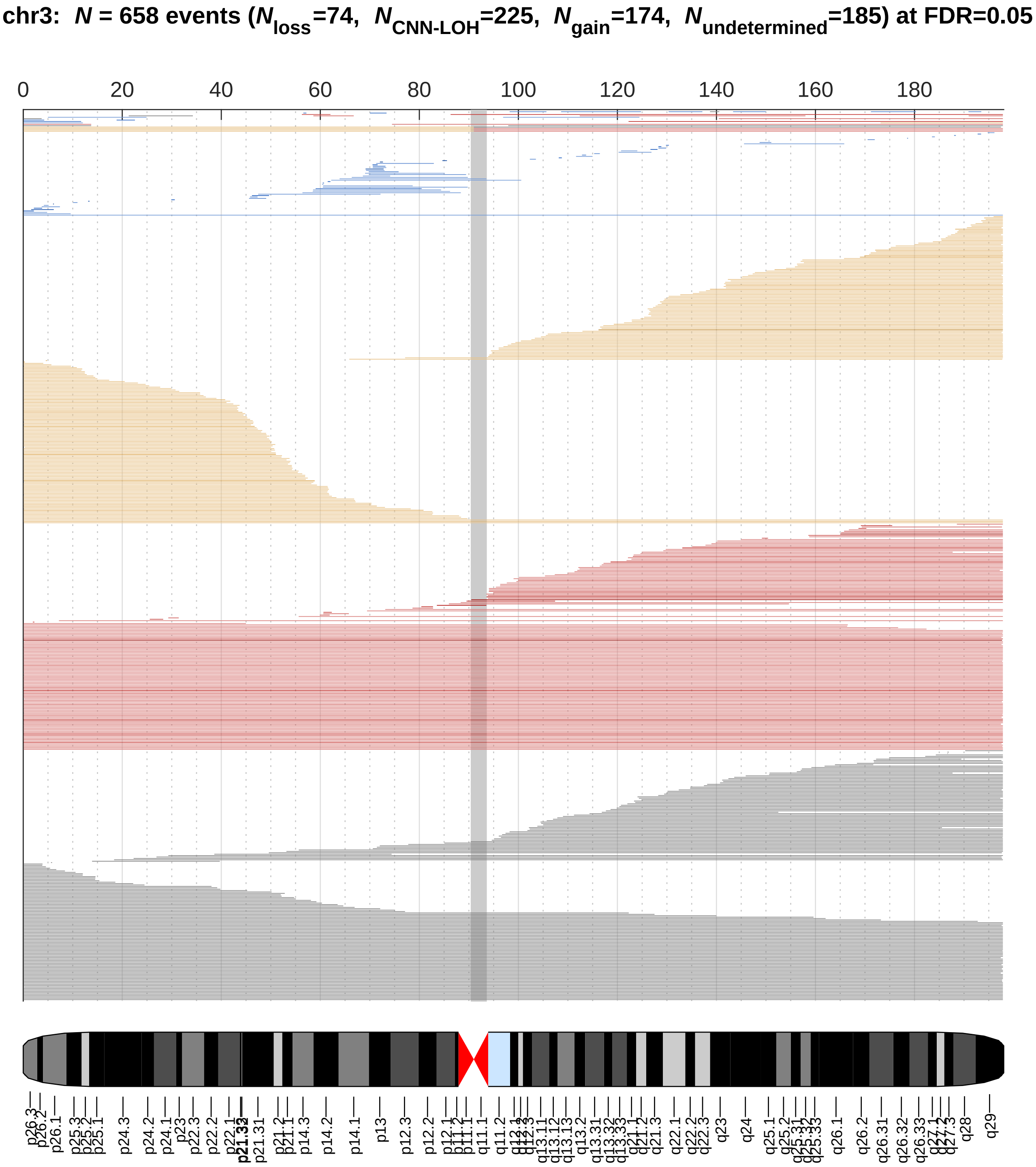

**Supplementary Figure 3. Detected mCAs on chromosome 3.** Events are color-coded by copy-number: loss (blue), CNN-LOH (orange), gain (red), undetermined (grey). Darker coloring indicates higher allelic fraction. Multiple events within a single individual are plotted with the same y-coordinate (at the top of the plot). Note that events with unknown copy number also generally have greater uncertainty in their boundaries due to low allelic fraction.

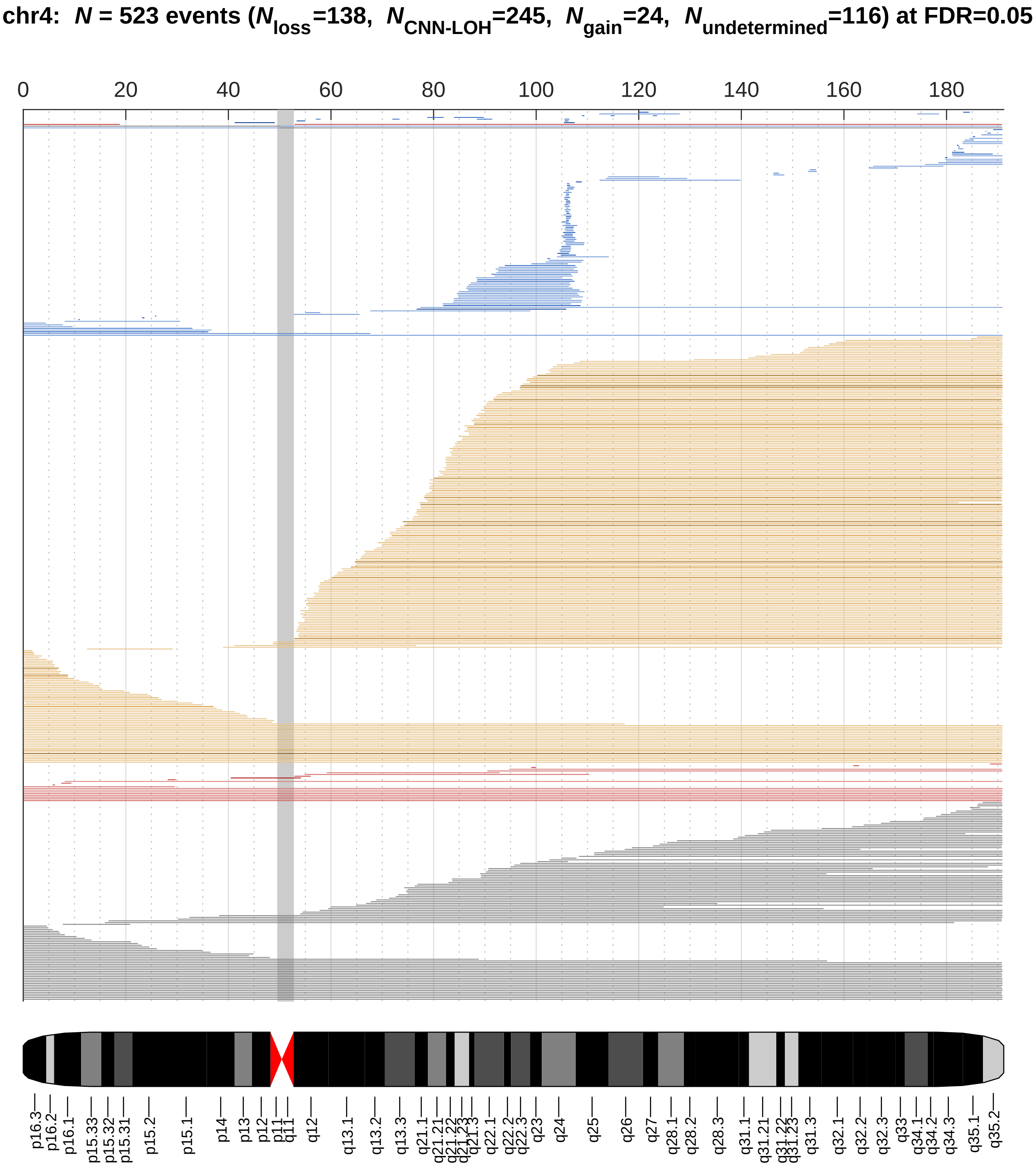

**Supplementary Figure 4. Detected mCAs on chromosome 4.** Events are color-coded by copy-number: loss (blue), CNN-LOH (orange), gain (red), undetermined (grey). Darker coloring indicates higher allelic fraction. Multiple events within a single individual are plotted with the same y-coordinate (at the top of the plot). Note that events with unknown copy number also generally have greater uncertainty in their boundaries due to low allelic fraction.

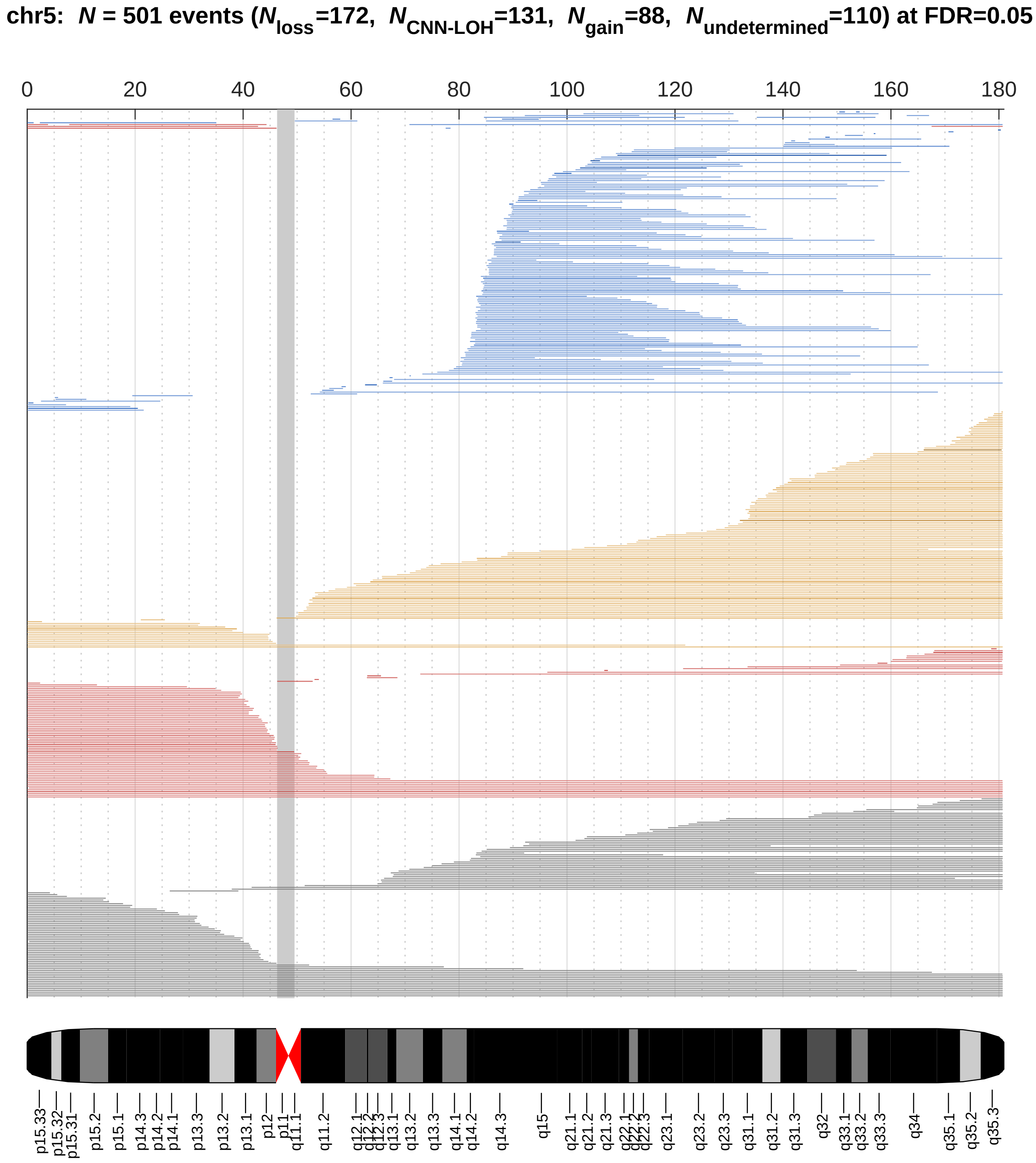

**Supplementary Figure 5. Detected mCAs on chromosome 5.** Events are color-coded by copy-number: loss (blue), CNN-LOH (orange), gain (red), undetermined (grey). Darker coloring indicates higher allelic fraction. Multiple events within a single individual are plotted with the same y-coordinate (at the top of the plot). Note that events with unknown copy number also generally have greater uncertainty in their boundaries due to low allelic fraction.

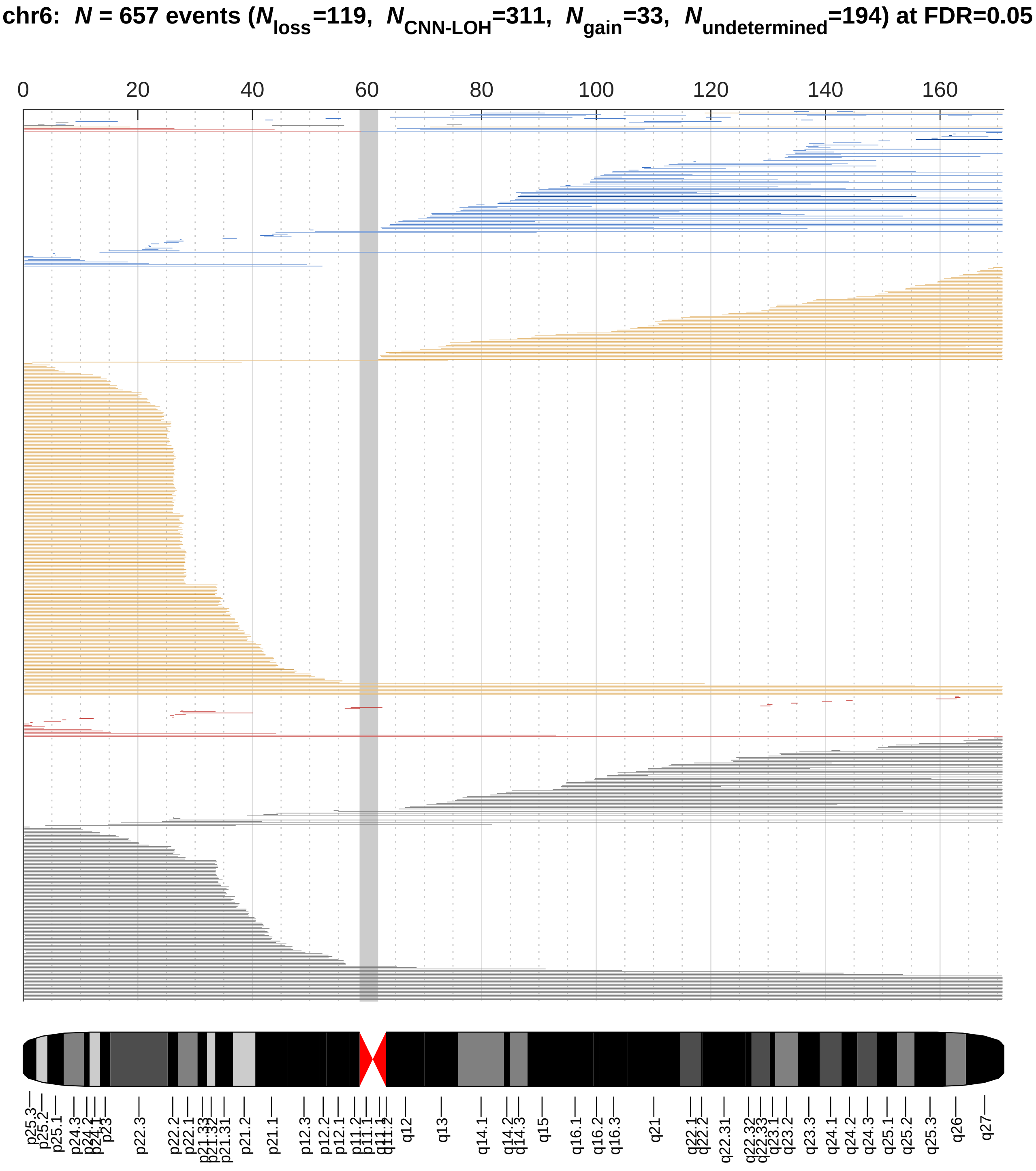

**Supplementary Figure 6. Detected mCAs on chromosome 6.** Events are color-coded by copy-number: loss (blue), CNN-LOH (orange), gain (red), undetermined (grey). Darker coloring indicates higher allelic fraction. Multiple events within a single individual are plotted with the same y-coordinate (at the top of the plot). Note that events with unknown copy number also generally have greater uncertainty in their boundaries due to low allelic fraction.

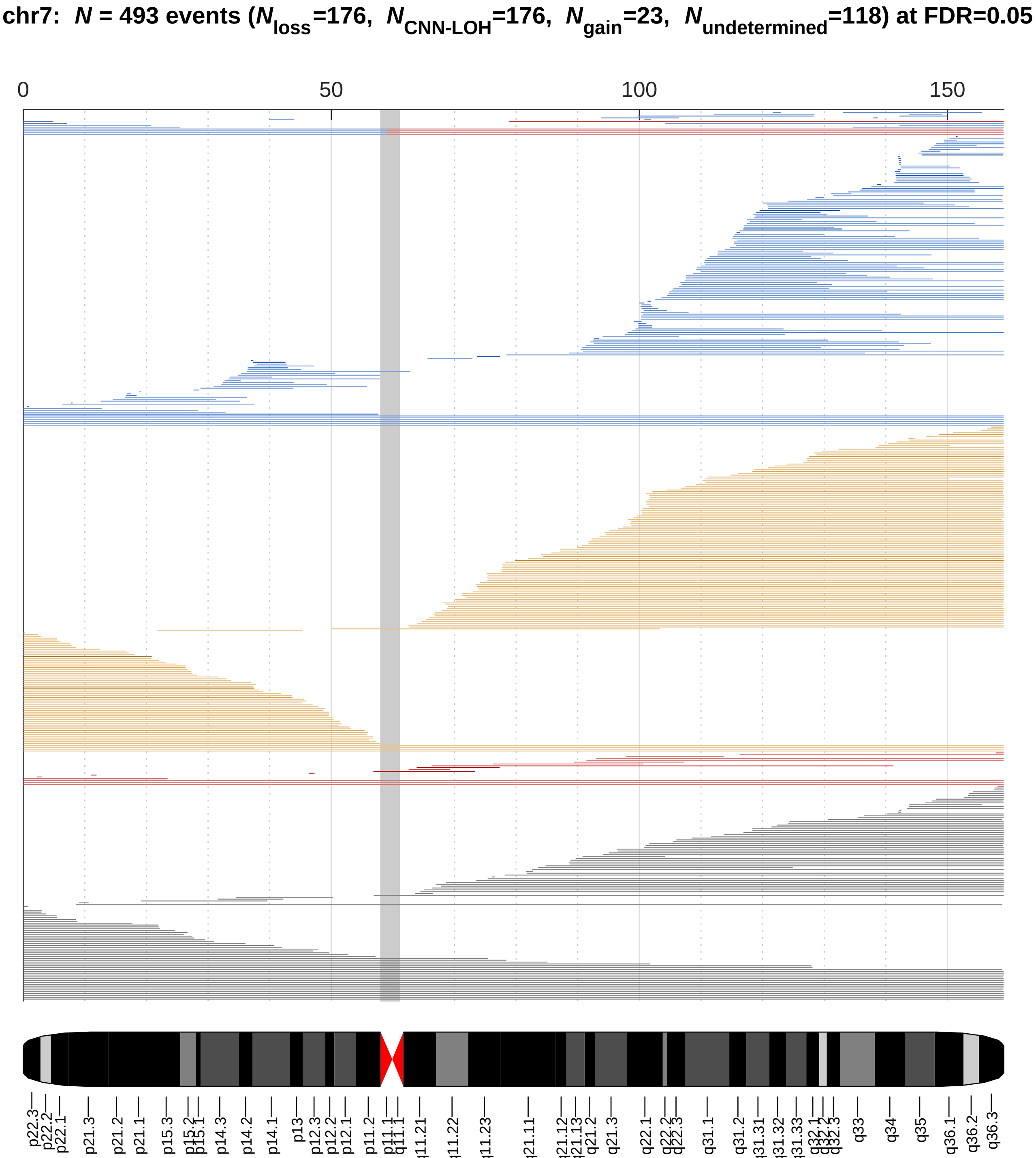

**Supplementary Figure 7. Detected mCAs on chromosome 7.** Events are color-coded by copy-number: loss (blue), CNN-LOH (orange), gain (red), undetermined (grey). Darker coloring indicates higher allelic fraction. Multiple events within a single individual are plotted with the same y-coordinate (at the top of the plot). Note that events with unknown copy number also generally have greater uncertainty in their boundaries due to low allelic fraction.

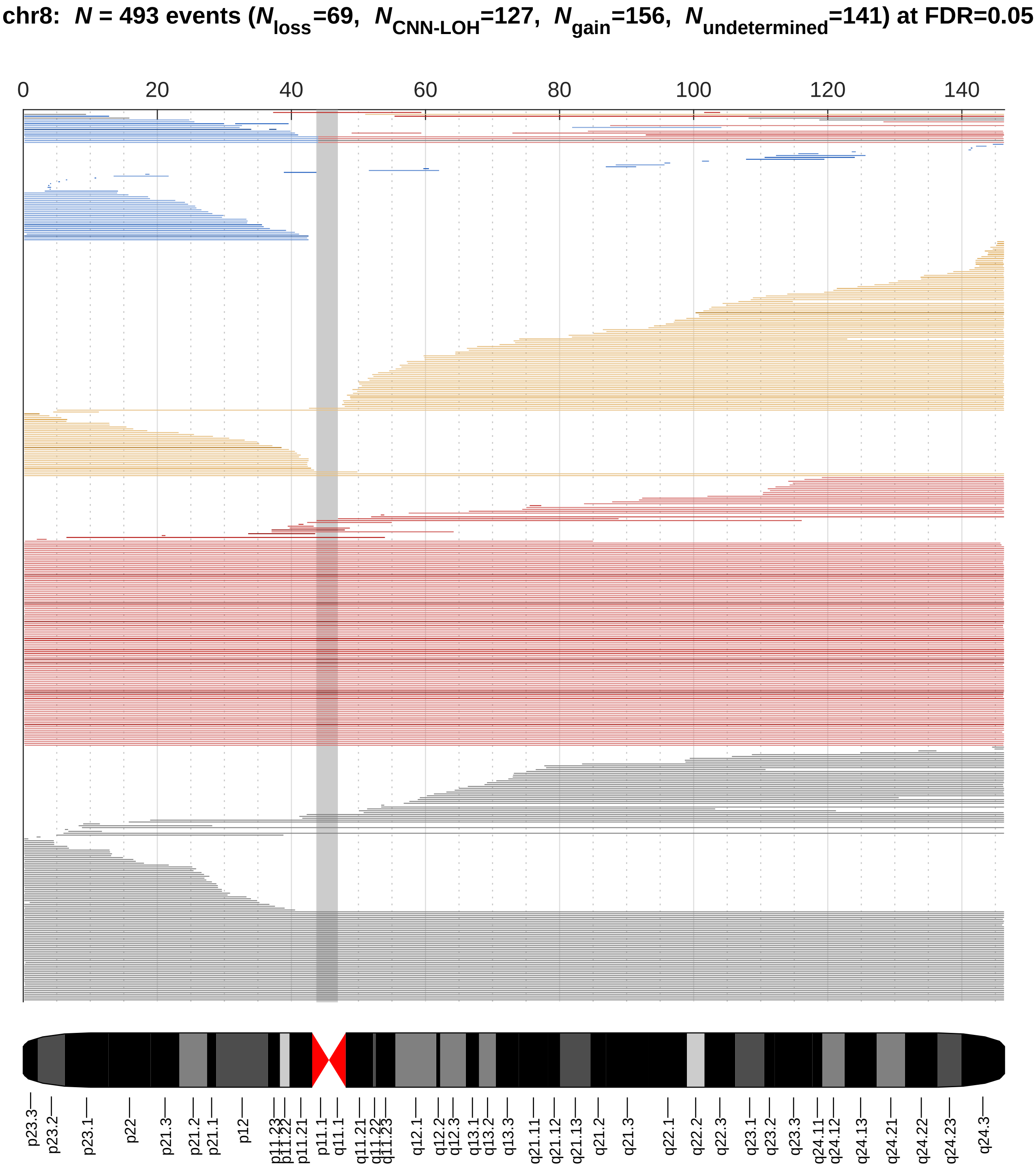

**Supplementary Figure 8. Detected mCAs on chromosome 8.** Events are color-coded by copy-number: loss (blue), CNN-LOH (orange), gain (red), undetermined (grey). Darker coloring indicates higher allelic fraction. Multiple events within a single individual are plotted with the same y-coordinate (at the top of the plot). Note that events with unknown copy number also generally have greater uncertainty in their boundaries due to low allelic fraction.

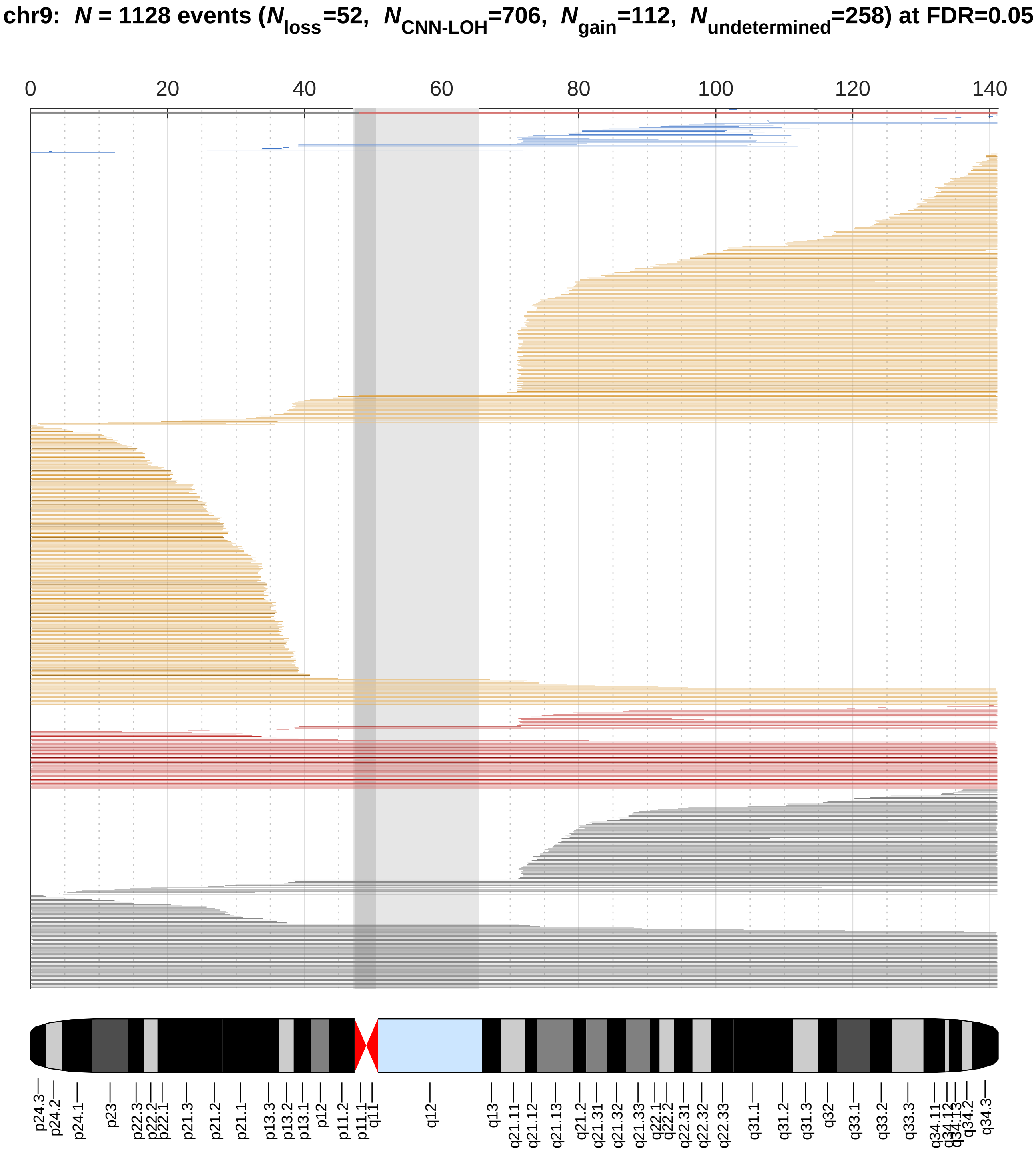

**Supplementary Figure 9. Detected mCAs on chromosome 9.** Events are color-coded by copy-number: loss (blue), CNN-LOH (orange), gain (red), undetermined (grey). Darker coloring indicates higher allelic fraction. Multiple events within a single individual are plotted with the same y-coordinate (at the top of the plot). Note that events with unknown copy number also generally have greater uncertainty in their boundaries due to low allelic fraction.

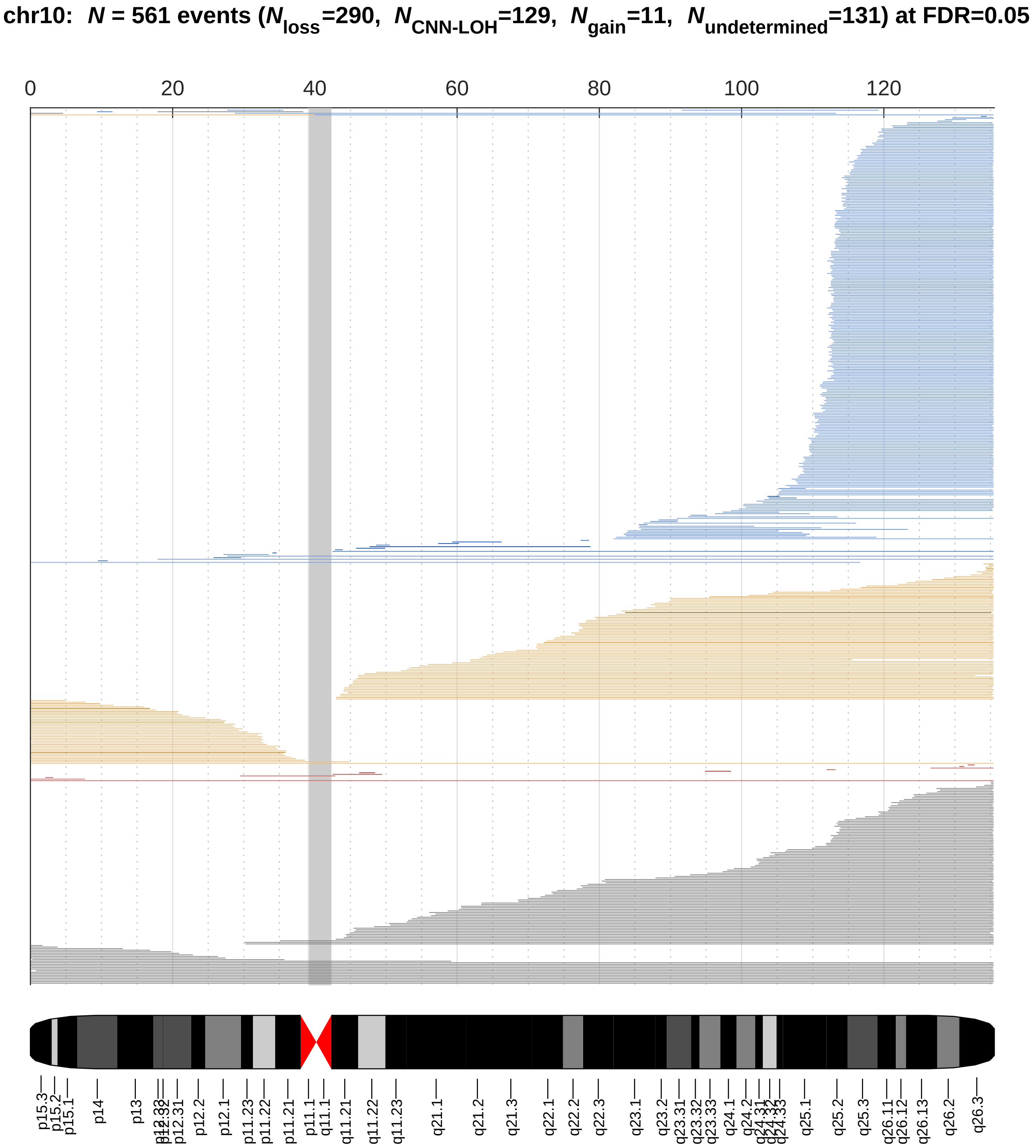

**Supplementary Figure 10. Detected mCAs on chromosome 10.** Events are color-coded by copy-number: loss (blue), CNN-LOH (orange), gain (red), undetermined (grey). Darker coloring indicates higher allelic fraction. Multiple events within a single individual are plotted with the same y-coordinate (at the top of the plot). Note that events with unknown copy number also generally have greater uncertainty in their boundaries due to low allelic fraction.

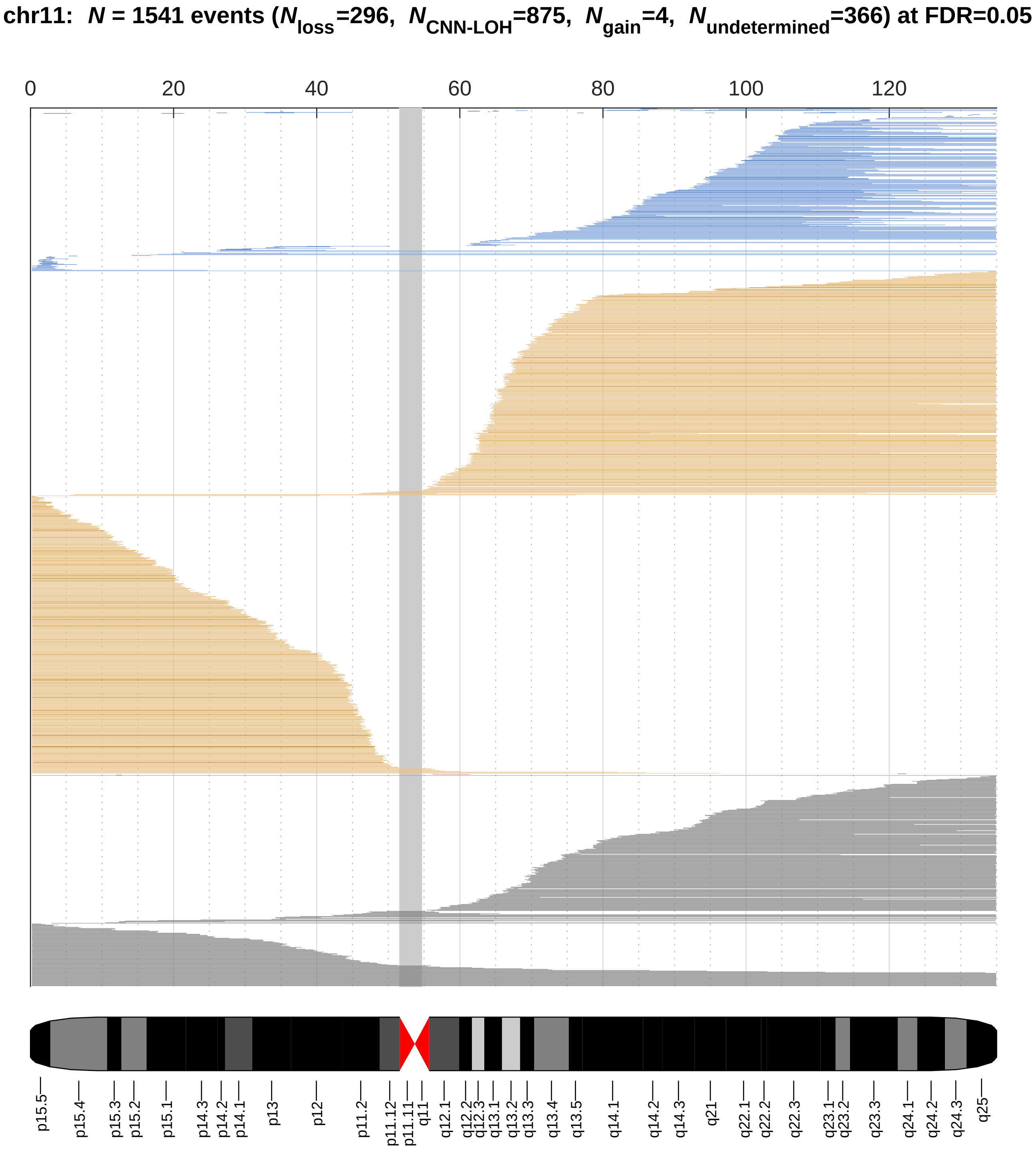

**Supplementary Figure 11. Detected mCAs on chromosome 11.** Events are color-coded by copy-number: loss (blue), CNN-LOH (orange), gain (red), undetermined (grey). Darker coloring indicates higher allelic fraction. Multiple events within a single individual are plotted with the same y-coordinate (at the top of the plot). Note that events with unknown copy number also generally have greater uncertainty in their boundaries due to low allelic fraction.

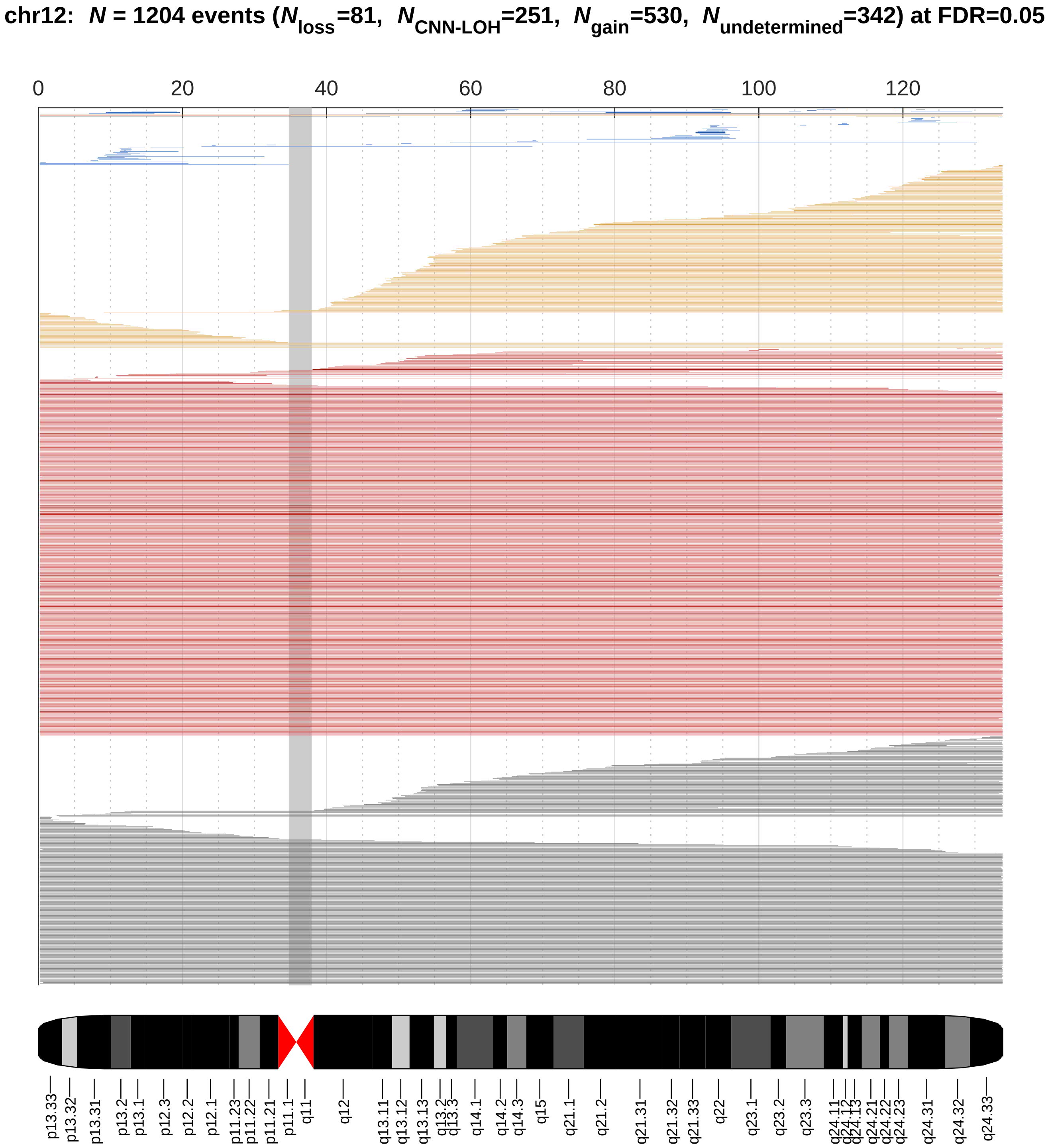

**Supplementary Figure 12. Detected mCAs on chromosome 12.** Events are color-coded by copy-number: loss (blue), CNN-LOH (orange), gain (red), undetermined (grey). Darker coloring indicates higher allelic fraction. Multiple events within a single individual are plotted with the same y-coordinate (at the top of the plot). Note that events with unknown copy number also generally have greater uncertainty in their boundaries due to low allelic fraction.

**chr13: $N$ = 1289 events ($N_{loss}$=596, $N_{CNN\text{-}LOH}$=444, $N_{gain}$=12, $N_{undetermined}$=237) at FDR=0.05**

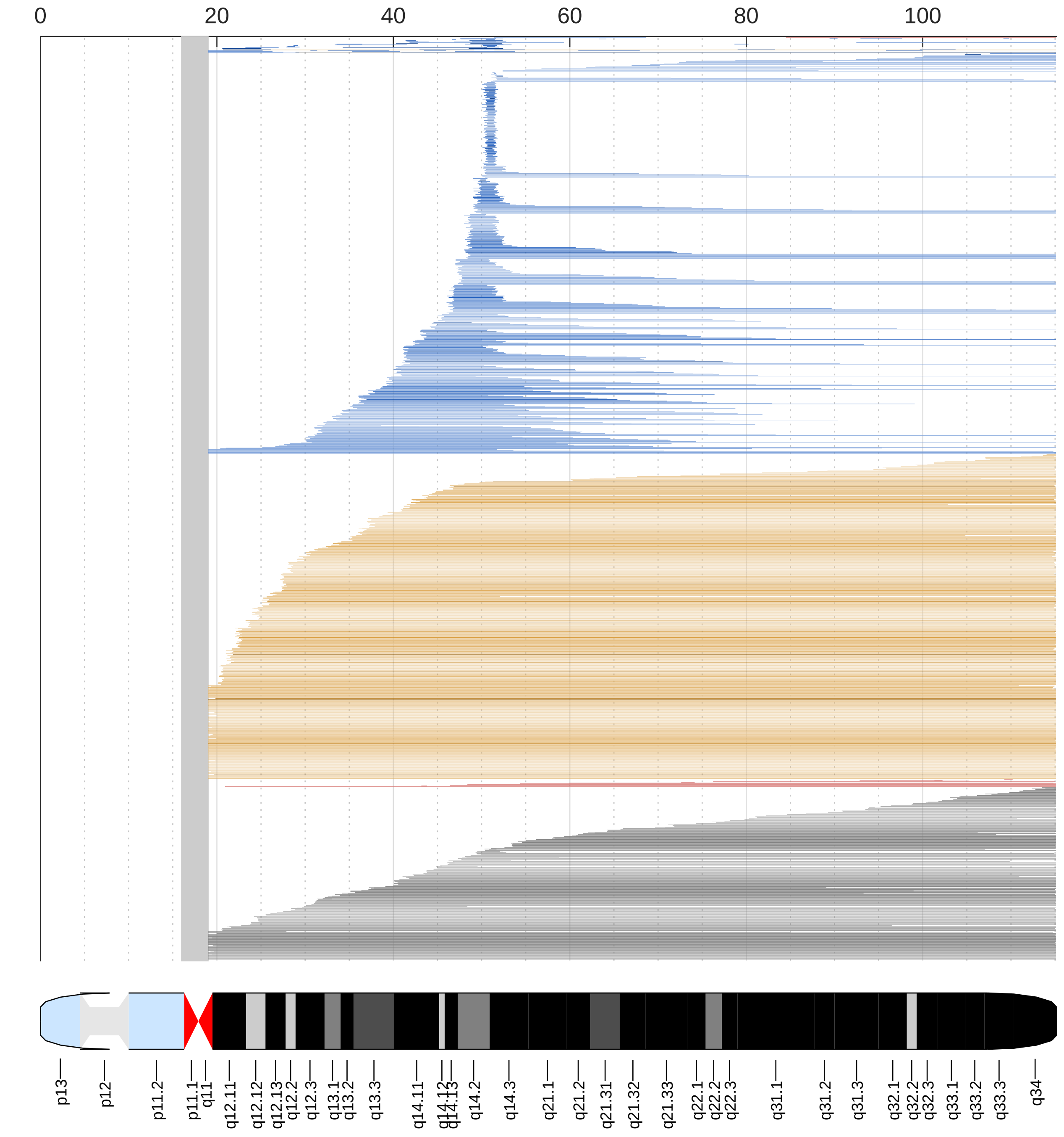

**Supplementary Figure 13. Detected mCAs on chromosome 13.** Events are color-coded by copy-number: loss (blue), CNN-LOH (orange), gain (red), undetermined (grey). Darker coloring indicates higher allelic fraction. Multiple events within a single individual are plotted with the same y-coordinate (at the top of the plot). Note that events with unknown copy number also generally have greater uncertainty in their boundaries due to low allelic fraction.

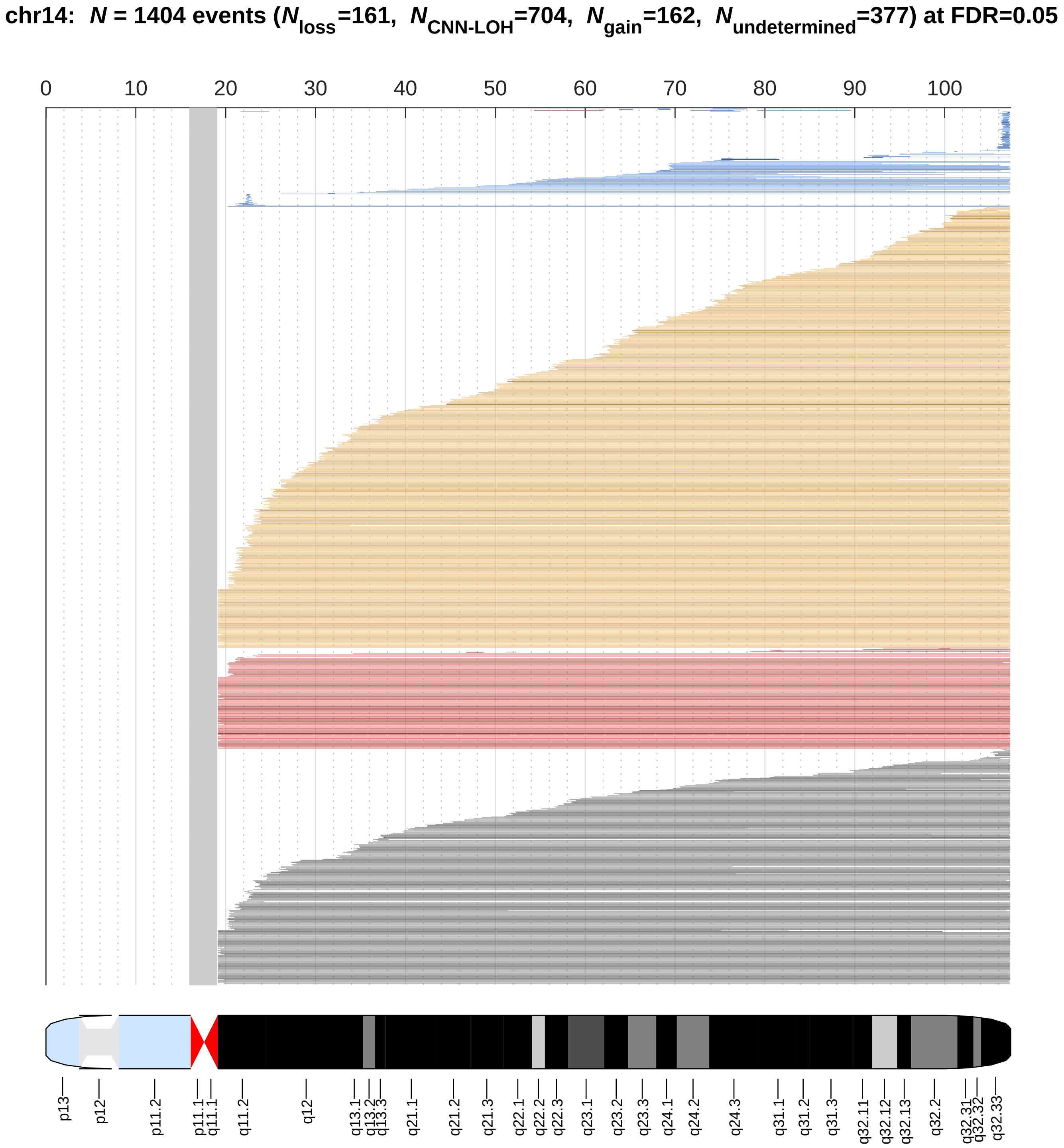

**Supplementary Figure 14. Detected mCAs on chromosome 14.** Events are color-coded by copy-number: loss (blue), CNN-LOH (orange), gain (red), undetermined (grey). Darker coloring indicates higher allelic fraction. Multiple events within a single individual are plotted with the same y-coordinate (at the top of the plot). Note that events with unknown copy number also generally have greater uncertainty in their boundaries due to low allelic fraction.

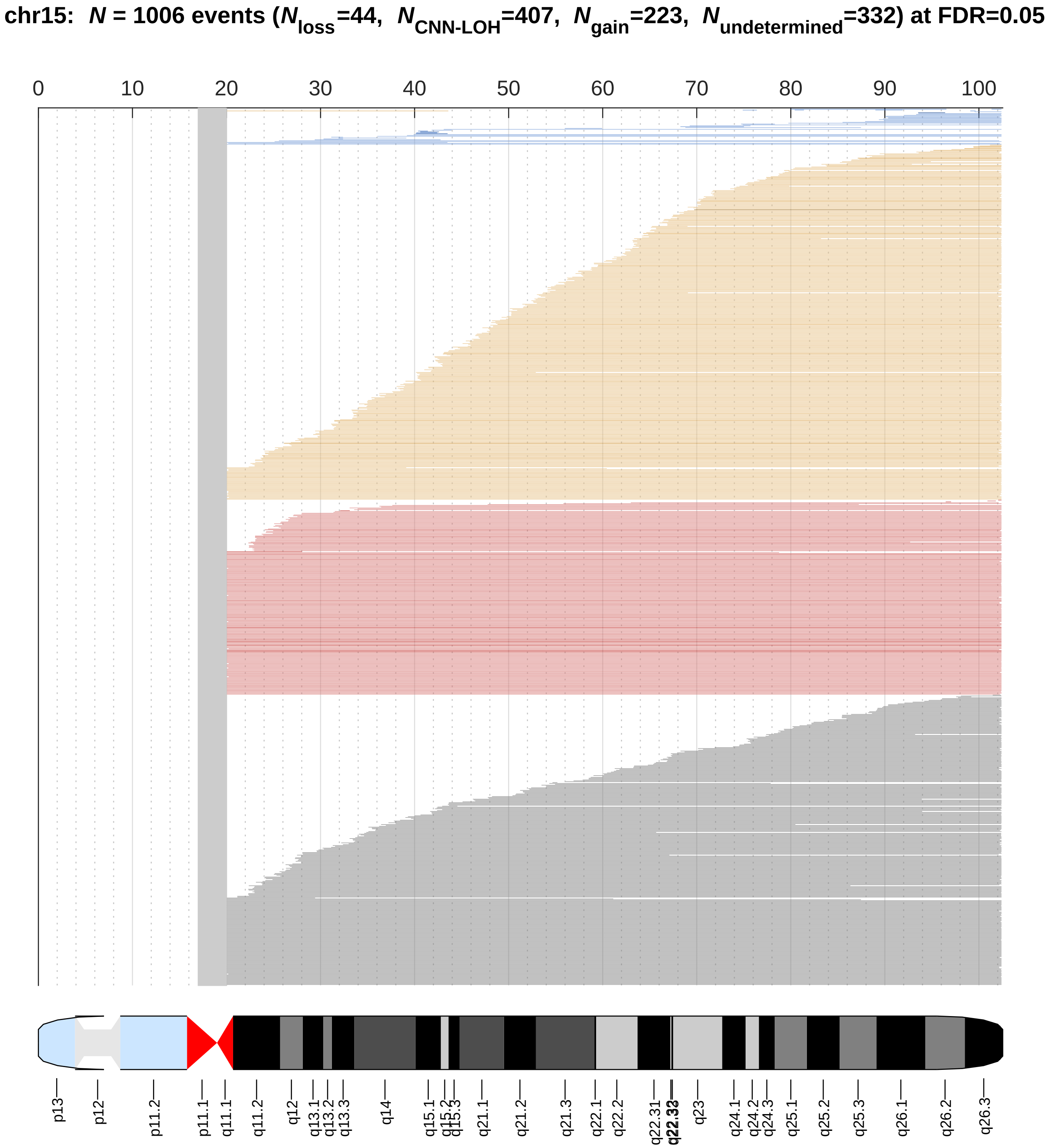

**Supplementary Figure 15. Detected mCAs on chromosome 15.** Events are color-coded by copy-number: loss (blue), CNN-LOH (orange), gain (red), undetermined (grey). Darker coloring indicates higher allelic fraction. Multiple events within a single individual are plotted with the same y-coordinate (at the top of the plot). Note that events with unknown copy number also generally have greater uncertainty in their boundaries due to low allelic fraction.

chr16: $N$ = 860 events ($N_{loss}$=180, $N_{CNN\text{-}LOH}$=470, $N_{gain}$=8, $N_{undetermined}$=202) at FDR=0.05

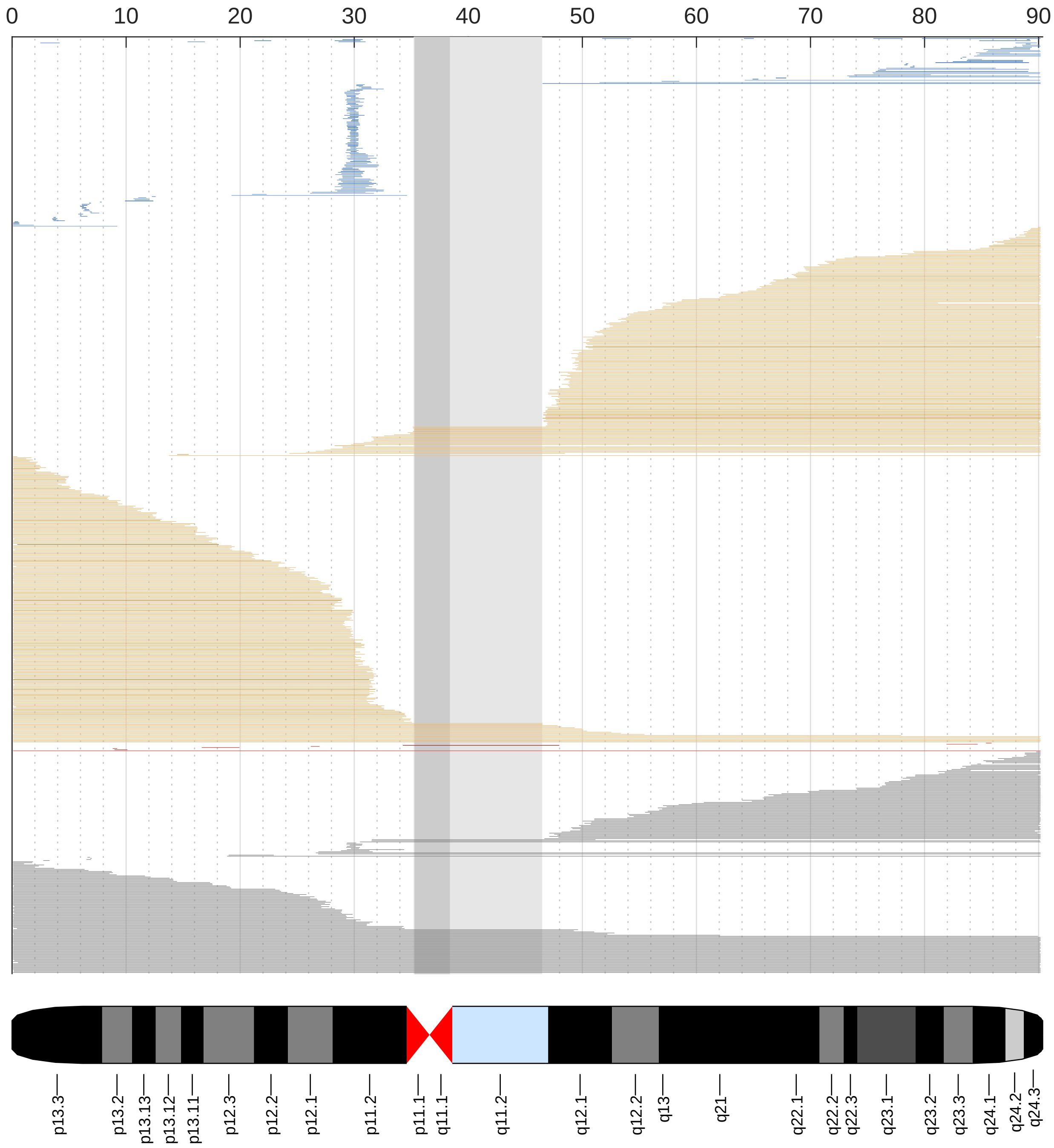

**Supplementary Figure 16. Detected mCAs on chromosome 16.** Events are color-coded by copy-number: loss (blue), CNN-LOH (orange), gain (red), undetermined (grey). Darker coloring indicates higher allelic fraction. Multiple events within a single individual are plotted with the same y-coordinate (at the top of the plot). Note that events with unknown copy number also generally have greater uncertainty in their boundaries due to low allelic fraction.

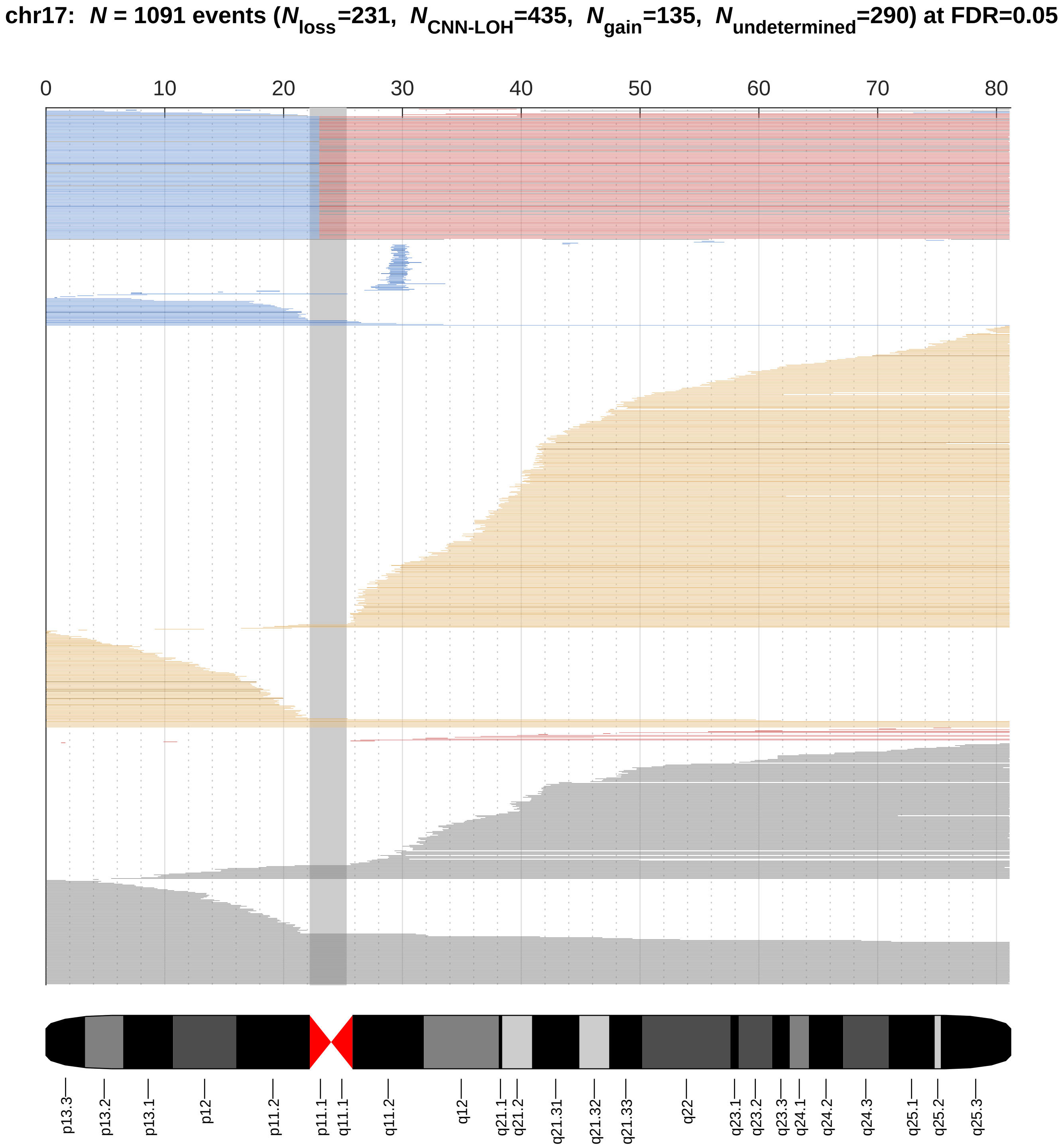

**Supplementary Figure 17. Detected mCAs on chromosome 17.** Events are color-coded by copy-number: loss (blue), CNN-LOH (orange), gain (red), undetermined (grey). Darker coloring indicates higher allelic fraction. Multiple events within a single individual are plotted with the same y-coordinate (at the top of the plot). Note that events with unknown copy number also generally have greater uncertainty in their boundaries due to low allelic fraction.

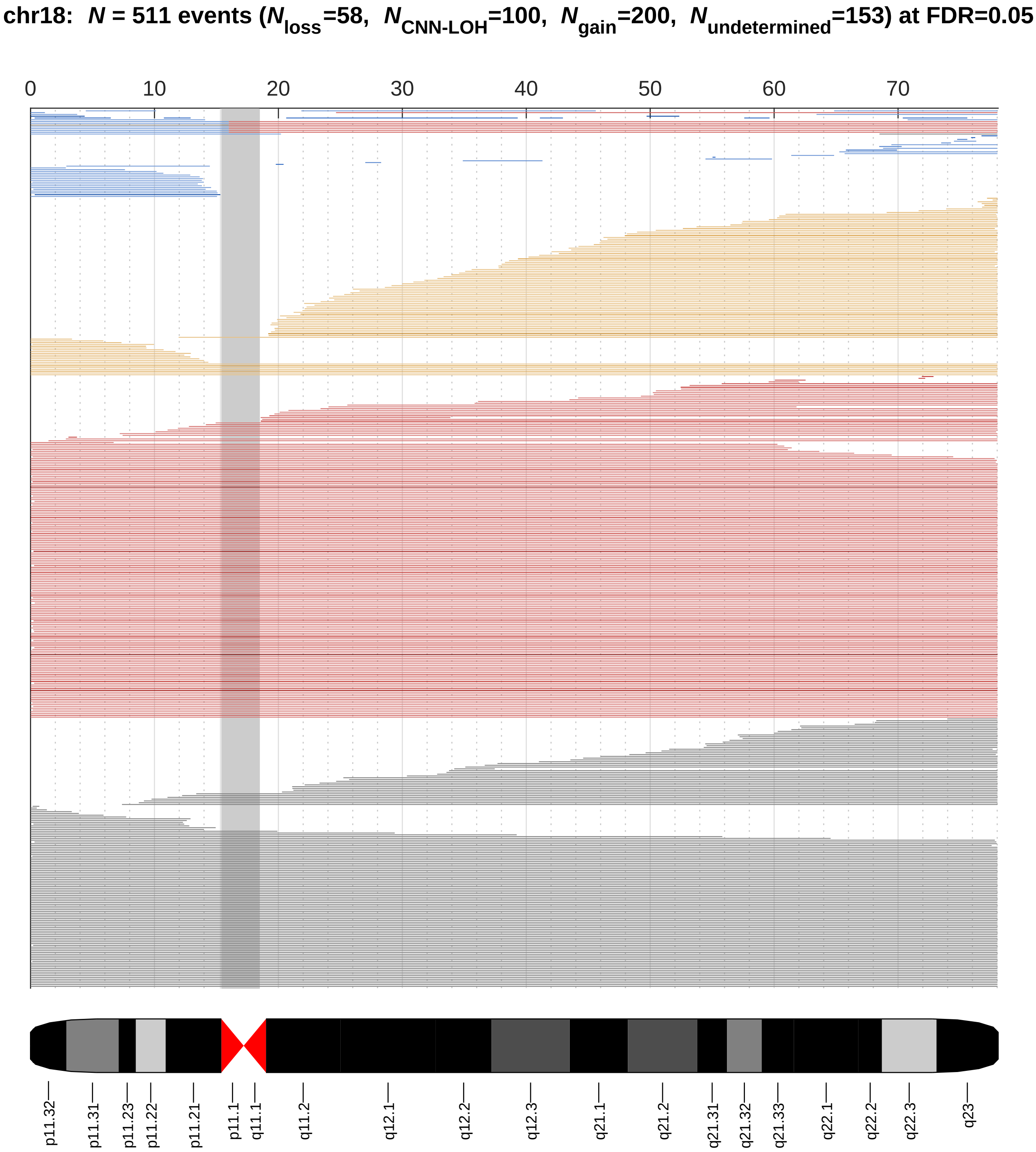

**Supplementary Figure 18. Detected mCAs on chromosome 18.** Events are color-coded by copy-number: loss (blue), CNN-LOH (orange), gain (red), undetermined (grey). Darker coloring indicates higher allelic fraction. Multiple events within a single individual are plotted with the same y-coordinate (at the top of the plot). Note that events with unknown copy number also generally have greater uncertainty in their boundaries due to low allelic fraction.

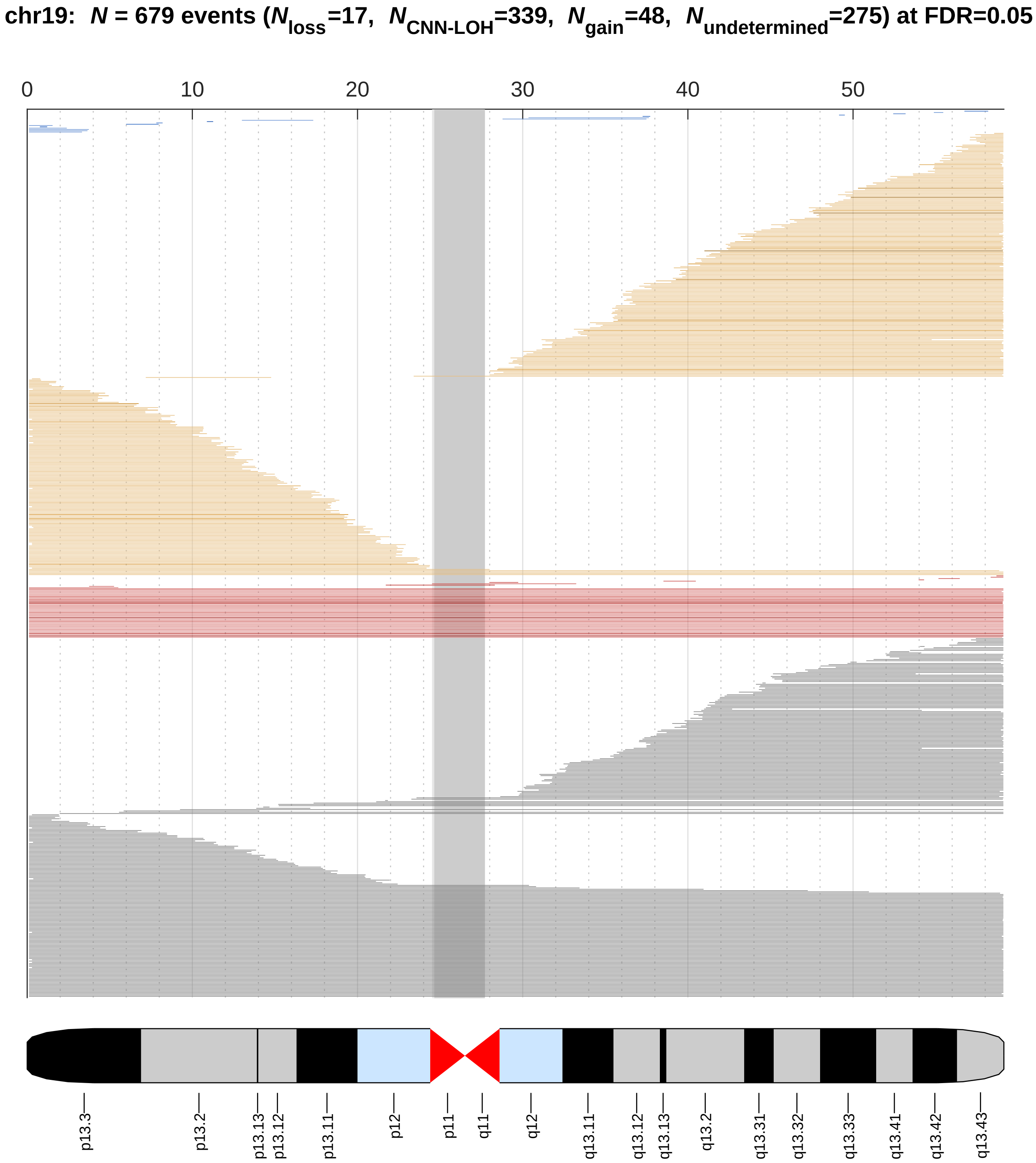

**Supplementary Figure 19. Detected mCAs on chromosome 19.** Events are color-coded by copy-number: loss (blue), CNN-LOH (orange), gain (red), undetermined (grey). Darker coloring indicates higher allelic fraction. Multiple events within a single individual are plotted with the same y-coordinate (at the top of the plot). Note that events with unknown copy number also generally have greater uncertainty in their boundaries due to low allelic fraction.

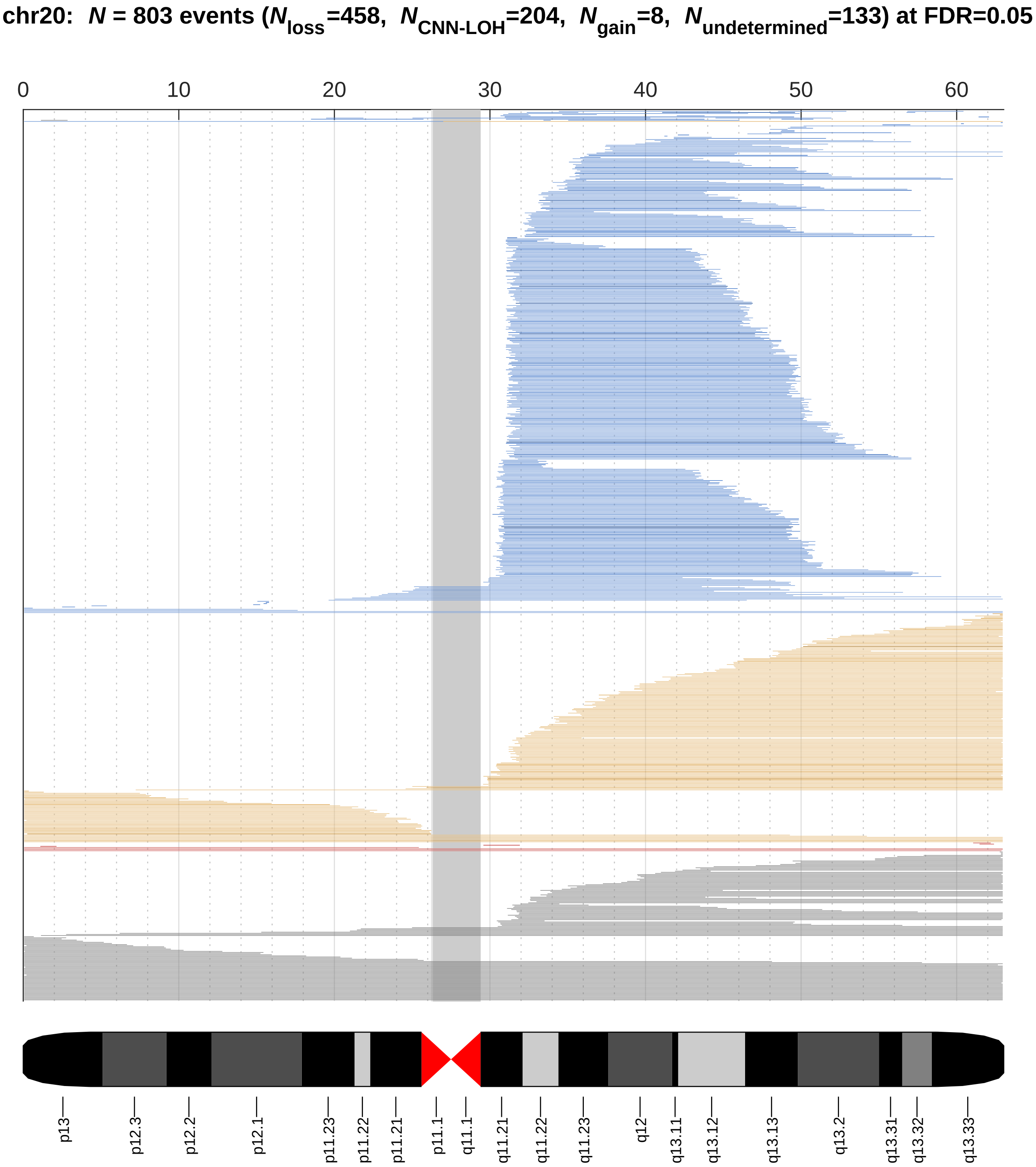

**Supplementary Figure 20. Detected mCAs on chromosome 20.** Events are color-coded by copy-number: loss (blue), CNN-LOH (orange), gain (red), undetermined (grey). Darker coloring indicates higher allelic fraction. Multiple events within a single individual are plotted with the same y-coordinate (at the top of the plot). Note that events with unknown copy number also generally have greater uncertainty in their boundaries due to low allelic fraction.

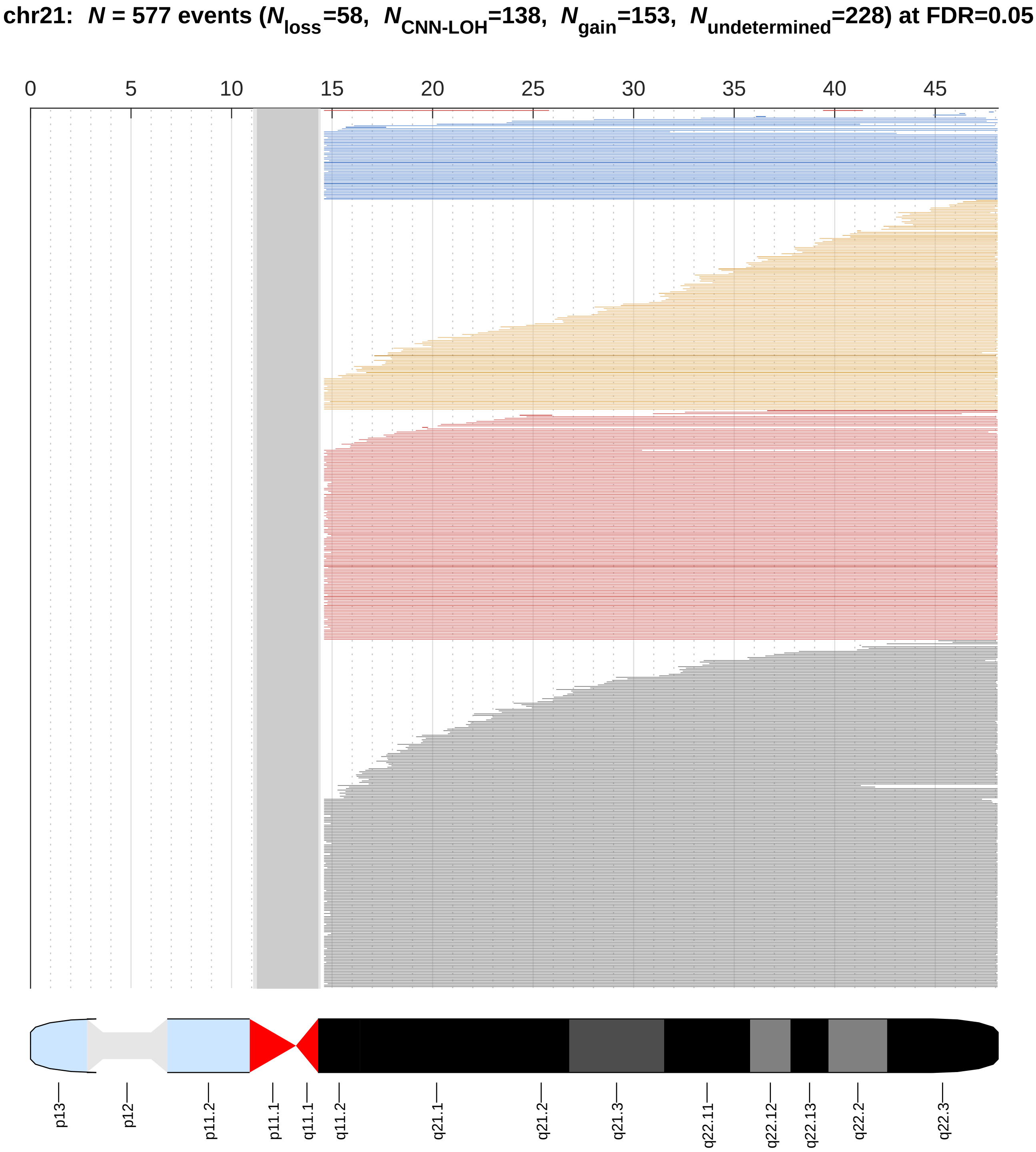

**Supplementary Figure 21. Detected mCAs on chromosome 21.** Events are color-coded by copy-number: loss (blue), CNN-LOH (orange), gain (red), undetermined (grey). Darker coloring indicates higher allelic fraction. Multiple events within a single individual are plotted with the same y-coordinate (at the top of the plot). Note that events with unknown copy number also generally have greater uncertainty in their boundaries due to low allelic fraction.

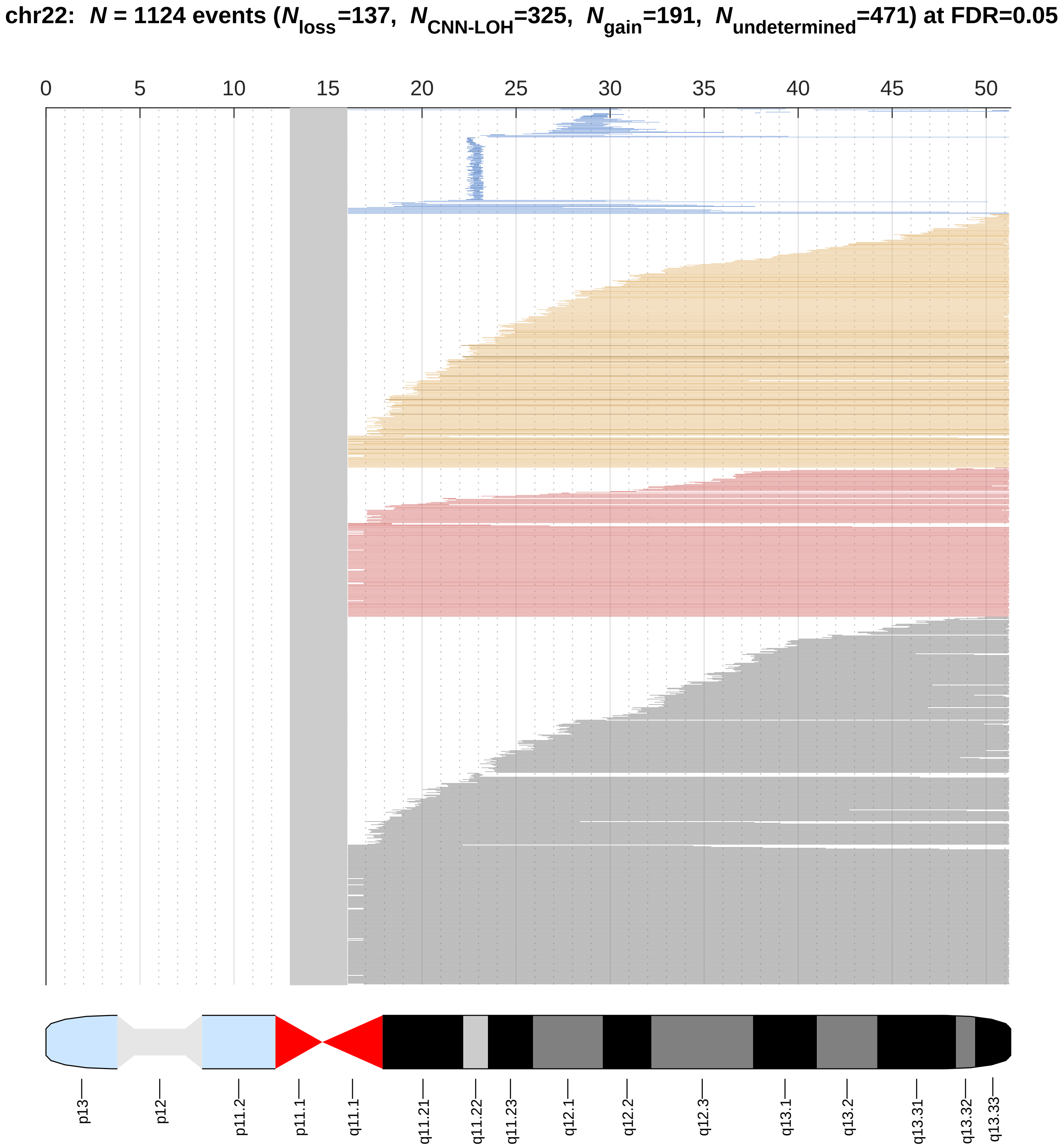

**Supplementary Figure 22. Detected mCAs on chromosome 22.** Events are color-coded by copy-number: loss (blue), CNN-LOH (orange), gain (red), undetermined (grey). Darker coloring indicates higher allelic fraction. Multiple events within a single individual are plotted with the same y-coordinate (at the top of the plot). Note that events with unknown copy number also generally have greater uncertainty in their boundaries due to low allelic fraction.

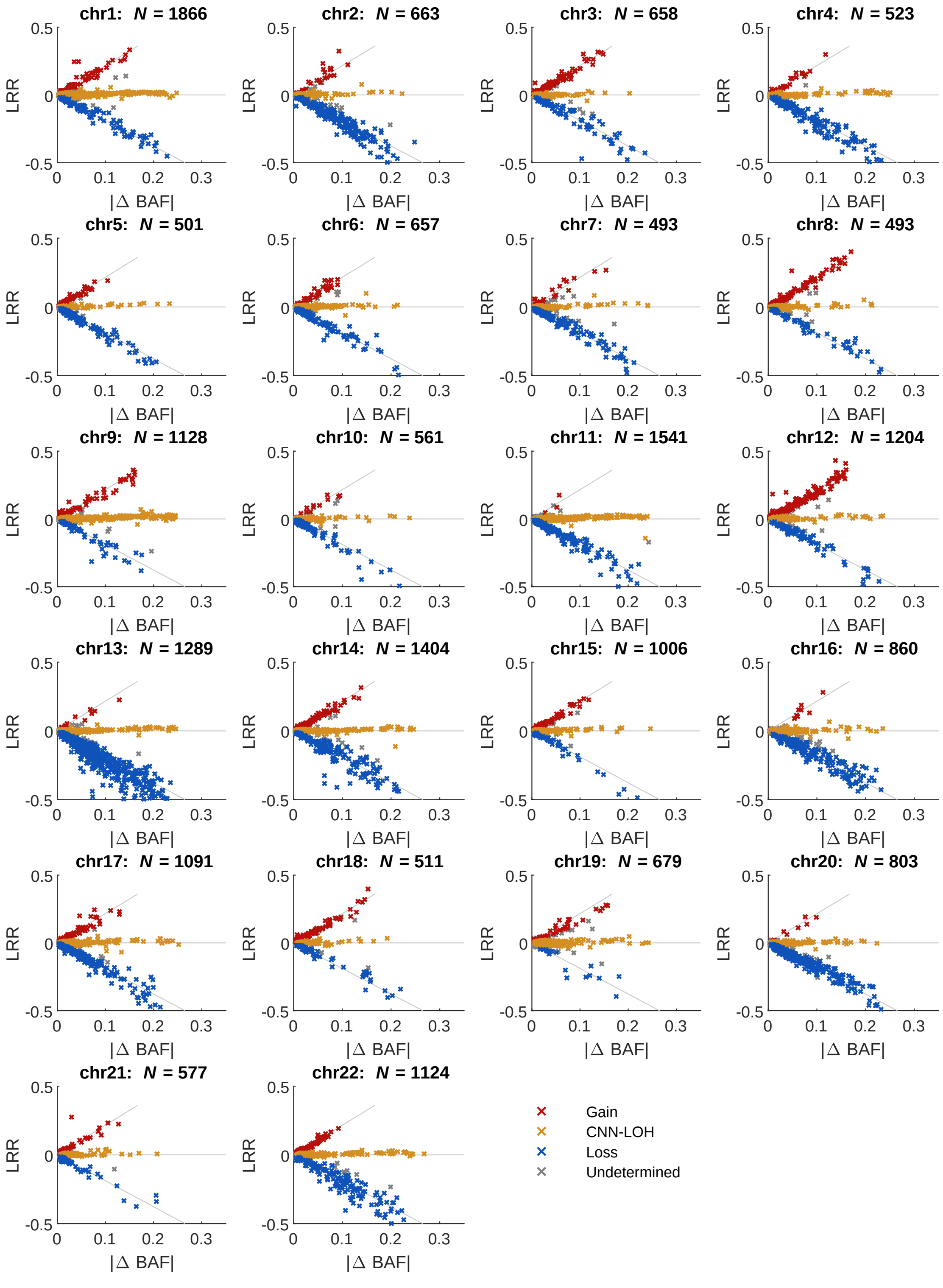

**Supplementary Figure 23. Total vs. relative allelic intensities of mCAs detected on each chromosome.** For each of $N$ mCAs, mean $\log_2$ R ratio (LRR) of each detected mCA is plotted against estimated change in B allele frequency at heterozygous sites ($|\Delta\text{BAF}|$). The data exhibits the characteristic "arrowhead" pattern in which LRR/$|\Delta\text{BAF}|$ approximately equals a positive constant for gain events, zero for CNN-LOH events, and a negative constant for loss events. This pattern is very consistent across chromosomes.

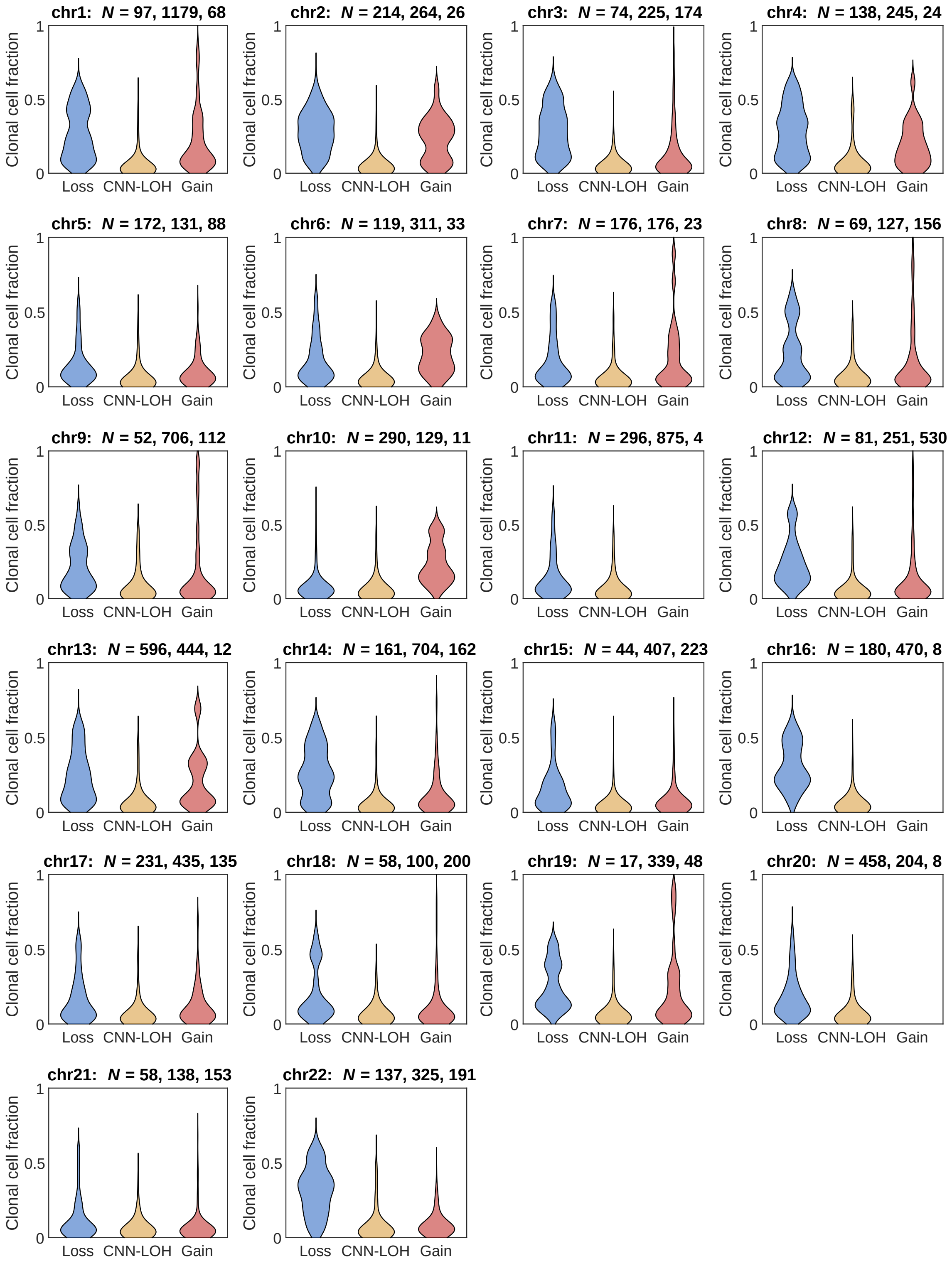

**Supplementary Figure 24. Extent of clonal proliferation of mCAs detected on each chromosome.** For each of $N$ mCAs called as a loss, CNN-LOH, or gain, we estimate its allelic fraction (i.e., fraction of blood cells with the mCA) from LRR and $|\Delta\text{BAF}|$. The violin plots show allelic fraction distributions stratified by chromosome and copy number (whenever at least ten events were called).

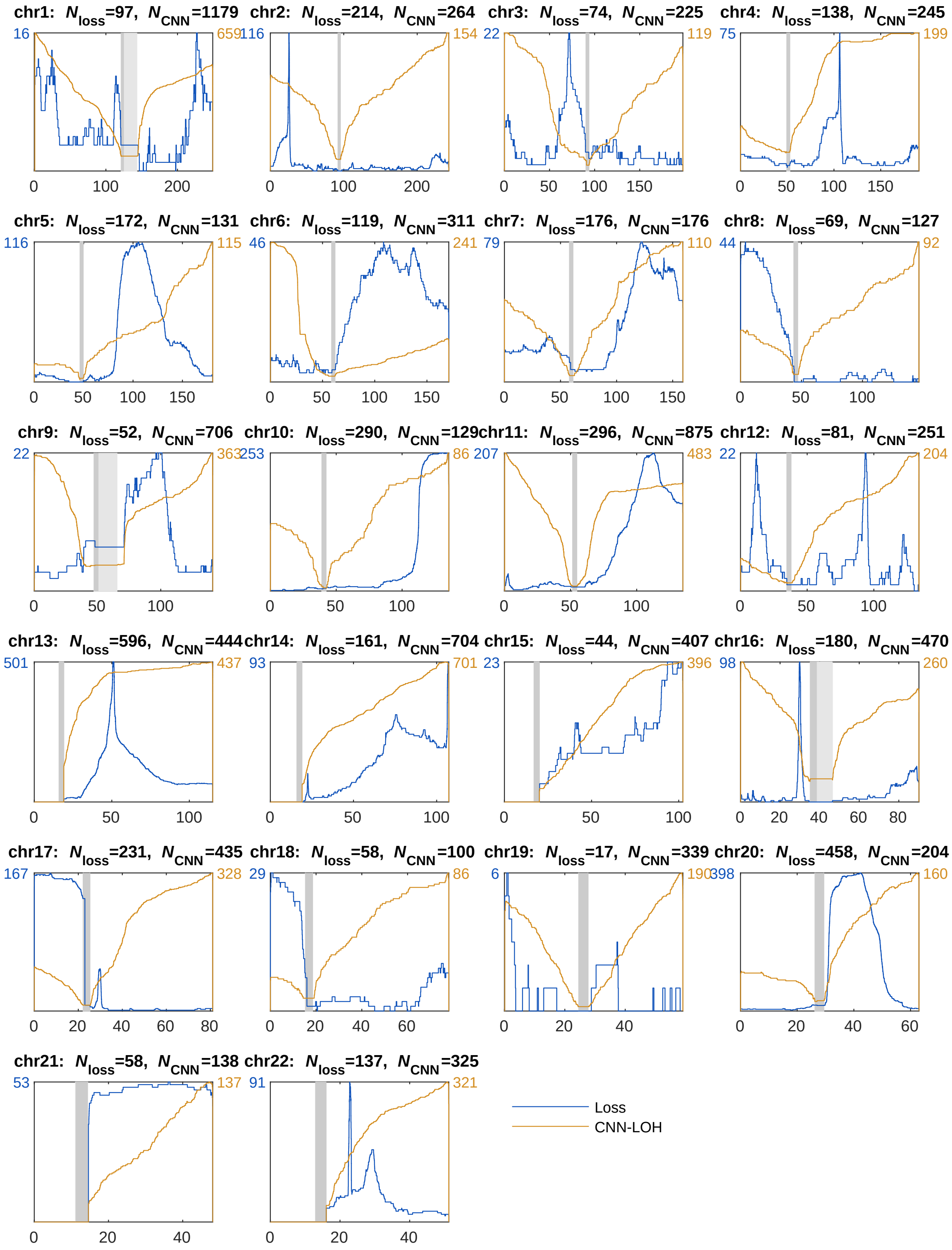

**Supplementary Figure 25. Genomic coverage by mosaic loss and CNN-LOH events.** The blue and orange curves indicate the total numbers of detected mosaic loss (blue) and CNN-LOH (orange) events covering each position in the genome.

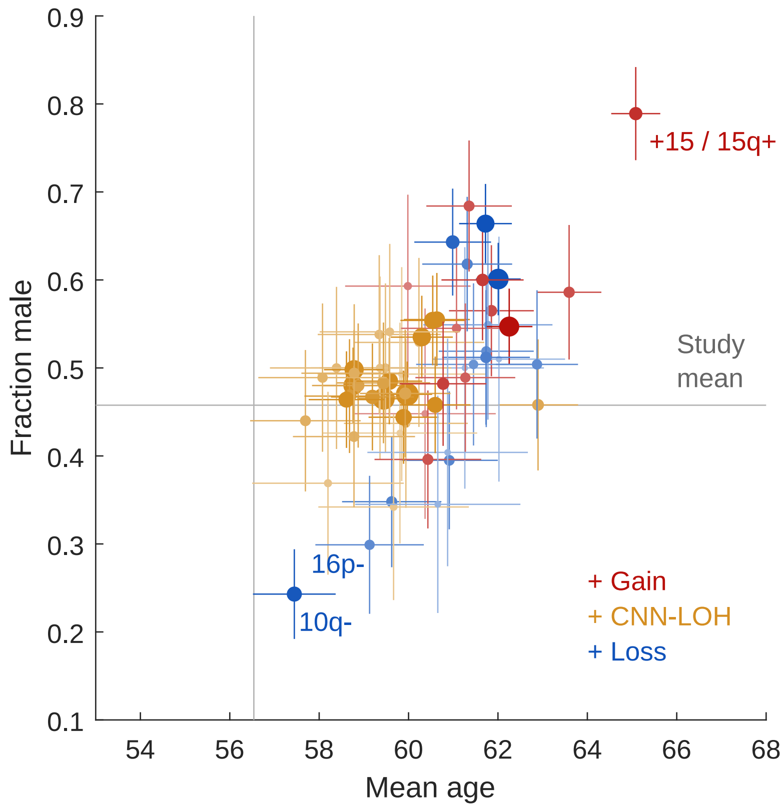

**Supplementary Figure 26. Sex and age distributions of individuals with detected mosaic events.** Carriers of mosaic chromosomal alterations on different chromosome arms and with different copy numbers exhibit different age and sex distributions. Marker size and color intensity increase with event frequency. Error bars, 95% CIs. Numeric data are provided in Supplementary Table 4.

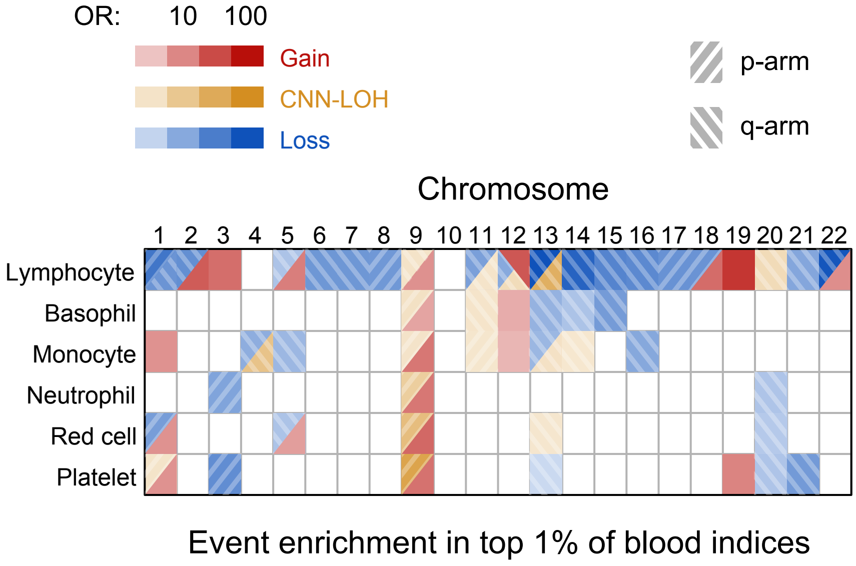

**Supplementary Figure 27. Enrichment of mosaic chromosomal alterations in individuals with anomalously high blood indices.** Different mCAs are significantly enriched (FDR 0.05) among individuals with anomalous blood counts in different blood lineages (adjusted for age, sex, and smoking status). Events were grouped by chromosome and copy number, with loss and CNN-LOH events subdivided by p-arm vs. q-arm. (We did not subdivide gain events by arm because most gain events are whole-chromosome trisomies.) Numeric data are provided in Supplementary Table 5.

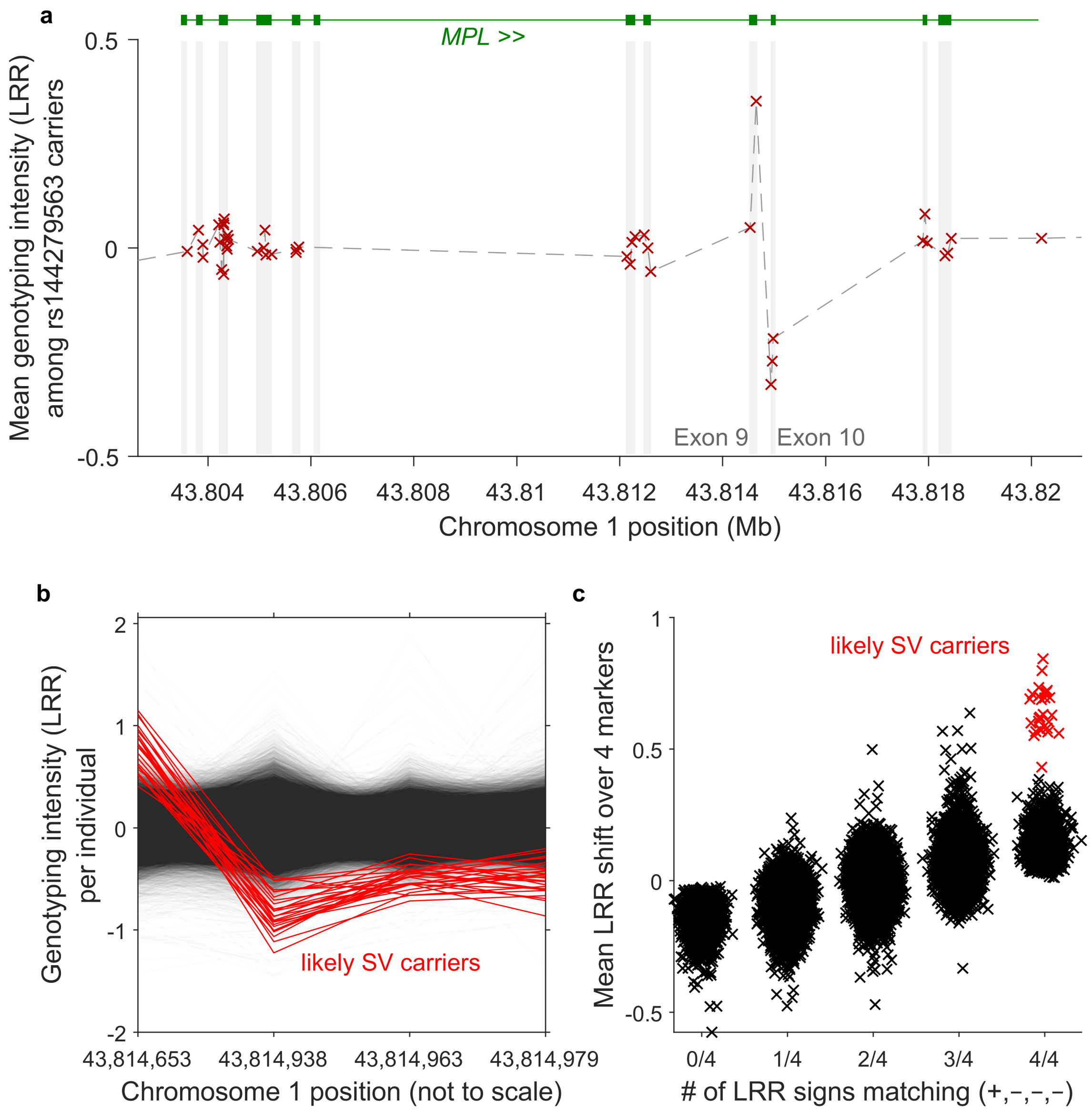

**Supplementary Figure 28. Identification of an inherited *MPL* structural variant from UK BiLEVE genotyping.** We suspected that an association between rs144279563 and acquired 1p CNN-LOH mutations might tag a causal structural variant in *MPL*. (While rs144279563 is ∼1.5Mb downstream from *MPL*, it is sufficiently rare to be in linkage disequilibrium with variants several megabases away.) We therefore examined genotyping intensities at *MPL* from 49,950 individuals typed on the BiLEVE chip (which has more coverage of *MPL* than the Biobank chip, on which the remaining individuals were typed.) (**a**) Mean genotyping intensities over 42 carriers of the rs144279563 rare allele exhibit a sharp increase at the end of *MPL* exon 9 (1 genotyping probe) followed by a sharp decrease in exon 10 (3 genotyping probes). (**b,c**) Closer inspection of genotyping intensities at the 4 probes across all BiLEVE individuals enabled identification of 27 individuals likely to carry an inherited structural variant (20 of which carry the rs144279563 rare allele). We called this variant in the BiLEVE cohort using two criteria: (i) correct sign of LRR at the 4 probes (+,–,–,–); and (ii) mean signed LRR shift >0.4 over the 4 probes.

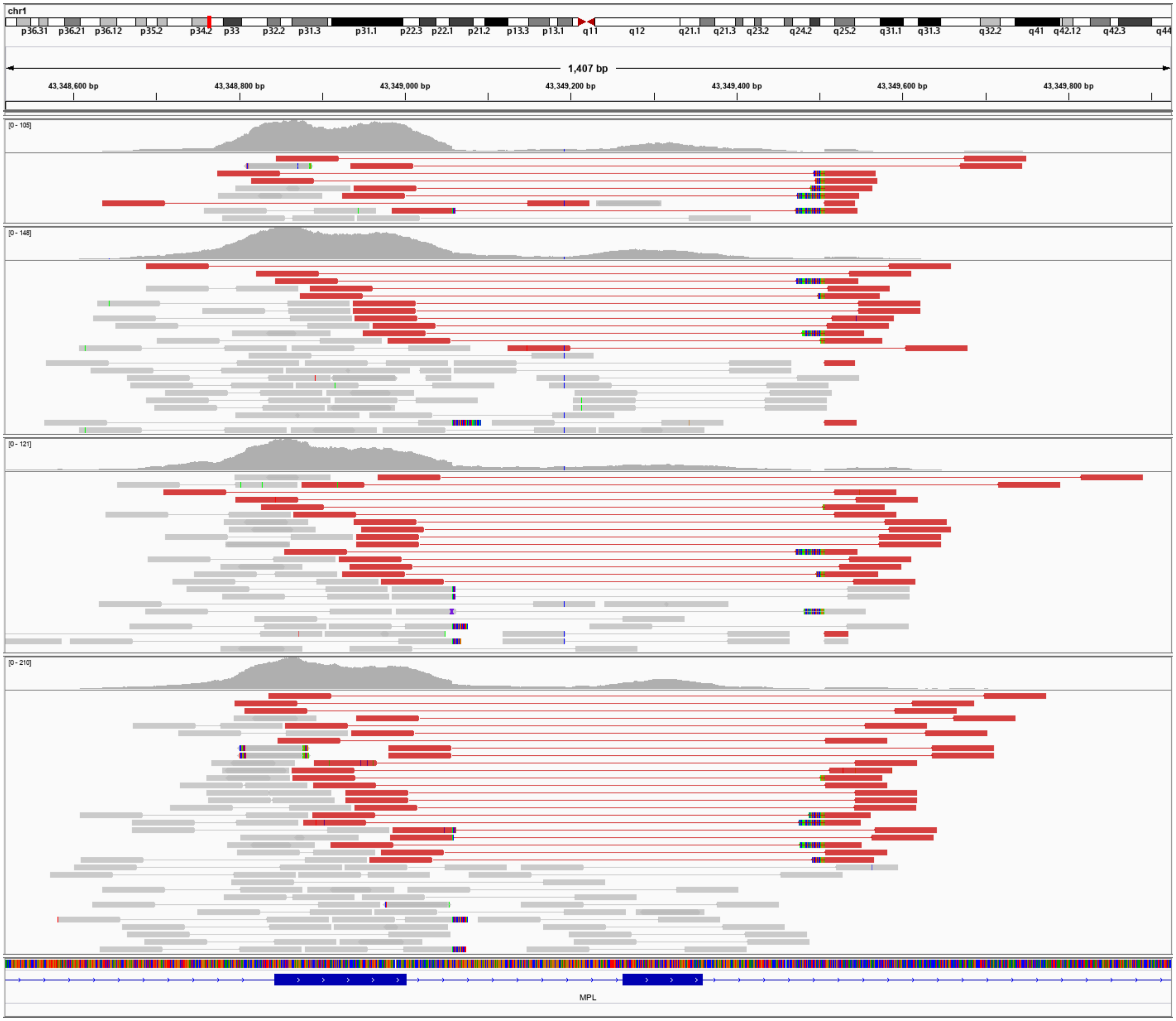

**Supplementary Figure 29. Read support for 454bp deletion spanning *MPL* exon 10 in exome-sequenced individuals.** We used IGV [70] to plot paired-end reads aligning in or near *MPL* exons 9 and 10 in four exome-sequenced individuals predicted to carry an inherited *MPL* structural variant (and also mosaic for 1p CNN-LOH events). Read pairs highlighted in red have unusually long insert sizes, consistent with a deletion of genomic sequence between the aligned reads. Multicolored read segments indicate clipped reads in which one end of a read stops aligning to the reference genome. On the left side of the deletion, clipped reads align right-to-left through hg19 base pair 43,814,728 (. . . AGGGA**CTGGG**), with mismatches consistently occurring starting from 43,814,729 rightward (CGCCG. . . ). On the right side of the deletion, clipped reads align left-to-right until 43,815,178 (**CTGGG**ACTCG. . . ), with mismatches from starting from 43,815,177 leftward (. . . CACCT). Examination of individual clipped reads revealed sequence matching . . . AGGGA**CTGGG**ACTCG. . . , indicating deletion of 5bp (**CTGGG**) in addition to the 449bp between aligning read segments. (Note that in this caption we have used hg19 coordinates for consistency with the rest of this manuscript; the IGV plot above uses hg38 coordinates because reads had been aligned to hg38 (amounting to an offset of –465,671 bases relative to hg19 at *MPL*).

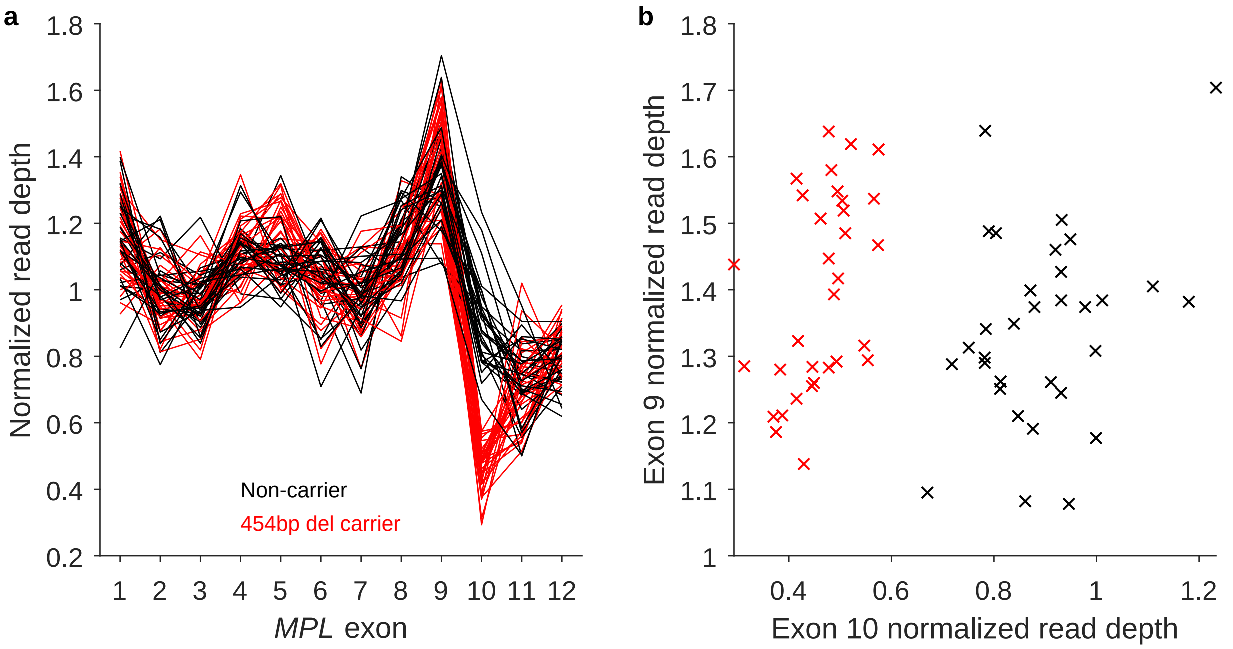

**Supplementary Figure 30. Decreased read depth at exon 10 in all 32 predicted carriers of inherited *MPL* deletion who had been exome-sequenced.** We used mosdepth [71] to compute mean read depth across all 12 *MPL* exons in 32 exome-sequenced individuals we had imputed to be carriers of the *MPL* exon 10 deletion along with 32 exome-sequenced control individuals. We normalized read depth in each individual by dividing by mean read depth across exons 1–8 and 11–12. All 32 imputed carriers of the exon 10 deletion had lower exon 10 normalized read depths than all 32 controls. We did not observe any evidence of increased read depth in exon 9 in carriers vs. controls.

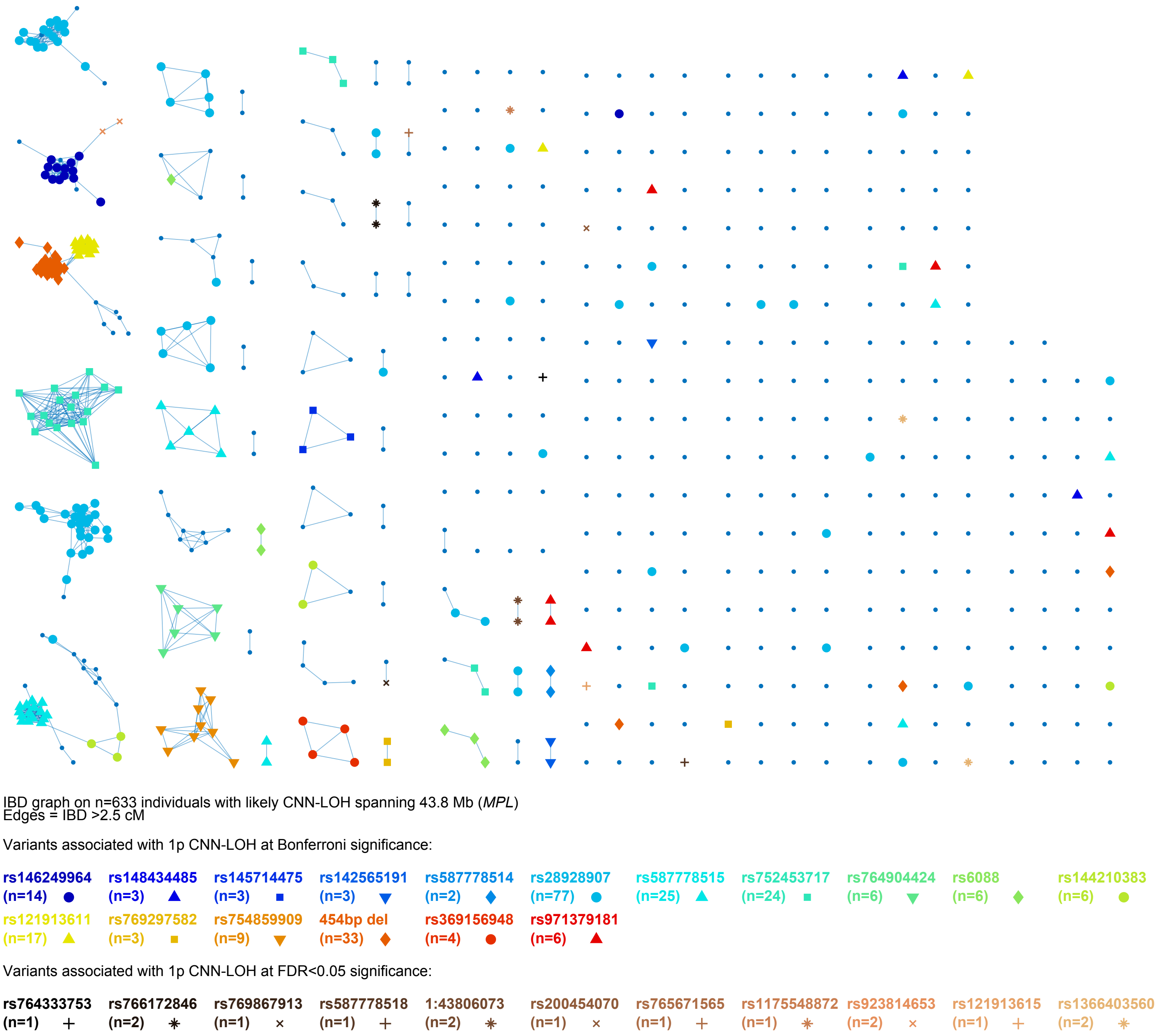

**Supplementary Figure 31. Identity-by-descent (IBD) graph at *MPL* among individuals with likley 1p CNN-LOH events spanning *MPL*.** We called IBD tracts using GERMLINE with haplotype extension [72]. Colored nodes indicate carriers of the 28 rare coding or splice variants we observed to be independently (and probably causally) associated with 1p CNN-LOH mutations (always replacing the rare allele with the reference allele; Table 1 and Supplementary Table 6). (Note that the numbers of carriers listed for each variant here are slightly higher than in the "Allelic shift" columns of Table 1 and Supplementary Table 6 because allelic shifts could only be confidently ascertained for a subset of carriers.) The presence of additional IBD clusters not carrying any of the 28 highlighted variants suggests that even more causal variants in *MPL* remain to be discovered.

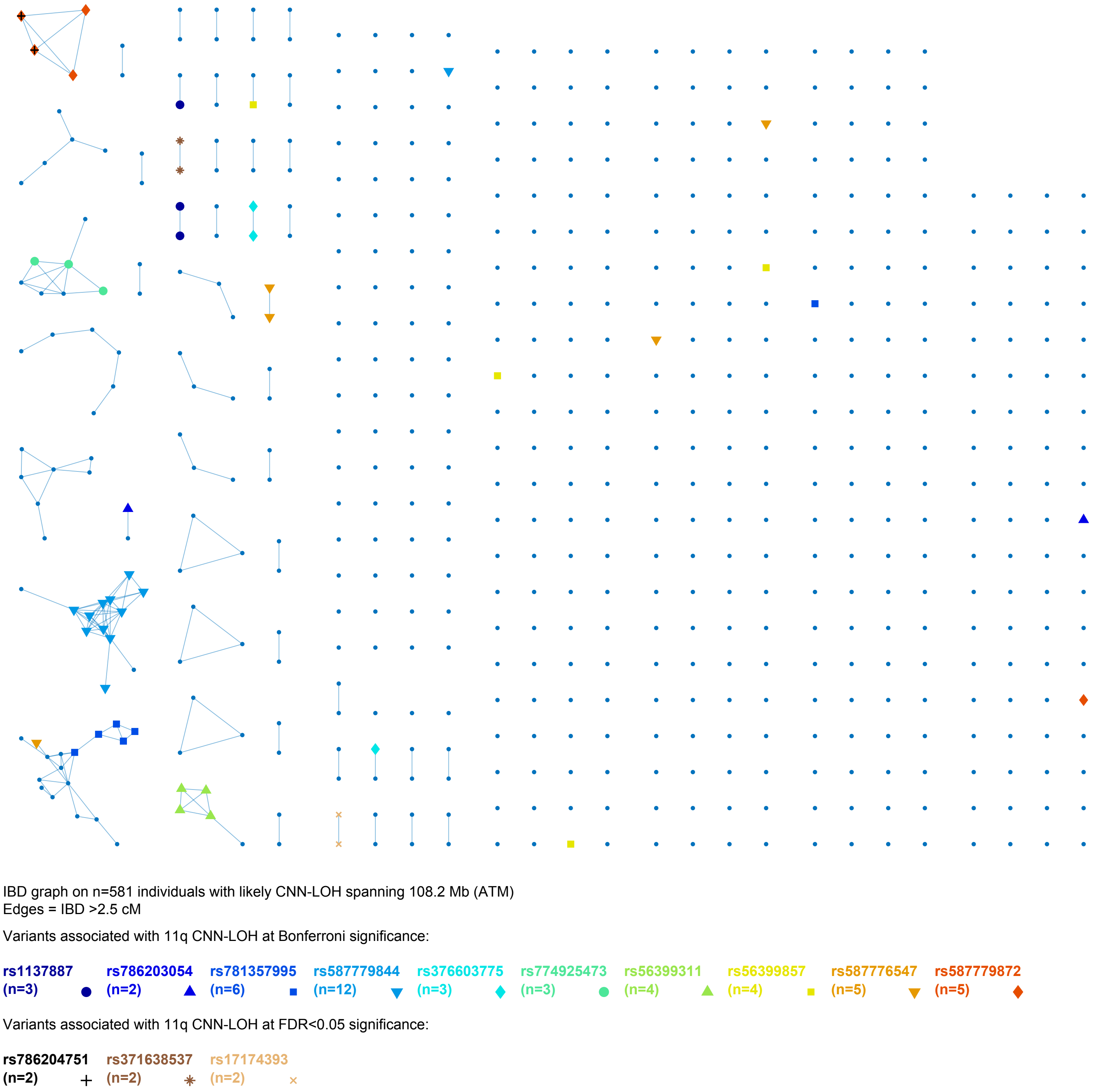

**Supplementary Figure 32. Identity-by-descent (IBD) graph at *ATM* among individuals with likely 11q CNN-LOH events spanning *ATM*.** We called IBD tracts using GERMLINE with haplotype extension [72]. Colored nodes indicate carriers of the eight rare coding or splice variants we observed to be independently (and probably causally) associated with 11q CNN-LOH mutations (always duplicating the rare allele; Table 1 and Supplementary Table 6). The presence of additional IBD clusters not carrying any of the highlighted variants suggests that even more causal variants in *ATM* remain to be discovered. The two carriers of rs786204751 are also carriers of rs587779872, as discussed in Methods.

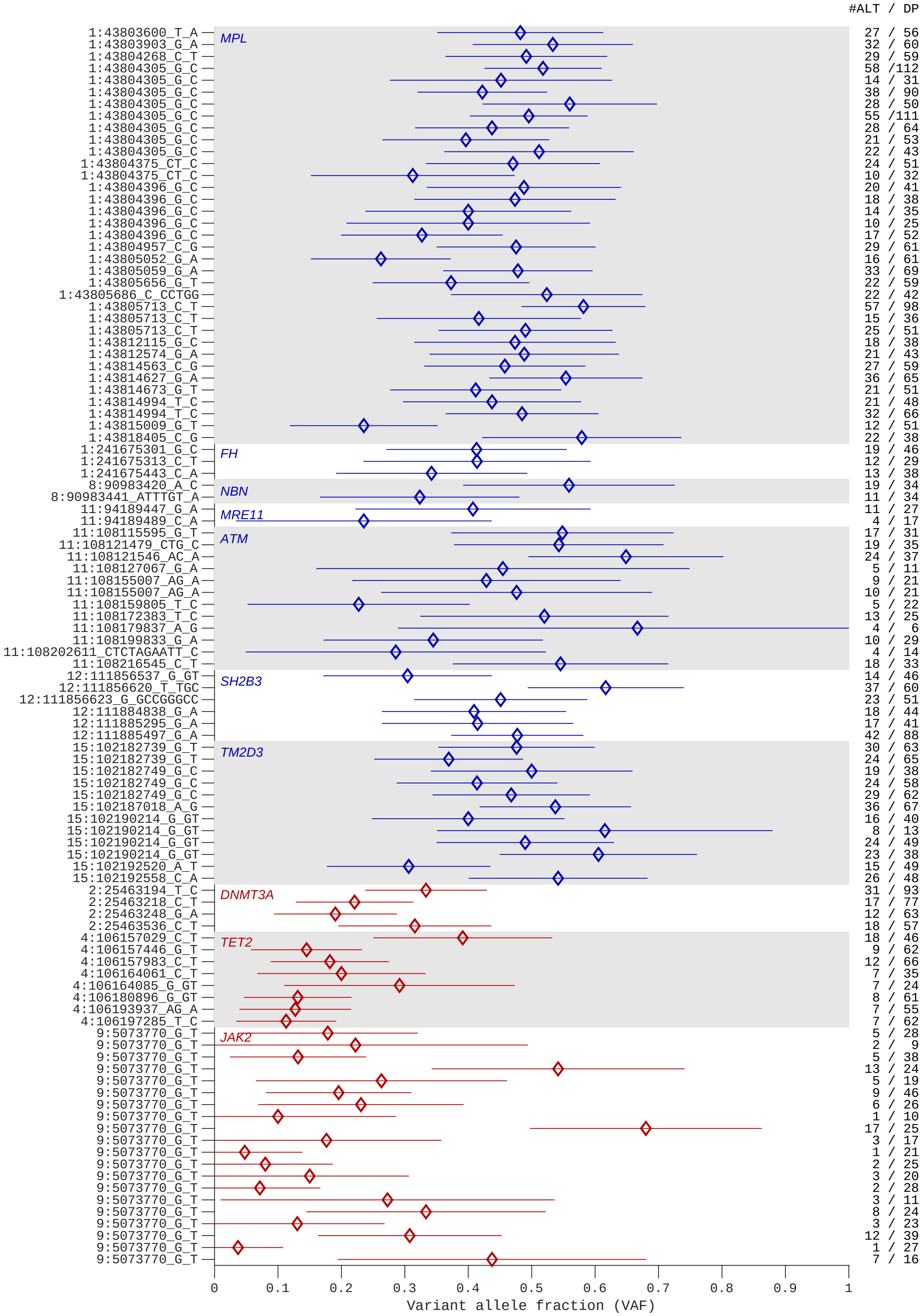

**Supplementary Figure 33. Variant allele fractions of rare coding or splice variants likely to be targets of CNN-LOH mutations in exome-sequenced individuals.** See next page for caption.

**Caption for Supplementary Fig 33.**

Variant allele fractions (VAF = number of reads matching the alternate allele divided by the total number of reads matching either the reference or the alternate allele) are plotted for each variant call identified as the potential target of a CNN-LOH event (either from our association analyses or burden analyses). Error bars, 95% CIs estimated as binomial standard error multiplied by 1.96.

Allelic read depths for variants identified at *DNMT3A*, *TET2*, and *JAK2* are broadly indicative of somatic origin (VAF$<$0.5), while read depths for variants at the seven inherited risk loci are broadly consistent with inherited variation (VAF$\approx$0.5). Read depths were generally insufficient to make a confident assessment of somatic vs. inherited origin on a per-variant level, as evidenced by wide VAF error bars; additionally, making this determination is further complicated by mapping bias toward the reference allele, which can produce VAF lower than 0.5 even for inherited variants [4].

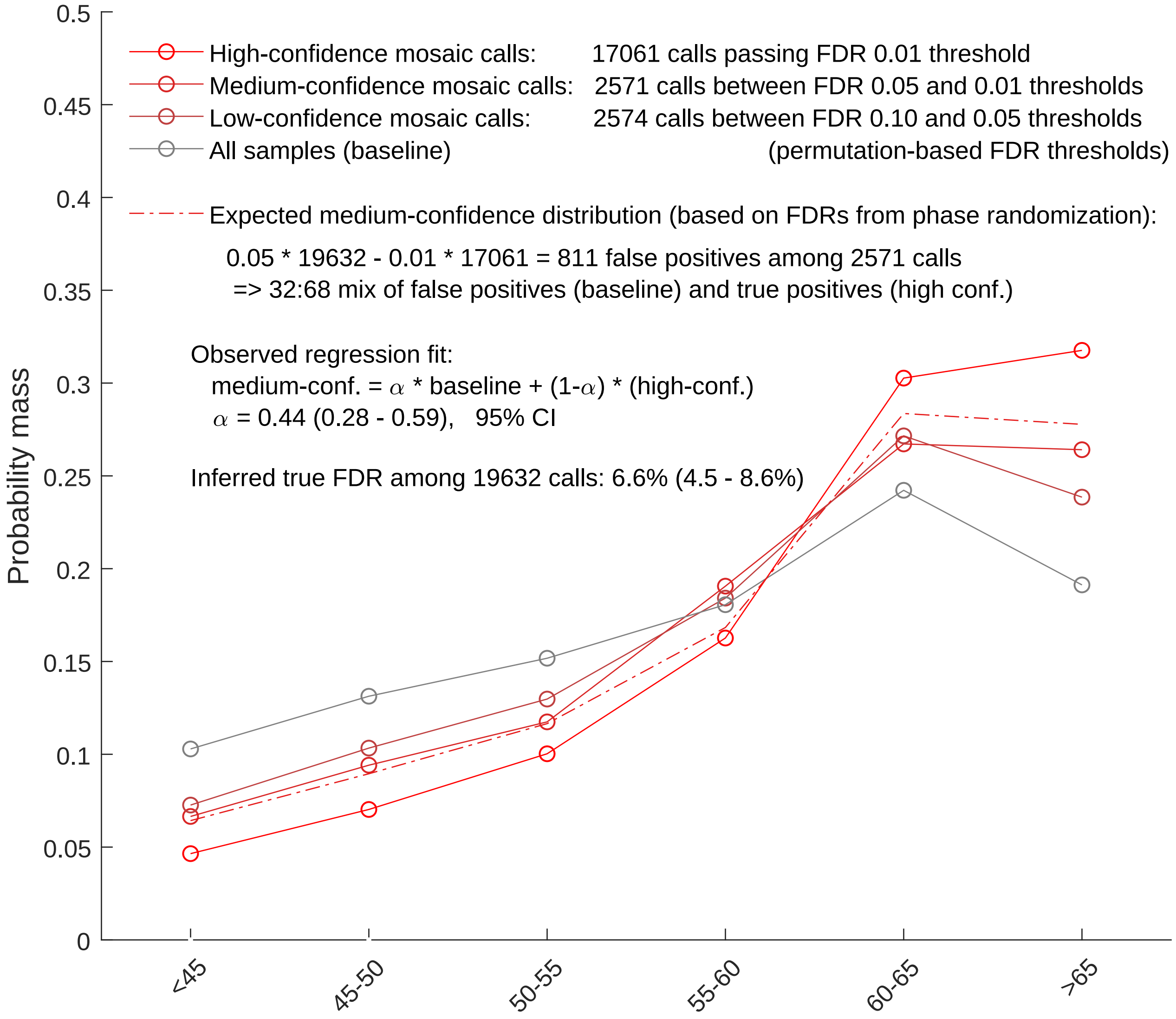

**Supplementary Figure 34. Estimation of true FDR using age distributions of individuals with mCA calls.** We generated age distributions for (i) "high-confidence" detected events passing a permutation-based FDR threshold of 0.01 (bright red), (ii) "medium-confidence" events below the FDR threshold of 0.01 but passing an FDR threshold of 0.05 (darker red), and (iii) "low-confidence" events below the FDR threshold of 0.05 but passing an FDR threshold of 0.10 (darkest red; not analyzed but plotted for context). We compared these distributions to the overall age distribution of UK Biobank participants (grey). Based on the numbers of events in each category, ≈32% of medium-confidence detected events are expected to be false positives. To estimate our true FDR, we regressed the medium-confidence age distribution on the high-confidence and overall age distributions, reasoning that the medium-confidence age distribution should be a mixture of (a) correctly-called events with age distribution similar to that of the high-confidence events and (b) spurious calls with age distribution similar to the overall cohort. We observed a regression weight of 0.44 for the component corresponding to spurious calls, in good agreement with expectation, and implying a true FDR of 6.6% (4.5–8.6%, 95% CI based on regression fit on $n$=6 age bins).

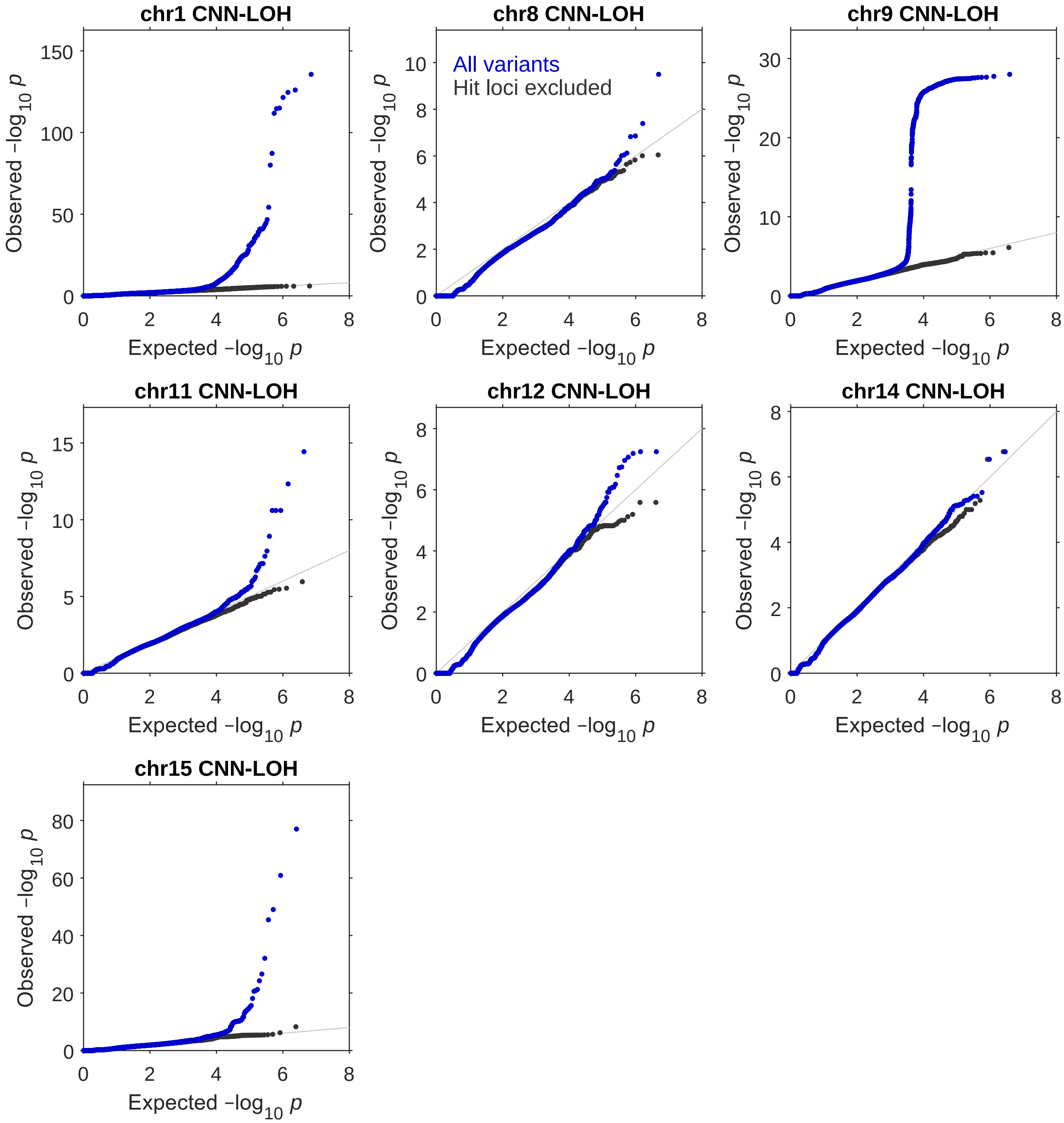

**Supplementary Figure 35. Quantile-quantile plots of *P*-values produced by association analyses.** These plots verify the calibration of the statistical tests we used to identify the genome-wide significant associations reported in Table 1. In each plot, the blue dots correspond to an analysis of all variants tested, while the black dots correspond to an analysis in which regions surrounding significant associations were excluded. Specifically, the plots respectively exclude 1:35-55Mb (*MPL*), 1:239–244Mb (*FH*), 8:88–93Mb (*NBN*), 9:2.5–7.5Mb (*JAK2*), 11:92–97Mb (*MRE11*), 11:103–108Mb (*ATM*), 12:109–114Mb (*SH2B3*), 14:92.5–102.5Mb (*TCL1A* and *DLK1*), and 15:100Mb–qter (*TM2D3*). In all cases, exclusion of the hit regions (which account for a small fraction of the variants tested) results in a distribution close to the expected null.

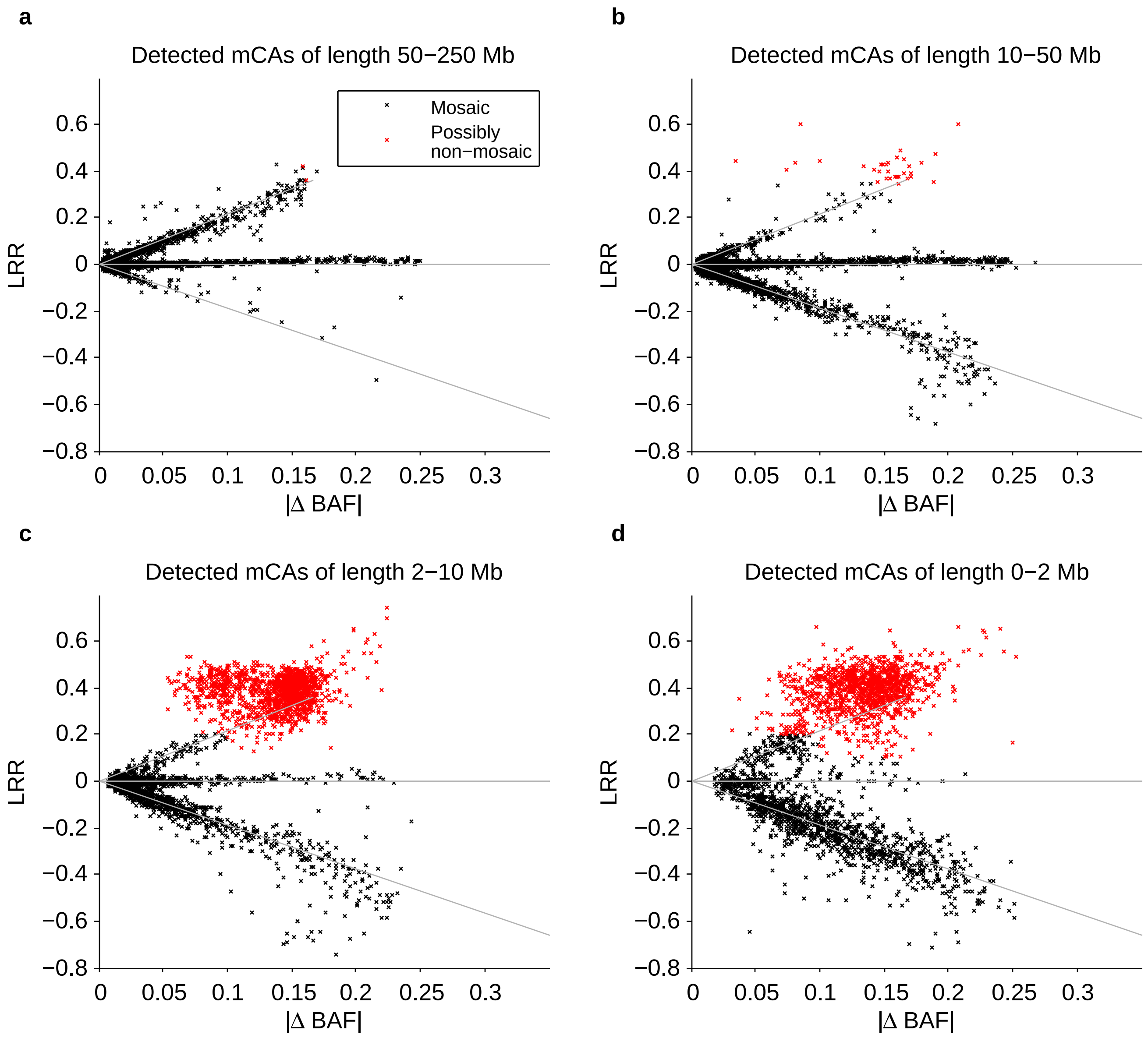

**Supplementary Figure 36. Exclusion of possible constitutional duplications.** We filtered subchromosomal events of length >10Mb with LRR>0.35 or with LRR>0.2 and $|\Delta\text{BAF}|>0.16$, and we filtered events of length <10Mb with LRR>0.2 or with LRR>0.1 and $|\Delta\text{BAF}|>0.1$. Constitutional duplications have expected $|\Delta\text{BAF}|=1/6$, corresponding to LRR of roughly 0.36. The bottom two panels (corresponding to event calls 2–10Mb and <2Mb) each clearly contain a cluster of calls around $|\Delta\text{BAF}|=1/6$, LRR=0.36. We chose exclusion thresholds to conservatively discard all calls that might belong to this cluster, applying more stringent filtering to shorter events because (i) most constitutional duplications are short and (ii) shorter events have noisier LRR and $|\Delta\text{BAF}|$ estimates.

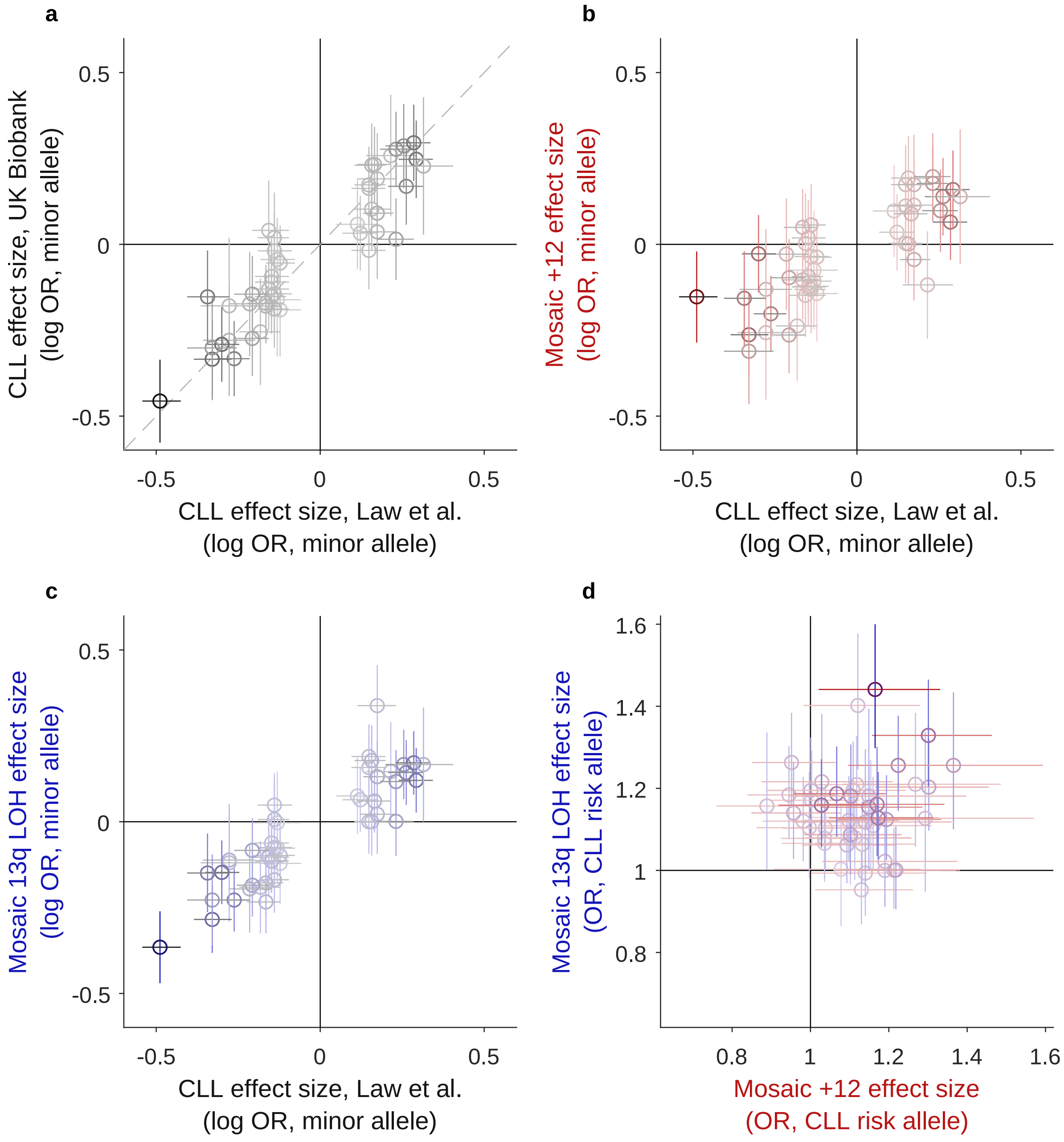

**Supplementary Figure 37. Effects of known CLL GWAS variants on mosaic +12 and 13q LOH risk.** For 46 lead CLL-associated variants reported by Law et al. [60], we plotted (**a**) CLL effect size, (**b**) mosaic +12 effect size, and (**c**) mosaic 13q LOH (i.e., del(13q) or 13q CNN-LOH) effect size in UK Biobank vs. reported CLL effect size in Law et al. [60]. We also plotted mosaic 13q LOH effect size vs. mosaic +12 effect size for the same 46 variants (**d**). Marker intensity is proportional to CLL association strength. Error bars, 95% CIs. Numeric data are provided in Supplementary Table 20, and details of analyses are provided in the Supplementary Note.

**Supplementary Table 1. Number of mosaic chromosomal alterations detected per chromosome.**

| Chromosome | $N_{\text{loss}}$ | $N_{\text{CNN-LOH}}$ | $N_{\text{gain}}$ | $N_{\text{undetermined}}$ | $N_{\text{total}}$ |
|---|---|---|---|---|---|
| chr1 | 97 | 1179 | 68 | 522 | 1866 |
| chr2 | 214 | 264 | 26 | 159 | 663 |
| chr3 | 74 | 225 | 174 | 185 | 658 |
| chr4 | 138 | 245 | 24 | 116 | 523 |
| chr5 | 172 | 131 | 88 | 110 | 501 |
| chr6 | 119 | 311 | 33 | 194 | 657 |
| chr7 | 176 | 176 | 23 | 118 | 493 |
| chr8 | 69 | 127 | 156 | 141 | 493 |
| chr9 | 52 | 706 | 112 | 258 | 1128 |
| chr10 | 290 | 129 | 11 | 131 | 561 |
| chr11 | 296 | 875 | 4 | 366 | 1541 |
| chr12 | 81 | 251 | 530 | 342 | 1204 |
| chr13 | 596 | 444 | 12 | 237 | 1289 |
| chr14 | 161* | 704 | 162 | 377 | 1404 |
| chr15 | 44 | 407 | 223 | 332 | 1006 |
| chr16 | 180 | 470 | 8 | 202 | 860 |
| chr17 | 231 | 435 | 135 | 290 | 1091 |
| chr18 | 58 | 100 | 200 | 153 | 511 |
| chr19 | 17 | 339 | 48 | 275 | 679 |
| chr20 | 458 | 204 | 8 | 133 | 803 |
| chr21 | 58 | 138 | 153 | 228 | 577 |
| chr22 | 137* | 325 | 191 | 471 | 1124 |
| All autosomes | 3718 | 8185 | 2389 | 5340 | 19632 |

*Deletions on chr14 and chr22 include V(D)J recombination events (61 events on chr14 and 80 events on chr22).

**Supplementary Table 2. Distribution of the number of detected somatic autosomal mCAs per individual.**

| mCA count | Frequency |
|---:|---:|
| 0 | 465678 |
| 1 | 15520 |
| 2 | 1084 |
| 3 | 329 |
| 4 | 95 |
| 5 | 40 |
| 6 | 18 |
| 7 | 6 |
| 8 | 6 |
| 9 | 1 |
| 10 | 3 |
| 11 | 0 |
| 12 | 1 |
| 13 | 1 |
| 14 | 2 |
| 15 | 1 |
| 16 | 1 |
| 17 | 2 |
| 18 | 0 |
| 19 | 0 |
| 20 | 0 |
| 21 | 0 |
| 22 | 1 |

Most individuals with several detected mCAs have prevalent or incident blood cancers.

**Supplementary Table 3. Fraction of individuals with detected mCAs as a function of age.**

| Age range | % of males with autosomal event (s.e.) | % of females with autosomal event (s.e.) |
|---|---|---|
| $<$45 | 1.8% (0.1%) | 1.8% (0.1%) |
| 45-50 | 2.1% (0.1%) | 2.0% (0.1%) |
| 50-55 | 2.6% (0.1%) | 2.4% (0.1%) |
| 55-60 | 3.5% (0.1%) | 3.1% (0.1%) |
| 60-65 | 4.7% (0.1%) | 4.0% (0.1%) |
| $>$65 | 6.0% (0.1%) | 4.9% (0.1%) |

Consistent with previous work [2, 3, 6, 7, 10], mosaic chromosomal alterations are detected more frequently with increasing age and in males.

**Supplementary Table 4. Age and sex distributions of individuals with detected mCAs on each chromosome.**

| | Loss events | | | | CNN-LOH events | | | | Gain events | |
|---|---|---|---|---|---|---|---|---|---|---|
| | p-arm | | q-arm | | p-arm | | q-arm | | | |
| chr | Mean age | Frac. male | Mean age | Frac. male | Mean age | Frac. male | Mean age | Frac. male | Mean age | Frac. male |
| 1 | 60.6 (1.1) | 0.40 (0.08) | 61.2 (0.9) | 0.62 (0.08) | 59.4 (0.3) | 0.46 (0.02) | 58.8 (0.4) | 0.48 (0.02) | 60.4 (0.8) | 0.45 (0.06) |
| 2 | 60.9 (0.6) | 0.39 (0.04) | 61.3 (0.9) | 0.50 (0.07) | 59.9 (0.7) | 0.44 (0.05) | 57.7 (0.6) | 0.44 (0.04) | 58.6 (1.4) | 0.54 (0.10) |
| 3 | 60.9 (1.2) | 0.57 (0.08) | 61.5 (1.6) | 0.36 (0.10) | 59.3 (0.7) | 0.54 (0.05) | 60.2 (0.7) | 0.53 (0.05) | 61.9 (0.5) | 0.56 (0.04) |
| 4 | 63.5 (1.1) | 0.25 (0.13) | 61.5 (0.7) | 0.50 (0.05) | 55.5 (1.3) | 0.49 (0.08) | 62.9 (0.5) | 0.46 (0.04) | 61.2 (1.9) | 0.52 (0.11) |
| 5 | 62.0 (2.7) | 0.40 (0.16) | 59.6 (0.6) | 0.35 (0.04) | 57.9 (1.7) | 0.57 (0.14) | 58.4 (0.8) | 0.50 (0.05) | 60.0 (0.7) | 0.59 (0.05) |
| 6 | 61.8 (1.3) | 0.39 (0.09) | 61.8 (0.7) | 0.55 (0.06) | 58.7 (0.5) | 0.47 (0.03) | 59.8 (1.0) | 0.49 (0.06) | 59.3 (1.5) | 0.55 (0.09) |
| 7 | 59.5 (1.2) | 0.30 (0.08) | 61.7 (0.5) | 0.52 (0.04) | 59.8 (0.9) | 0.43 (0.06) | 59.5 (0.8) | 0.50 (0.05) | 59.0 (1.8) | 0.43 (0.11) |
| 8 | 62.0 (0.8) | 0.51 (0.07) | 62.0 (1.3) | 0.56 (0.13) | 56.3 (1.4) | 0.41 (0.09) | 59.4 (0.8) | 0.50 (0.05) | 60.4 (0.6) | 0.40 (0.04) |
| 9 | 66.6 (1.1) | 0.43 (0.20) | 61.1 (1.4) | 0.47 (0.08) | 60.6 (0.4) | 0.55 (0.03) | 59.9 (0.4) | 0.44 (0.03) | 61.1 (0.6) | 0.54 (0.05) |
| 10 | 63.7 (2.1) | 0.50 (0.22) | 57.4 (0.5) | 0.24 (0.03) | 58.5 (1.3) | 0.51 (0.08) | 58.2 (0.9) | 0.37 (0.05) | 58.8 (2.8) | 0.36 (0.15) |
| 11 | 59.0 (1.3) | 0.57 (0.08) | 61.0 (0.4) | 0.64 (0.03) | 58.8 (0.3) | 0.50 (0.02) | 60.5 (0.4) | 0.55 (0.03) | – | – |
| 12 | 62.3 (1.1) | 0.57 (0.09) | 60.7 (1.1) | 0.52 (0.08) | 57.8 (1.2) | 0.34 (0.07) | 58.9 (0.5) | 0.48 (0.04) | 62.3 (0.3) | 0.55 (0.02) |
| 13 | – | – | 62.0 (0.3) | 0.60 (0.02) | – | – | 60.3 (0.4) | 0.54 (0.02) | 56.7 (2.9) | 0.75 (0.13) |
| 14 | – | – | 61.3 (0.5) | 0.62 (0.04) | – | – | 60.0 (0.3) | 0.47 (0.02) | 63.6 (0.4) | 0.59 (0.04) |
| 15 | – | – | 62.2 (1.1) | 0.46 (0.08) | – | – | 59.6 (0.4) | 0.48 (0.03) | 65.1 (0.3) | 0.79 (0.03) |
| 16 | 59.1 (0.6) | 0.30 (0.04) | 61.7 (1.1) | 0.61 (0.08) | 59.2 (0.5) | 0.47 (0.03) | 59.4 (0.5) | 0.48 (0.03) | 55.0 (3.8) | 0.50 (0.19) |
| 17 | 61.7 (0.5) | 0.51 (0.04) | 60.9 (0.9) | 0.40 (0.07) | 59.6 (0.8) | 0.54 (0.05) | 58.6 (0.4) | 0.46 (0.03) | 61.3 (0.6) | 0.49 (0.04) |
| 18 | 60.6 (1.1) | 0.67 (0.09) | 61.5 (1.6) | 0.32 (0.10) | 60.3 (1.8) | 0.71 (0.13) | 59.7 (0.9) | 0.34 (0.05) | 61.7 (0.5) | 0.60 (0.03) |
| 19 | 56.8 (2.3) | 0.70 (0.15) | 61.7 (2.5) | 0.71 (0.18) | 58.8 (0.7) | 0.42 (0.04) | 59.9 (0.5) | 0.47 (0.04) | 59.2 (1.1) | 0.69 (0.07) |
| 20 | 62.8 (1.4) | 0.50 (0.15) | 61.7 (0.3) | 0.66 (0.02) | 57.4 (1.3) | 0.44 (0.08) | 58.8 (0.6) | 0.49 (0.04) | 57.4 (1.3) | 0.25 (0.16) |
| 21 | – | – | 60.7 (0.9) | 0.34 (0.06) | – | – | 58.1 (0.7) | 0.49 (0.04) | 61.4 (0.5) | 0.68 (0.04) |
| 22 | – | – | 62.9 (0.5) | 0.50 (0.04) | – | – | 60.6 (0.4) | 0.46 (0.03) | 60.8 (0.5) | 0.48 (0.04) |

This table provides numerical data plotted in Supplementary Fig. 26. (Events detected in fewer than 50 individuals were excluded from Supplementary Fig. 26 for clarity, and events detected in fewer than 5 individuals are excluded here. Chromosomes 13, 14, 15, 21, and 22 are acrocentric and have little or no p-arm genotyping.)

**Supplementary Table 5. Enrichment of mCAs in individuals with anomalous (top 1%) blood indices.**

| mCA | Blood index | *P*-value | *q*-value | OR (95% CI) |
|---|---|---|---|---|
| 1p– | Lymphocyte # | 1.5e-7 | 3.4e-6 | 31.3 (12.5–78.4) |
| 1p– | Lymphocyte % | 4.1e-9 | 1.1e-7 | 38.6 (16.1–92.4) |
| 1p– | RBC dist. width | 0.0019 | 0.022 | 13.5 (4.0–45.2) |
| 1q– | Lymphocyte # | 0.00015 | 0.0021 | 17.2 (6.0–49.9) |
| 1q– | Lymphocyte % | 0.00015 | 0.0021 | 17.2 (6.0–49.9) |
| 1p= | Platelet # | 8.9e-5 | 0.0014 | 3.0 (1.9–4.9) |
| 1p= | Platelet crit | 8.9e-5 | 0.0014 | 3.0 (1.9–4.9) |
| 1+ | Monocyte # | 0.002 | 0.022 | 8.1 (2.9–22.4) |
| 1+ | Monocyte % | 0.002 | 0.022 | 8.1 (2.9–22.4) |
| 1+ | RBC dist. width | 0.002 | 0.022 | 8.1 (2.9–22.4) |
| 1+ | Platelet dist. width | 0.002 | 0.022 | 8.1 (2.9–22.4) |
| 2q– | Lymphocyte # | 2e-7 | 4.4e-6 | 19.8 (8.8–44.7) |
| 2q– | Lymphocyte % | 3.8e-6 | 7.8e-5 | 16.5 (7.0–39.2) |
| 2+ | Lymphocyte # | 0.00051 | 0.0066 | 22.9 (6.5–80.3) |
| 3p– | Neutrophil % | 0.003 | 0.031 | 11.4 (3.5–37.8) |
| 3p– | Platelet # | 9.7e-6 | 0.00017 | 20.6 (7.9–54.1) |
| 3p– | Platelet crit | 0.00019 | 0.0028 | 15.9 (5.5–45.6) |
| 3+ | Lymphocyte # | 1.9e-12 | 7.4e-11 | 16.2 (9.2–28.5) |
| 3+ | Lymphocyte % | 6.2e-9 | 1.7e-7 | 12.3 (6.6–23.0) |
| 4q– | Monocyte # | 6.2e-5 | 0.00098 | 7.6 (3.5–16.5) |
| 4q– | Monocyte % | 7e-6 | 0.00013 | 8.8 (4.3–18.2) |
| 4q= | Monocyte # | 0.00025 | 0.0036 | 5.2 (2.5–10.5) |
| 4q= | Monocyte % | 1.1e-12 | 4.2e-11 | 11.6 (7.0–19.3) |
| 5q– | Lymphocyte # | 0.0018 | 0.021 | 5.0 (2.2–11.3) |
| 5q– | Monocyte % | 4.4e-5 | 0.0007 | 6.7 (3.3–13.8) |
| 5q– | RBC dist. width | 0.0018 | 0.021 | 5.0 (2.2–11.3) |
| 5+ | Lymphocyte # | 4.6e-6 | 9e-5 | 11.8 (5.4–25.8) |
| 5+ | RBC dist. width | 0.0044 | 0.043 | 6.4 (2.3–17.6) |
| 6q– | Lymphocyte # | 4.4e-6 | 8.8e-5 | 16.1 (6.8–38.1) |
| 6q– | Lymphocyte % | 0.0009 | 0.012 | 10.2 (3.6–28.5) |
| 7q– | Lymphocyte # | 7.6e-11 | 2.7e-9 | 16.1 (8.7–29.6) |
| 7q– | Lymphocyte % | 1.2e-9 | 3.9e-8 | 14.6 (7.7–27.4) |
| 8p– | Lymphocyte # | 0.00051 | 0.0066 | 12.0 (4.3–33.9) |
| 8p– | Lymphocyte % | 3.3e-5 | 0.00055 | 15.5 (6.0–39.8) |
| 9p= | Basophil # | 0.001 | 0.013 | 3.4 (1.8–6.4) |
| 9p= | Monocyte # | 0.0036 | 0.037 | 3.1 (1.6–6.0) |
| 9p= | Neutrophil # | 6.7e-12 | 2.5e-10 | 7.5 (4.8–11.7) |
| 9p= | Neutrophil % | 3.6e-10 | 1.2e-8 | 6.7 (4.2–10.7) |
| 9p= | Red # | 6.9e-24 | 5.3e-22 | 12.3 (8.6–17.7) |
| 9p= | Hematocrit | 1.3e-15 | 6.7e-14 | 9.0 (6.0–13.6) |
| 9p= | RBC dist. width | 1.5e-29 | 1.2e-27 | 14.5 (10.3–20.4) |
| 9p= | Platelet # | 3.2e-97 | 1.1e-94 | 40.6 (31.5–52.2) |
| 9p= | Platelet crit | 5.1e-94 | 1.4e-91 | 39.2 (30.4–50.6) |
| 9p= | Platelet dist. width | 1e-13 | 4.8e-12 | 8.3 (5.4–12.6) |
| 9q= | Lymphocyte # | 0.0029 | 0.031 | 3.2 (1.6–6.1) |
| 9+ | Lymphocyte # | 3.9e-5 | 0.00063 | 8.3 (3.8–17.9) |
| 9+ | Basophil # | 0.0023 | 0.025 | 5.8 (2.3–14.2) |
| 9+ | Monocyte # | 2.2e-9 | 6.3e-8 | 13.6 (7.3–25.6) |
| 9+ | Neutrophil # | 3e-8 | 7.3e-7 | 12.2 (6.3–23.6) |
| 9+ | Neutrophil % | 2.2e-9 | 6.3e-8 | 13.6 (7.3–25.6) |
| 9+ | RBC dist. width | 5.1e-13 | 2.1e-11 | 18.1 (10.2–31.9) |
| 9+ | Platelet # | 1.5e-10 | 5.2e-9 | 15.1 (8.2–27.7) |
| 9+ | Platelet crit | 2.2e-9 | 6.3e-8 | 13.6 (7.3–25.6) |
| 9+ | Platelet dist. width | 2.2e-9 | 6.3e-8 | 13.6 (7.3–25.6) |
| 11q– | Lymphocyte # | 7.3e-17 | 4.3e-15 | 12.9 (8.3–20.2) |
| 11q– | Lymphocyte % | 1.3e-9 | 4.2e-8 | 8.5 (5.0–14.4) |
| 11p= | Monocyte % | 0.0017 | 0.021 | 2.8 (1.6–5.0) |
| 11q= | Lymphocyte % | 0.0024 | 0.026 | 3.0 (1.6–5.7) |
| 11q= | Basophil # | 0.0024 | 0.026 | 3.0 (1.6–5.7) |
| 12q– | Lymphocyte % | 0.0044 | 0.043 | 9.9 (3.0–32.5) |
| 12q= | Lymphocyte # | 0.0019 | 0.022 | 4.2 (2.0–9.0) |
| 12+ | Lymphocyte # | 1.6e-77 | 3.4e-75 | 25.0 (19.6–32.0) |
| 12+ | Lymphocyte % | 1.3e-58 | 1.6e-56 | 19.8 (15.2–25.8) |
| 12+ | Basophil # | 5e-8 | 1.2e-6 | 4.9 (3.1–7.7) |
| 12+ | Monocyte # | 5e-6 | 9.6e-5 | 4.1 (2.5–6.7) |
| 13q– | Lymphocyte # | 1.9e-293 | 2.1e-290 | 106.9 (87.8–130.3) |
| 13q– | Lymphocyte % | 3.6e-242 | 1.9e-239 | 82.1 (67.3–100.1) |
| 13q– | Basophil # | 3.8e-16 | 2e-14 | 7.7 (5.3–11.2) |
| 13q– | Monocyte # | 1.6e-13 | 7e-12 | 6.8 (4.6–10.2) |
| 13q– | Platelet dist. width | 0.0029 | 0.031 | 2.8 (1.5–5.1) |
| 13q= | Lymphocyte # | 8.2e-73 | 1.5e-70 | 25.9 (20.0–33.4) |
| 13q= | Lymphocyte % | 3.2e-67 | 5e-65 | 24.1 (18.6–31.3) |
| 13q= | Monocyte # | 0.0004 | 0.0055 | 3.4 (1.9–6.0) |
| 13q= | RBC dist. width | 0.0046 | 0.044 | 2.8 (1.5–5.2) |
| 14q– | Lymphocyte # | 1.8e-53 | 1.8e-51 | 63.0 (42.6–93.3) |
| 14q– | Lymphocyte % | 1.8e-53 | 1.8e-51 | 63.0 (42.6–93.3) |
| 14q– | Basophil # | 0.0044 | 0.043 | 4.9 (2.0–12.0) |
| 14q= | Monocyte % | 0.00049 | 0.0066 | 2.7 (1.6–4.4) |
| 15q– | Lymphocyte # | 3.6e-6 | 7.5e-5 | 26.1 (9.7–69.9) |
| 15q– | Lymphocyte % | 9e-5 | 0.0014 | 19.8 (6.8–58.0) |
| 15q– | Basophil # | 0.0017 | 0.021 | 14.2 (4.2–47.5) |
| 16q– | Lymphocyte # | 7.2e-7 | 1.5e-5 | 22.9 (9.4–55.6) |
| 16q– | Lymphocyte % | 1.6e-5 | 0.00028 | 18.4 (7.1–47.7) |
| 16q– | Monocyte # | 0.004 | 0.04 | 10.2 (3.1–33.7) |
| 17p– | Lymphocyte # | 4.8e-18 | 3.2e-16 | 18.2 (11.3–29.4) |
| 17p– | Lymphocyte % | 8.8e-9 | 2.3e-7 | 10.2 (5.6–18.5) |
| 18p– | Lymphocyte % | 0.001 | 0.013 | 17.5 (5.1–59.7) |
| 18+ | Lymphocyte # | 2.2e-17 | 1.4e-15 | 16.7 (10.4–26.9) |
| 18+ | Lymphocyte % | 2e-8 | 5.1e-7 | 9.4 (5.2–17.0) |
| 19+ | Lymphocyte # | 3.6e-13 | 1.6e-11 | 47.3 (22.2–100.4) |
| 19+ | Lymphocyte % | 2.1e-8 | 5.2e-7 | 28.9 (12.5–67.2) |
| 19+ | Platelet dist. width | 0.0036 | 0.037 | 10.6 (3.2–34.9) |
| 20q– | Neutrophil % | 3.3e-7 | 7.2e-6 | 4.8 (2.9–7.7) |
| 20q– | RBC dist. width | 7.3e-6 | 0.00013 | 4.2 (2.5–7.0) |
| 20q– | Platelet # | 0.0015 | 0.019 | 3.0 (1.7–5.5) |
| 20q– | Platelet crit | 0.00045 | 0.0061 | 3.3 (1.9–5.9) |
| 20q– | Platelet dist. width | 3.3e-7 | 7.2e-6 | 4.8 (2.9–7.7) |
| 20q= | Lymphocyte % | 0.0032 | 0.033 | 4.4 (1.9–9.9) |
| 21q– | Lymphocyte % | 0.0036 | 0.037 | 10.6 (3.2–34.9) |
| 21q– | Platelet dist. width | 0.00025 | 0.0036 | 14.7 (5.1–42.0) |
| 22q– | Lymphocyte # | 1.4e-63 | 1.9e-61 | 90.0 (60.1–134.8) |
| 22q– | Lymphocyte % | 1.6e-48 | 1.4e-46 | 63.7 (42.1–96.3) |
| 22+ | Lymphocyte # | 6.1e-6 | 0.00011 | 6.6 (3.5–12.5) |
| 22+ | Lymphocyte % | 1.2e-8 | 3e-7 | 8.7 (4.9–15.4) |

This table provides numerical data plotted in Supplementary Fig. 27. Mosaic chromosomal alterations significantly enriched (at an FDR threshold of 0.05; one-sided Fisher's exact test) in individuals with anomalous blood indices (top 1% among self-reported white individuals) are reported. Events were grouped by chromosome and copy number, with loss and CNN-LOH events subdivided by p-arm vs. q-arm. (We did not subdivide gain events by arm because most gain events are whole-chromosome trisomies; e.g., "3+" combines all gains—partial or complete—on chromosome 3.)

#### Supplementary Table 6. Rare coding or splice variants associated at FDR<0.05 significance with mosaic CNN-LOH mutations in *cis*.

| | | | | | | | | GWAS | | Allelic shift in hets | | |
|---|---|---|---|---|---|---|---|---|---|---|---|---|
| Chr | Position[a] | Variant | Effect[b] | Alleles[c] | AF[d] | Source | INFO/R2 | $P$ | OR (95% CI) | $N_{REF}$[e] | $N_{ALT}$ | $P$ |
| ***MPL*: 28/61 tested variants significant at FDR<0.05** | | | | | | | | | | | | |
| 1 | 43803600 | rs146249964 | splice donor | T/A | 0.0001 | HRC imp | 0.685 | $2.8\times10^{-23}$ | 97 (55–171) | 12 | 0 | 0.00049 |
| 1 | 43803817 | rs148434485 | stop gained | C/T | $2\times10^{-5}$ | BB array | – | $1.6\times10^{-6}$ | 128 (37–446) | 2 | 0 | 0.5 |
| 1 | 43803824 | rs145714475 | missense | T/C | $2\times10^{-5}$ | HRC imp | 0.394 | $1.9\times10^{-6}$ | 120 (35–414) | 3 | 0 | 0.25 |
| 1 | 43803835 | rs764333753 | missense | A/G | $3\times10^{-5}$ | WES imp | 0.782 | $1.7\times10^{-2}$ | 31 (4–235) | 1 | 0 | 1 |
| 1 | 43803877 | rs766172846 | missense | T/C | $4\times10^{-5}$ | WES imp | 0.849 | $7.7\times10^{-4}$ | 37 (9–156) | 2 | 0 | 0.5 |
| 1 | 43803903 | rs142565191 | splice donor | G/A | $4\times10^{-5}$ | WES imp | 0.914 | $7.5\times10^{-6}$ | 72 (22–238) | 3 | 0 | 0.25 |
| 1 | 43804234 | rs587778514 | frameshift | CCT/C | $1\times10^{-5}$ | BB array | – | $3.9\times10^{-5}$ | 199 (40–987) | 2 | 0 | 0.5 |
| 1 | 43804305 | rs28928907 | missense | G/C | 0.0006 | BB array | – | $1.9\times10^{-130}$ | 142 (111–184) | 70 | 0 | $1.7\times10^{-21}$ |
| 1 | 43804375 | rs587778515 | frameshift | CT/C | 0.0002 | BB array | – | $7.0\times10^{-41}$ | 105 (68–161) | 24 | 0 | $1.2\times10^{-7}$ |
| 1 | 43804396 | rs752453717 | splice modifier | G/C | 0.0003 | BB array | – | $5.8\times10^{-36}$ | 74 (48–113) | 24 | 0 | $1.2\times10^{-7}$ |
| 1 | 43804957 | rs764904424 | missense | C/G | 0.0001 | WES imp | 0.750 | $2.1\times10^{-8}$ | 35 (15–79) | 6 | 0 | 0.031 |
| 1 | 43805052 | rs6088 | missense | G/A | $9\times10^{-5}$ | WES imp | 0.825 | $8.3\times10^{-10}$ | 61 (26–141) | 6 | 0 | 0.031 |
| 1 | 43805059 | rs769867913 | missense | G/A | $2\times10^{-5}$ | WES imp | 0.439 | $1.4\times10^{-2}$ | 37 (5–282) | 1 | 0 | 1 |
| 1 | 43805656 | rs144210383 | missense | G/T | 0.0001 | WES imp | 0.676 | $5.3\times10^{-9}$ | 44 (19–101) | 6 | 0 | 0.031 |
| 1 | 43805686 | rs587778518 | frameshift | C/CCTGG | $3\times10^{-5}$ | WES imp | 0.825 | $1.6\times10^{-2}$ | 33 (4–249) | 1 | 0 | 1 |
| 1 | 43805713 | rs121913611 | missense | C/T | 0.0002 | BB array | – | $3.3\times10^{-28}$ | 102 (61–171) | 17 | 0 | $1.5\times10^{-5}$ |
| 1 | 43806073 | 1:43806073 | missense | A/C | $2\times10^{-5}$ | WES imp | 0.853 | $1.5\times10^{-4}$ | 92 (21–408) | 2 | 0 | 0.5 |
| 1 | 43812115 | rs769297582 | splice acceptor | G/C | $2\times10^{-5}$ | WES imp | 0.760 | $5.1\times10^{-7}$ | 199 (54–737) | 3 | 0 | 0.25 |
| 1 | 43812574 | rs200454070 | missense | G/A | $3\times10^{-5}$ | WES imp | 0.568 | $2.0\times10^{-2}$ | 26 (4–192) | 1 | 0 | 1 |
| 1 | 43814551 | rs765671565 | missense | T/A | $9\times10^{-6}$ | WES imp | 0.771 | $5.9\times10^{-3}$ | 100 (12–827) | 1 | 0 | 1 |
| 1 | 43814590 | rs1175548872 | missense | G/C | $1\times10^{-5}$ | WES imp | 0.898 | $7.5\times10^{-3}$ | 75 (9–597) | 1 | 0 | 1 |
| 1 | 43814627 | rs754859909 | stop gained | G/A | $7\times10^{-5}$ | WES imp | 0.939 | $1.7\times10^{-16}$ | 126 (61–258) | 9 | 0 | 0.0039 |
| 1 | 43814673 | rs923814653 | missense | G/T | $3\times10^{-5}$ | WES imp | 0.966 | $4.1\times10^{-4}$ | 52 (12–221) | 3 | 0 | 0.25 |
| 1 | 43814729 | 454bp del[f] | exon 10 deletion | ref/del | 0.0002 | array LRR | – | $3.6\times10^{-58}$ | 153 (104–225) | 31 | 0 | $9.3\times10^{-10}$ |
| 1 | 43815009 | rs121913615 | missense | G/T | $2\times10^{-5}$ | WES imp | 0.936 | $1.4\times10^{-2}$ | 37 (5–282) | 1 | 0 | 1 |
| 1 | 43817942 | rs369156948 | stop gained | C/T | $3\times10^{-5}$ | HRC imp | 0.225 | $4.8\times10^{-8}$ | 114 (39–333) | 4 | 0 | 0.12 |
| 1 | 43817973 | rs971379181 | frameshift | CG/C | $3\times10^{-5}$ | BB array | – | $5.8\times10^{-13}$ | 240 (93–618) | 6 | 0 | 0.031 |
| 1 | 43818435 | rs1366403560 | stop gained | C/T | $2\times10^{-5}$ | WES imp | 0.866 | $1.3\times10^{-4}$ | 100 (22–446) | 2 | 0 | 0.5 |
| ***FH*: 1/41 tested variants significant at FDR<0.05** | | | | | | | | | | | | |
| 1 | 241675301 | rs199822819 | missense | G/C | 0.0003 | WES imp | 0.869 | $4.9\times10^{-11}$ | 28 (14–55) | 1 | 8 | 0.039 |
| ***NBN*: 2/48 tested variants significant at FDR<0.05** | | | | | | | | | | | | |
| 8 | 90983420 | rs777460725 | missense | A/C | 0.0001 | WES imp | 0.794 | $8.1\times10^{-5}$ | 114 (28–465) | 0 | 2 | 0.5 |
| 8 | 90983441 | rs1187082186 | frameshift | ATTTGT/A | 0.0002 | WES imp | 0.844 | $4.8\times10^{-13}$ | 210 (92–484) | 0 | 6 | 0.031 |
| ***MRE11*: 1/42 tested variants significant at FDR<0.05** | | | | | | | | | | | | |
| 11 | 94189489 | rs587781384 | stop gained | C/A | $4\times10^{-5}$ | WES imp | 0.945 | $5.6\times10^{-10}$ | 130 (50–338) | 0 | 5 | 0.062 |
| ***ATM*: 13/352 tested variants significant at FDR<0.05** | | | | | | | | | | | | |
| 11 | 108127067 | rs1137887 | splice modifier | G/A | $4\times10^{-5}$ | WES imp | 0.768 | $9.6\times10^{-6}$ | 65 (20–214) | 0 | 2 | 0.5 |
| 11 | 108141801 | rs786203054 | missense | T/G | $7\times10^{-6}$ | BB array | – | $1.2\times10^{-5}$ | 437 (73–2618) | 0 | 2 | 0.5 |
| 11 | 108155007 | rs781357995 | frameshift | AG/A | 0.0001 | WES imp | 0.888 | $3.0\times10^{-9}$ | 48 (21–111) | 0 | 6 | 0.031 |
| 11 | 108172425 | rs587779844 | missense | C/T | 0.0001 | BB array | – | $3.5\times10^{-20}$ | 96 (52–177) | 0 | 12 | 0.00049 |
| 11 | 108175420 | rs786204751 | stop gained | C/T | $2\times10^{-5}$ | BLVE imp | 0.467 | $1.6\times10^{-4}$ | 87 (20–380) | 0 | 1 | 1 |
| 11 | 108175528 | rs376603775 | stop gained | C/T | $6\times10^{-5}$ | BLVE imp | 0.848 | $2.8\times10^{-5}$ | 44 (14–143) | 0 | 4 | 0.12 |
| 11 | 108179837 | rs774925473 | splice modifier | A/G | $8\times10^{-5}$ | BLVE imp | 0.677 | $6.8\times10^{-5}$ | 33 (10–104) | 0 | 3 | 0.25 |
| 11 | 108181006 | rs56399311 | missense | A/G | $8\times10^{-5}$ | WES imp | 0.957 | $1.7\times10^{-6}$ | 44 (16–120) | 0 | 4 | 0.12 |
| 11 | 108201108 | rs56399857 | missense | T/G | 0.0002 | WES imp | 0.921 | $4.9\times10^{-5}$ | 18 (6.6–48) | 0 | 4 | 0.12 |
| 11 | 108202611 | rs587776547 | inframe deletion | CTCTAGAATT/C | $7\times10^{-5}$ | WES imp | 0.754 | $8.5\times10^{-9}$ | 73 (29–183) | 0 | 5 | 0.062 |
| 11 | 108206686 | rs371638537 | stop gained | A/T | $6\times10^{-5}$ | WES imp | 0.877 | $9.9\times10^{-4}$ | 33 (8–135) | 0 | 2 | 0.5 |
| 11 | 108216545 | rs587779872 | missense | C/T | $2\times10^{-5}$ | WES imp | 0.594 | $3.6\times10^{-11}$ | 251 (89–706) | 0 | 5 | 0.062 |
| 11 | 108224608 | rs17174393 | splice donor | G/A | $4\times10^{-5}$ | WES imp | 0.824 | $6.5\times10^{-4}$ | 41 (10–170) | 0 | 2 | 0.5 |
| ***SH2B3*: 2/57 tested variants significant at FDR<0.05** | | | | | | | | | | | | |
| 12 | 111885295 | rs148636776 | missense | G/A | 0.0004 | WES imp | 0.861 | $4.0\times10^{-5}$ | 19 (7–50) | 0 | 5 | 0.062 |
| 12 | 111885310 | rs72650673 | missense | G/A | 0.002 | WES imp | 0.882 | $3.1\times10^{-8}$ | 11 (5.8–20) | 1 | 8 | 0.039 |
| ***TM2D3*: 5/15 tested variants significant at FDR<0.05** | | | | | | | | | | | | |
| 15 | 102151467 | 70kb del[g] | gene deletion | ref/del | 0.0003 | array LRR | – | $9.8\times10^{-224}$ | 555 (425–724) | 2 | 110 | $2.4\times10^{-30}$ |
| 15 | 102182739 | rs113189685 | missense | G/T | $3\times10^{-5}$ | WES imp | 0.544 | $2.8\times10^{-8}$ | 132 (45–389) | 1 | 3 | 0.62 |
| 15 | 102182749 | rs754640606 | missense | G/C | $5\times10^{-5}$ | WES imp | 0.769 | $1.2\times10^{-40}$ | 544 (289–1025) | 0 | 19 | $3.8\times10^{-6}$ |
| 15 | 102182761 | rs976377433 | missense | A/G | $3\times10^{-5}$ | WES imp | 0.761 | $2.3\times10^{-8}$ | 140 (47–413) | 0 | 4 | 0.12 |
| 15 | 102190214 | rs768556490 | frameshift | G/GT | $3\times10^{-5}$ | WES imp | 0.850 | $8.2\times10^{-29}$ | 758 (327–1759) | 1 | 11 | 0.0063 |

See next page for caption.

**Caption for Supplementary Table 6.**

Results of two independent statistical tests are reported: (i) a Fisher test treating individuals with a mosaic CNN-LOH mutation in *cis* as cases; and (ii) a binomial test for biased allelic imbalance in heterozygous cases. Loci reaching FDR$<$0.05 significance in the first test (based on the number of variants tested in each gene) are reported. Note that both tests are two-sided (to retain consistency with Table 1); *P*-values for biased allelic imbalance in the expected directions (removing rare alleles in *MPL* and duplicating rare alleles in other genes) would be half the values reported in the last column of this table. For full details of statistical tests, see Methods.

[a]Base pair position in hg19 coordinates.
[b]Variant effects according to Ensembl VEP [20] (coding variants) or ClinVar [21] (splice variants).
[c]Reference/alternate allele.
[d]Alternate allele frequency (in UK Biobank European-ancestry individuals).
[e]Number of mosaic individuals heterozygous for the variant in which the somatic event shifted the allelic balance in favor of the reference allele (by duplication of its chromosomal segment and loss of the homologous segment).
[f]This 454bp deletion spans chr1:43,814,729-43,815,182, deleting *MPL* exon 10 (Supplementary Figures 28–30).
[g]This $\sim$70kb deletion spans chr15:102.15–102.22Mb, deleting *TM2D3* and part of *TARSL2* [10].

**Supplementary Table 7. Previously reported CNN-LOH risk variants at *MPL* and *ATM* tag likely causal coding variants.**

| Locus | Previously reported variant [10] | Likely causal coding variant in Table 1 | $R^2$ |
|---|---|---|---|
| *MPL* | rs182971382 | rs28928907 (missense) | 0.83 |
| *MPL* | rs144279563 | 454bp del (exon 10 deletion) | 0.36 |
| *MPL* | rs369156948 | rs369156948 (stop gained) | 1 |
| *ATM* | rs532198118 | rs587779844 (missense) | 0.19 |

Note that $R^2$ reported above may be underestimated due to imputation error.

**Supplementary Table 8. Numbers of distinct coding or splice variants at each risk locus likely to causally drive associations with mosaic CNN-LOH events in *cis*.**

| Gene | Mosaic event | Variants associated at Bonferroni significance | Variants associated at FDR<0.05 significance | Additional coding or splice variants contributing to ultra-rare burden |
|---|---|---|---|---|
| *MPL* | 1p CNN-LOH | 17 | 28 | +4 |
| *FH* | 1q CNN-LOH | 1 | 1 | +2 |
| *NBN* | 8q CNN-LOH | 2 | 2 | +0 |
| *MRE11* | 11q CNN-LOH | 1 | 1 | +1 |
| *ATM* | 11q CNN-LOH | 10 | 13 | +6 |
| *SH2B3* | 12q CNN-LOH | 2 | 2 | +5 |
| *TM2D3* | 15q CNN-LOH | 5 | 5 | +3 |
| Total | | 38 | 52 | +21 |

This table summarizes the numbers of likely-causal variants at each CNN-LOH risk locus identified by our association analyses (Table 1 and Supplementary Table 6) and burden analyses (Supplementary Table 9).

**Supplementary Table 9. Burden of additional ultra-rare coding or splice variants in genes frequently targeted by CNN-LOH events in *cis*.**

| Gene | Mosaic event | Exome-sequenced mosaic individuals not carrying FDR-significant variant (Supplementary Table 6) | Carriers of ultra-rare coding or splice variants among these individuals: Observed | Expected | $P$ |
|---|---|---|---|---|---|
| *MPL* | 1p CNN-LOH | 37 | 5 | 0.067 | $8.4\times10^{-9}$ |
| *FH* | 1q CNN-LOH | 51 | 2 | 0.132 | 0.0079 |
| *NBN* | 8q CNN-LOH | 5 | 0 | 0.017 | 1 |
| *MRE11* | 11q CNN-LOH | 52 | 1 | 0.222 | 0.20 |
| *ATM* | 11q CNN-LOH | 57 | 6 | 0.682 | $6.3\times10^{-5}$ |
| *SH2B3* | 12q CNN-LOH | 25 | 5 | 0.110 | $8.3\times10^{-8}$ |
| *TM2D3* | 15q CNN-LOH | 44 | 3 | 0.051 | $2.0\times10^{-5}$ |
| *DNMT3A* | 2p CNN-LOH | 16 | 4 | 0.206 | $4.4\times10^{-5}$ |
| *TET2* | 4q CNN-LOH | 21 | 6 | 0.197 | $3.3\times10^{-8}$ |
| *JAK2* | 9p CNN-LOH | 33 | 15 | 0.217 | $1.7\times10^{-24}$ |

For each gene, we examined individuals with CNN-LOH events spanning the gene (not already explained by any of the 52 variants identified in our association analyses) and tabulated the number of such individuals who carried a rare coding or splice variant under consideration (see Methods). We then computed a burden $P$-value using a one-sided binomial test comparing the observed count to expectation. Note that the counts of observed carriers include two 1p CNN-LOH individuals with the same *MPL* variant (rs1362911656, 1:43814994_T_C) and fifteen 9p CNN-LOH individuals with *JAK2* V617F. An additional five 9p CNN-LOH individuals had at least one read supporting *JAK2* V617F (but did not have *JAK2* V617F genotype calls). Allelic read depth analyses indicated that all or most of the rare variant burden in the seven inherited risk loci arose from inherited variants, while all or most of the burden in *DNMT3A*, *TET2*, and *JAK2* arose from somatic point mutations (Supplementary Fig. 33).

#### Supplementary Table 10. Rare coding or splice variants carried by exome-sequenced individuals with mosaic CNN-LOH events spanning frequently-targeted genes.

| Gene | Mosaic event | Fraction of mosaic individuals with a rare coding/splice variant | Count | Variant | Effect |
|---|---|---|---|---|---|
| *MPL* | 1p CNN-LOH | 39 / 71 | 8 | 1:43804305_G_C | missense |
| | | (expected: 0.525 / 71) | 5 | 1:43804396_G_C | splice modifier |
| | | | 4 | 454bp del | exon 10 deletion |
| | | | 3 | 1:43805713_C_T | missense |
| | | | 2 | 1:43804375_CT_C | frameshift |
| | | | 2 | 1:43814994_T_C | missense |
| | | | 1 | 1:43803600_T_A | splice donor |
| | | | 1 | 1:43803903_G_A | splice donor |
| | | | 1 | 1:43804268_C_T | stop gained |
| | | | 1 | 1:43804957_C_G | missense |
| | | | 1 | 1:43805052_G_A | missense |
| | | | 1 | 1:43805059_G_A | missense |
| | | | 1 | 1:43805656_G_T | missense |
| | | | 1 | 1:43805686_C_CCTGG | frameshift |
| | | | 1 | 1:43812115_G_C | splice acceptor |
| | | | 1 | 1:43812574_G_A | missense |
| | | | 1 | 1:43814563_C_G | missense |
| | | | 1 | 1:43814627_G_A | stop gained |
| | | | 1 | 1:43814673_G_T | missense |
| | | | 1 | 1:43815009_G_T | missense |
| | | | 1 | 1:43818405_C_G | missense |
| *FH* | 1q CNN-LOH | 3 / 52 | 1 | 1:241675301_G_C | missense |
| | | (expected: 0.163 / 52) | 1 | 1:241675313_C_T | missense |
| | | | 1 | 1:241675443_C_A | missense |
| *NBN* | 8q CNN-LOH | 2 / 7 | 1 | 8:90983420_A_C | missense |
| | | (expected: 0.027 / 7) | 1 | 8:90983441_ATTTGT_A | frameshift |
| *MRE11* | 11q CNN-LOH | 2 / 57 | 1 | 11:94189447_G_A | missense |
| | | (expected: 0.247 / 57) | 1 | 11:94189489_C_A | stop gained |
| *ATM* | 11q CNN-LOH | 12 / 64 | 2 | 11:108155007_AG_A | frameshift |
| | | (expected: 0.880 / 64) | 1 | 11:108115595_G_T | missense |
| | | | 1 | 11:108121479_CTG_C | frameshift |
| | | | 1 | 11:108121546_AC_A | frameshift |
| | | | 1 | 11:108127067_G_A | splice modifier |
| | | | 1 | 11:108159805_T_C | missense |
| | | | 1 | 11:108172383_T_C | missense |
| | | | 1 | 11:108179837_A_G | splice modifier |
| | | | 1 | 11:108199833_G_A | missense |
| | | | 1 | 11:108202611_CTCTAGAATT_C | inframe deletion |
| | | | 1 | 11:108216545_C_T | missense |
| *SH2B3* | 12q CNN-LOH | 6 / 26 | 1 | 12:111856537_G_GT | frameshift |
| | | (expected: 0.226 / 26) | 1 | 12:111856620_T_TGC | frameshift |
| | | | 1 | 12:111856623_G_GCCGGGCC | frameshift |
| | | | 1 | 12:111884838_G_A | splice donor |
| | | | 1 | 12:111885295_G_A | missense |
| | | | 1 | 12:111885497_G_A | missense |
| *TM2D3* | 15q CNN-LOH | 20 / 61 | 8 | 70kb del | gene deletion |
| | | (expected: 0.131 / 61) | 4 | 15:102190214_G_GT | frameshift |
| | | | 3 | 15:102182749_G_C | missense |
| | | | 2 | 15:102182739_G_T | missense |
| | | | 1 | 15:102187018_A_G | missense |
| | | | 1 | 15:102192520_A_T | stop gained |
| | | | 1 | 15:102192558_C_A | stop gained |
| *DNMT3A* | 2p CNN-LOH | 4 / 16 | 1 | 2:25463194_T_C | missense |
| | | (expected: 0.206 / 16) | 1 | 2:25463218_C_T | missense |
| | | | 1 | 2:25463248_G_A | missense |
| | | | 1 | 2:25463536_C_T | missense |
| *TET2* | 4q CNN-LOH | 6 / 21 | 1 | 4:106157029_C_T | stop gained |
| | | (expected: 0.197 / 21) | 1 | 4:106157446_G_T, 4:106157983_C_T, 4:106193937_AG_A | stop gained, frameshift |
| | | | 1 | 4:106164061_C_T | stop gained |
| | | | 1 | 4:106164085_G_GT | splice donor |
| | | | 1 | 4:106180896_G_GT | frameshift |
| | | | 1 | 4:106197285_T_C | missense |
| *JAK2* | 9p CNN-LOH | 15 / 33 | 15 | 9:5073770_G_T | missense |
| | | (expected: 0.217 / 33) | | | |

See next page for caption.

**Caption for Supplementary Table 10.**

This table lists variants found in exome-sequenced mosaic CNN-LOH individuals at each locus at which inherited or somatic variants are targets of clonal CNN-LOH events. Variants identified by our association analyses (Table 1 and Supplementary Table 6) and burden analyses (Supplementary Table 9) are both included. Allelic read depth analyses indicated that all or most of the variants found in the seven inherited risk loci arose from inherited variants, while all or most of the variants found in *DNMT3A*, *TET2*, and *JAK2* arose from somatic point mutations (Supplementary Fig. 33). We note that while this table indicates that fifteen individuals with 9p CNN-LOH events were carriers of *JAK2* V617F (9:5073770_G_T), an additional five 9p CNN-LOH individuals had at least one read supporting *JAK2* V617F (but did not have *JAK2* V617F genotype calls). One individual with 4q CNN-LOH appeared to have three distinct somatic mutations in *TET2*.

**Supplementary Table 11. Associations of mosaic CNN-LOH mutations with inherited common variants in *cis*.**

| Arm | Locus | Position[a] | Variant | Alleles[b] | AF[c] | GWAS | | Allelic shift in hets | | | |
|---|---|---|---|---|---|---|---|---|---|---|---|
| | | | | | | $P$ | OR (95% CI) | $N_{REF}$[d] | $N_{ALT}$ | $P$ | $P_{combined}$ |
| **Novel common variant associations with CNN-LOH in *cis*** | | | | | | | | | | | |
| 14q | *TCL1A* | 96180695 | rs2887399 | G/T | 0.20 | 0.0024 | 0.84 (0.75–0.94) | 195 | 102 | $7.4\times10^{-8}$ | $4.2\times10^{-9}$ |
| 14q | *DLK1* | 101172227 | rs7141110 | G/C | 0.22 | $1.4\times10^{-5}$ | 1.24 (1.13–1.37) | 252 | 162 | $1.1\times10^{-5}$ | $3.6\times10^{-9}$ |
| **Previously reported common variant associations with CNN-LOH in *cis*** | | | | | | | | | | | |
| 9p | *JAK2* | 5037393 | rs75032480[e] | A/C | 0.26 | $2.6\times10^{-29}$ | 2.29 (1.99–2.63) | 31 | 170 | $2\times10^{-24}$ | $6.3\times10^{-51}$ |

Results of two independent statistical tests are reported: (i) a Fisher test treating individuals with a mosaic CNN-LOH mutation in *cis* as cases; and (ii) a binomial test for biased allelic imbalance in heterozygous cases. Loci reaching genome-wide significance in the combination of the tests (Fisher's combined $P$) are reported. For full details of statistical tests, see Methods.

[a]Base pair position in hg19 coordinates.

[b]Reference/alternate allele.

[c]Alternate allele frequency (in UK Biobank European-ancestry individuals).

[d]Number of mosaic individuals heterozygous for the variant in which the somatic event shifted the allelic balance in favor of the reference allele (by duplication of its chromosomal segment and loss of the homologous segment).

[e]rs75032480 belongs to the *JAK2* 46/1 haplotype [47–49].

**Supplementary Table 12. Common variants associated with detectable mosaic chromosomal alterations on any autosome.**

| Locus | Variant | Chr | Position | REF/ALT | AAF | OR (95% CI) | *P* |
|---|---|---|---|---|---|---|---|
| *SP140* | rs13023767 | 2 | 231122057 | T/G | 0.25 | 1.07 (1.05–1.10) | $2.3\times10^{-8}$ |
| | rs776205558 | 2 | 231122089 | CAGTA/C | 0.25 | 1.07 (1.05–1.10) | $3.2\times10^{-8}$ |
| | rs55657711 | 2 | 231122210 | A/G | 0.30 | 1.07 (1.05–1.10) | $2.0\times10^{-8}$ |
| | rs62191185 | 2 | 231122290 | G/A | 0.42 | 1.06 (1.04–1.09) | $4.0\times10^{-8}$ |
| | rs1356532206 | 2 | 231124230 | TA/T | 0.26 | 1.07 (1.05–1.10) | $3.3\times10^{-8}$ |
| | rs6755306 | 2 | 231126528 | G/A | 0.25 | 1.08 (1.05–1.10) | $1.2\times10^{-8}$ |
| | rs1582833 | 2 | 231129729 | C/G | 0.31 | 1.07 (1.05–1.10) | $1.7\times10^{-8}$ |
| | rs62191195 | 2 | 231129794 | C/T | 0.25 | 1.08 (1.05–1.10) | $9.4\times10^{-9}$ |
| | rs34790921 | 2 | 231130508 | G/T | 0.25 | 1.08 (1.05–1.10) | $1.2\times10^{-8}$ |
| | rs890581 | 2 | 231131387 | G/A | 0.25 | 1.08 (1.05–1.10) | $9.7\times10^{-9}$ |
| | rs767031837 | 2 | 231134078 | AGCGTG/A | 0.25 | 1.08 (1.05–1.10) | $1.2\times10^{-8}$ |
| | rs62191198 | 2 | 231141196 | C/G | 0.25 | 1.08 (1.05–1.10) | $9.5\times10^{-9}$ |
| | rs12694846 | 2 | 231148128 | A/G | 0.27 | 1.07 (1.05–1.10) | $2.1\times10^{-8}$ |
| | rs34004493 | 2 | 231154012 | A/G | 0.27 | 1.07 (1.05–1.10) | $2.6\times10^{-8}$ |
| | rs6710297 | 2 | 231157512 | A/G | 0.27 | 1.07 (1.05–1.10) | $2.2\times10^{-8}$ |
| | rs35256947 | 2 | 231161026 | T/C | 0.27 | 1.07 (1.05–1.10) | $2.0\times10^{-8}$ |
| | rs13007094 | 2 | 231171194 | C/T | 0.25 | 1.08 (1.05–1.10) | $1.1\times10^{-8}$ |
| | rs2396742 | 2 | 231171423 | C/T | 0.25 | 1.08 (1.05–1.10) | $1.9\times10^{-8}$ |
| *TERC* | rs12638862 | 3 | 169477506 | A/G | 0.26 | 0.93 (0.91–0.96) | $2.9\times10^{-8}$ |
| | rs9811216 | 3 | 169487501 | T/C | 0.26 | 0.93 (0.91–0.96) | $3.4\times10^{-8}$ |
| *TERT* | rs33961405 | 5 | 1277577 | G/A | 0.52 | 0.93 (0.91–0.96) | $6.4\times10^{-9}$ |
| | rs10054203 | 5 | 1279964 | G/C | 0.40 | 1.07 (1.05–1.10) | $3.0\times10^{-9}$ |
| | rs7734992 | 5 | 1280128 | T/C | 0.42 | 1.09 (1.06–1.11) | $4.2\times10^{-13}$ |
| | rs4975538 | 5 | 1280830 | G/C | 0.36 | 1.08 (1.05–1.10) | $1.2\times10^{-10}$ |
| | rs6897196 | 5 | 1280938 | A/G | 0.39 | 1.08 (1.06–1.10) | $5.6\times10^{-11}$ |
| | rs749685059 | 5 | 1280940 | GAGCCCACC/G | 0.38 | 1.08 (1.06–1.11) | $8.0\times10^{-12}$ |
| | rs7726159 | 5 | 1282319 | C/A | 0.33 | 1.10 (1.07–1.12) | $2.8\times10^{-14}$ |
| | rs7725218 | 5 | 1282414 | G/A | 0.34 | 1.09 (1.06–1.11) | $2.0\times10^{-12}$ |
| | rs4449583 | 5 | 1284135 | C/T | 0.33 | 1.10 (1.07–1.12) | $2.3\times10^{-14}$ |
| | rs7705526 | 5 | 1285974 | C/A | 0.33 | 1.11 (1.08–1.14) | $6.9\times10^{-18}$ |
| | rs2736100 | 5 | 1286516 | C/A | 0.50 | 0.92 (0.90–0.94) | $1.2\times10^{-12}$ |
| | rs2853677 | 5 | 1287194 | G/A | 0.58 | 0.94 (0.92–0.96) | $9.5\times10^{-9}$ |

Results from BOLT-LMM [26, 44] analysis of the "any autosomal mCA" phenotype are reported for all common variants (MAF>0.05) passing a significance threshold of $P<5\times10^{-8}$. AAF = ALT allele frequency; the ALT allele is the effect allele for reported odds ratios.

**Supplementary Table 13. Mean changes in polygenic scores for blood count and Y loss traits produced by CNN-LOH mutations.**

| Arm | Platelet # | Red cell # | Basophil # | Neutrophil # | Eosinophil # | Monocyte # | Lymphocyte # | Y loss risk |
|---|---|---|---|---|---|---|---|---|
| 1p | **–0.0632 (0.0055)** | 0.0032 (0.0022) | –0.0003 (0.0006) | 0.0005 (0.0022) | 0.0013 (0.0021) | 0.0022 (0.0022) | –0.0011 (0.0020) | 0.0003 (0.0004) |
| 1q | 0.0056 (0.0040) | –0.0029 (0.0030) | 0.0003 (0.0013) | 0.0034 (0.0027) | –0.0045 (0.0023) | –0.0037 (0.0031) | 0.0002 (0.0022) | 0.0011 (0.0006) |
| 2p | 0.0026 (0.0053) | 0.0035 (0.0056) | 0.0020 (0.0011) | 0.0061 (0.0039) | 0.0071 (0.0038) | –0.0023 (0.0035) | 0.0035 (0.0042) | –0.0006 (0.0010) |
| 2q | **0.0192 (0.0059)** | 0.0055 (0.0046) | 0.0014 (0.0015) | **0.0153 (0.0063)** | 0.0048 (0.0048) | 0.0094 (0.0068) | 0.0043 (0.0043) | **–0.0014 (0.0007)** |
| 3p | 0.0007 (0.0057) | 0.0053 (0.0042) | 0.0013 (0.0017) | **0.0082 (0.0030)** | –0.0066 (0.0043) | 0.0045 (0.0059) | 0.0016 (0.0042) | 0.0011 (0.0009) |
| 3q | 0.0084 (0.0062) | –0.0080 (0.0047) | 0.0016 (0.0014) | **0.0087 (0.0036)** | 0.0051 (0.0045) | **0.0093 (0.0047)** | 0.0015 (0.0035) | 0.0015 (0.0013) |
| 4p | 0.0018 (0.0060) | 0.0037 (0.0034) | 0.0000 (0.0009) | 0.0054 (0.0043) | 0.0001 (0.0043) | **0.0107 (0.0040)** | 0.0028 (0.0043) | 0.0010 (0.0009) |
| 4q | 0.0004 (0.0038) | –0.0012 (0.0040) | 0.0000 (0.0010) | 0.0008 (0.0037) | 0.0014 (0.0030) | 0.0006 (0.0034) | –0.0011 (0.0035) | 0.0005 (0.0006) |
| 5p | 0.0004 (0.0049) | 0.0003 (0.0044) | –0.0002 (0.0009) | 0.0061 (0.0041) | –0.0069 (0.0038) | **0.0099 (0.0025)** | –0.0022 (0.0035) | –0.0001 (0.0009) |
| 5q | 0.0094 (0.0052) | –0.0001 (0.0037) | 0.0018 (0.0011) | 0.0000 (0.0042) | 0.0011 (0.0064) | **0.0104 (0.0047)** | 0.0006 (0.0037) | 0.0004 (0.0009) |
| 6p | **–0.0090 (0.0043)** | –0.0050 (0.0036) | 0.0003 (0.0008) | –0.0006 (0.0027) | 0.0045 (0.0036) | 0.0035 (0.0035) | 0.0004 (0.0034) | 0.0008 (0.0005) |
| 6q | –0.0037 (0.0086) | –0.0011 (0.0106) | 0.0005 (0.0013) | 0.0004 (0.0035) | 0.0036 (0.0050) | 0.0028 (0.0053) | 0.0031 (0.0048) | 0.0021 (0.0018) |
| 7p | 0.0039 (0.0044) | 0.0024 (0.0047) | –0.0012 (0.0018) | –0.0047 (0.0050) | 0.0009 (0.0052) | 0.0046 (0.0052) | 0.0002 (0.0047) | 0.0007 (0.0015) |
| 7q | 0.0007 (0.0062) | –0.0030 (0.0050) | –0.0018 (0.0017) | 0.0024 (0.0031) | 0.0039 (0.0043) | 0.0037 (0.0035) | 0.0006 (0.0034) | **0.0030 (0.0013)** |
| 8p | 0.0058 (0.0035) | 0.0013 (0.0038) | –0.0032 (0.0031) | –0.0037 (0.0042) | –0.0030 (0.0037) | 0.0018 (0.0052) | –0.0029 (0.0042) | 0.0010 (0.0008) |
| 8q | –0.0025 (0.0057) | –0.0002 (0.0038) | 0.0002 (0.0012) | 0.0001 (0.0041) | 0.0010 (0.0045) | –0.0011 (0.0068) | –0.0005 (0.0057) | 0.0004 (0.0008) |
| 9p | –0.0081 (0.0054) | 0.0043 (0.0023) | 0.0004 (0.0004) | **0.0053 (0.0019)** | 0.0067 (0.0034) | –0.0028 (0.0019) | –0.0035 (0.0019) | 0.0001 (0.0003) |
| 9q | 0.0050 (0.0039) | –0.0029 (0.0033) | –0.0002 (0.0006) | 0.0004 (0.0024) | 0.0024 (0.0026) | –0.0085 (0.0044) | 0.0003 (0.0021) | –0.0002 (0.0006) |
| 10p | 0.0059 (0.0059) | –0.0005 (0.0062) | –0.0003 (0.0037) | 0.0053 (0.0041) | 0.0050 (0.0075) | 0.0011 (0.0054) | –0.0031 (0.0053) | –0.0009 (0.0009) |
| 10q | –0.0029 (0.0077) | –0.0024 (0.0054) | –0.0005 (0.0021) | 0.0086 (0.0055) | 0.0105 (0.0053) | 0.0017 (0.0065) | 0.0020 (0.0050) | –0.0001 (0.0008) |
| 11p | 0.0012 (0.0031) | 0.0014 (0.0022) | 0.0007 (0.0004) | 0.0006 (0.0019) | 0.0032 (0.0017) | –0.0015 (0.0018) | 0.0002 (0.0014) | –0.0001 (0.0004) |
| 11q | 0.0049 (0.0032) | **0.0084 (0.0019)** | –0.0004 (0.0011) | 0.0039 (0.0021) | 0.0024 (0.0023) | 0.0007 (0.0023) | 0.0014 (0.0020) | **0.0032 (0.0005)** |
| 12p | 0.0012 (0.0079) | **–0.0122 (0.0055)** | –0.0000 (0.0016) | 0.0016 (0.0036) | –0.0012 (0.0060) | –0.0072 (0.0052) | 0.0154 (0.0081) | –0.0001 (0.0022) |
| 12q | **0.0358 (0.0090)** | **0.0105 (0.0041)** | 0.0015 (0.0011) | **0.0112 (0.0034)** | **0.0180 (0.0042)** | **0.0115 (0.0035)** | **0.0154 (0.0041)** | 0.0003 (0.0007) |
| 13q | –0.0002 (0.0028) | –0.0024 (0.0020) | 0.0000 (0.0005) | –0.0005 (0.0017) | –0.0006 (0.0019) | –0.0068 (0.0036) | –0.0002 (0.0023) | 0.0004 (0.0005) |
| 14q | **0.0097 (0.0029)** | 0.0023 (0.0017) | –0.0008 (0.0005) | **0.0036 (0.0015)** | **0.0038 (0.0018)** | 0.0024 (0.0019) | 0.0003 (0.0015) | **0.0039 (0.0007)** |
| 15q | 0.0042 (0.0025) | –0.0016 (0.0024) | 0.0003 (0.0009) | –0.0028 (0.0018) | 0.0016 (0.0021) | –0.0001 (0.0025) | 0.0031 (0.0020) | –0.0005 (0.0003) |
| 16p | –0.0019 (0.0025) | 0.0040 (0.0029) | **0.0011 (0.0004)** | 0.0010 (0.0016) | 0.0018 (0.0029) | 0.0008 (0.0020) | 0.0008 (0.0020) | 0.0002 (0.0003) |
| 16q | –0.0015 (0.0034) | **0.0071 (0.0034)** | 0.0002 (0.0007) | 0.0015 (0.0027) | 0.0003 (0.0028) | –0.0034 (0.0068) | 0.0008 (0.0026) | –0.0004 (0.0007) |
| 17p | 0.0037 (0.0055) | –0.0030 (0.0033) | –0.0015 (0.0015) | –0.0039 (0.0049) | 0.0023 (0.0043) | –0.0038 (0.0046) | –0.0036 (0.0045) | –0.0001 (0.0008) |
| 17q | 0.0012 (0.0034) | –0.0015 (0.0028) | 0.0007 (0.0011) | 0.0005 (0.0034) | –0.0019 (0.0026) | –0.0007 (0.0035) | 0.0022 (0.0026) | 0.0001 (0.0005) |
| 18p | –0.0011 (0.0046) | –0.0036 (0.0046) | –0.0006 (0.0010) | –0.0029 (0.0039) | 0.0034 (0.0039) | 0.0020 (0.0037) | –0.0028 (0.0025) | –0.0006 (0.0011) |
| 18q | 0.0039 (0.0042) | 0.0049 (0.0037) | **0.0044 (0.0014)** | **0.0074 (0.0035)** | **0.0136 (0.0040)** | 0.0058 (0.0043) | –0.0003 (0.0031) | **0.0097 (0.0027)** |
| 19p | 0.0001 (0.0037) | 0.0027 (0.0044) | 0.0026 (0.0013) | 0.0019 (0.0034) | –0.0027 (0.0035) | –0.0029 (0.0047) | –0.0033 (0.0050) | 0.0006 (0.0006) |
| 19q | 0.0021 (0.0035) | 0.0014 (0.0030) | 0.0020 (0.0012) | 0.0023 (0.0029) | –0.0011 (0.0029) | –0.0010 (0.0043) | –0.0005 (0.0024) | 0.0002 (0.0005) |
| 20p | –0.0018 (0.0049) | –0.0011 (0.0031) | 0.0002 (0.0013) | –0.0027 (0.0032) | –0.0035 (0.0033) | –0.0035 (0.0048) | –0.0039 (0.0049) | 0.0003 (0.0007) |
| 20q | 0.0070 (0.0039) | 0.0023 (0.0036) | 0.0005 (0.0009) | 0.0013 (0.0022) | 0.0009 (0.0021) | 0.0052 (0.0043) | 0.0001 (0.0029) | 0.0005 (0.0006) |
| 21q | **0.0058 (0.0027)** | 0.0011 (0.0028) | –0.0004 (0.0007) | **0.0035 (0.0017)** | 0.0066 (0.0034) | 0.0020 (0.0020) | 0.0021 (0.0018) | 0.0004 (0.0004) |
| 22q | 0.0052 (0.0030) | 0.0044 (0.0026) | 0.0009 (0.0006) | 0.0013 (0.0013) | –0.0007 (0.0021) | 0.0005 (0.0019) | 0.0011 (0.0020) | **0.0008 (0.0004)** |
| Any | –0.0014 (0.0008) | **0.0012 (0.0006)** | 0.0003 (0.0002) | **0.0022 (0.0005)** | **0.0020 (0.0005)** | 0.0004 (0.0006) | 0.0009 (0.0005) | **0.0009 (0.0001)** |

This table provides numerical data plotted in Fig. 3b. Units for polygenic scores are standard deviations for blood count traits; for Y loss, polygenic scores were computed on a 0/1 binary trait (modeled additively). Mean changes in polygenic scores reaching nominal significance ($P<0.05$ before multiple hypothesis correction) are indicated in bold; those that reached significance at FDR 0.05 or after Bonferroni correction are indicated in Fig. 3b.

**Supplementary Table 14. Mean changes in polygenic scores for six non-proliferation-related control traits produced by CNN-LOH mutations.**

| Arm | Height | BMI | Bone mineral density | FEV1/FVC | Blood pressure (systolic) | Blood pressure (diastolic) |
|---|---|---|---|---|---|---|
| 1p | –0.0027 (0.0031) | –0.0033 (0.0019) | –0.0012 (0.0028) | *–0.0055 (0.0021)* | –0.0022 (0.0021) | –0.0009 (0.0019) |
| 1q | 0.0015 (0.0039) | –0.0001 (0.0022) | 0.0017 (0.0030) | –0.0006 (0.0025) | *0.0041 (0.0019)* | 0.0038 (0.0021) |
| 2p | 0.0047 (0.0072) | –0.0065 (0.0048) | 0.0035 (0.0057) | –0.0070 (0.0046) | –0.0005 (0.0038) | –0.0017 (0.0040) |
| 2q | 0.0042 (0.0075) | 0.0017 (0.0043) | –0.0065 (0.0060) | –0.0054 (0.0051) | *0.0092 (0.0039)* | *0.0082 (0.0039)* |
| 3p | –0.0063 (0.0056) | 0.0023 (0.0039) | 0.0057 (0.0045) | –0.0010 (0.0043) | *0.0081 (0.0037)* | 0.0047 (0.0034) |
| 3q | 0.0082 (0.0066) | *0.0083 (0.0042)* | –0.0026 (0.0041) | –0.0061 (0.0039) | –0.0036 (0.0034) | –0.0017 (0.0036) |
| 4p | 0.0105 (0.0090) | 0.0049 (0.0052) | 0.0010 (0.0069) | –0.0036 (0.0039) | –0.0041 (0.0044) | –0.0011 (0.0041) |
| 4q | 0.0049 (0.0065) | 0.0007 (0.0034) | –0.0035 (0.0044) | –0.0007 (0.0053) | 0.0027 (0.0039) | 0.0019 (0.0038) |
| 5p | –0.0036 (0.0061) | 0.0021 (0.0049) | –0.0010 (0.0047) | –0.0044 (0.0046) | 0.0003 (0.0045) | 0.0034 (0.0038) |
| 5q | 0.0107 (0.0081) | 0.0004 (0.0044) | –0.0028 (0.0048) | –0.0074 (0.0059) | –0.0057 (0.0036) | –0.0023 (0.0038) |
| 6p | –0.0026 (0.0046) | –0.0016 (0.0025) | –0.0035 (0.0030) | *0.0071 (0.0034)* | 0.0027 (0.0021) | 0.0011 (0.0022) |
| 6q | 0.0087 (0.0095) | 0.0005 (0.0044) | 0.0077 (0.0104) | 0.0001 (0.0059) | 0.0075 (0.0040) | *0.0111 (0.0041)* |
| 7p | –0.0065 (0.0089) | *0.0109 (0.0043)* | –0.0013 (0.0075) | –0.0038 (0.0043) | –0.0038 (0.0049) | –0.0028 (0.0042) |
| 7q | 0.0015 (0.0057) | –0.0068 (0.0037) | –0.0008 (0.0088) | –0.0016 (0.0035) | –0.0015 (0.0036) | 0.0016 (0.0036) |
| 8p | 0.0017 (0.0066) | –0.0022 (0.0039) | –0.0017 (0.0047) | 0.0009 (0.0038) | 0.0039 (0.0046) | 0.0032 (0.0045) |
| 8q | 0.0096 (0.0079) | –0.0045 (0.0040) | 0.0009 (0.0049) | *–0.0085 (0.0037)* | 0.0070 (0.0044) | 0.0071 (0.0040) |
| 9p | 0.0039 (0.0025) | –0.0010 (0.0020) | –0.0014 (0.0021) | –0.0012 (0.0019) | –0.0017 (0.0016) | –0.0014 (0.0014) |
| 9q | 0.0035 (0.0050) | 0.0041 (0.0024) | –0.0051 (0.0028) | 0.0004 (0.0025) | 0.0001 (0.0019) | –0.0003 (0.0021) |
| 10p | 0.0014 (0.0073) | –0.0007 (0.0041) | 0.0149 (0.0080) | 0.0019 (0.0066) | 0.0084 (0.0048) | 0.0076 (0.0053) |
| 10q | –0.0105 (0.0073) | 0.0028 (0.0056) | 0.0117 (0.0078) | –0.0031 (0.0054) | –0.0045 (0.0058) | –0.0051 (0.0060) |
| 11p | –0.0002 (0.0028) | –0.0016 (0.0019) | –0.0037 (0.0024) | *–0.0035 (0.0016)* | –0.0009 (0.0021) | –0.0014 (0.0021) |
| 11q | 0.0020 (0.0027) | –0.0017 (0.0021) | –0.0017 (0.0033) | 0.0016 (0.0020) | –0.0002 (0.0021) | *0.0050 (0.0020)* |
| 12p | 0.0033 (0.0083) | –0.0019 (0.0046) | 0.0003 (0.0061) | 0.0059 (0.0052) | –0.0014 (0.0041) | 0.0027 (0.0044) |
| 12q | 0.0022 (0.0059) | 0.0009 (0.0031) | –0.0001 (0.0037) | 0.0017 (0.0029) | 0.0026 (0.0031) | *0.0067 (0.0032)* |
| 13q | –0.0016 (0.0031) | 0.0016 (0.0022) | –0.0048 (0.0028) | –0.0011 (0.0021) | –0.0007 (0.0017) | –0.0024 (0.0018) |
| 14q | 0.0005 (0.0028) | –0.0006 (0.0017) | 0.0019 (0.0020) | –0.0023 (0.0017) | 0.0008 (0.0015) | 0.0004 (0.0014) |
| 15q | –0.0079 (0.0047) | 0.0021 (0.0020) | –0.0036 (0.0024) | –0.0025 (0.0029) | 0.0006 (0.0020) | 0.0002 (0.0023) |
| 16p | 0.0013 (0.0035) | *–0.0064 (0.0025)* | 0.0005 (0.0026) | –0.0035 (0.0019) | –0.0034 (0.0020) | –0.0026 (0.0020) |
| 16q | 0.0015 (0.0043) | 0.0028 (0.0033) | *0.0123 (0.0035)* | *0.0086 (0.0029)* | 0.0014 (0.0022) | *0.0053 (0.0022)* |
| 17p | 0.0034 (0.0059) | 0.0024 (0.0030) | 0.0004 (0.0049) | 0.0009 (0.0034) | –0.0050 (0.0031) | 0.0008 (0.0028) |
| 17q | 0.0053 (0.0047) | –0.0027 (0.0021) | 0.0000 (0.0036) | –0.0020 (0.0026) | –0.0041 (0.0023) | –0.0028 (0.0020) |
| 18p | –0.0076 (0.0058) | 0.0087 (0.0048) | 0.0025 (0.0077) | –0.0045 (0.0052) | 0.0003 (0.0051) | –0.0032 (0.0039) |
| 18q | –0.0042 (0.0049) | –0.0034 (0.0046) | –0.0019 (0.0036) | –0.0014 (0.0032) | –0.0012 (0.0034) | –0.0026 (0.0034) |
| 19p | 0.0005 (0.0048) | –0.0026 (0.0025) | –0.0044 (0.0038) | –0.0016 (0.0023) | –0.0042 (0.0030) | –0.0021 (0.0028) |
| 19q | *0.0091 (0.0034)* | *–0.0046 (0.0022)* | 0.0012 (0.0027) | 0.0003 (0.0024) | 0.0007 (0.0017) | –0.0000 (0.0019) |
| 20p | 0.0019 (0.0069) | *0.0085 (0.0035)* | –0.0023 (0.0073) | –0.0053 (0.0038) | 0.0016 (0.0040) | 0.0018 (0.0046) |
| 20q | 0.0030 (0.0043) | –0.0009 (0.0026) | 0.0014 (0.0028) | –0.0027 (0.0026) | –0.0031 (0.0028) | –0.0034 (0.0031) |
| 21q | –0.0035 (0.0032) | 0.0016 (0.0020) | –0.0041 (0.0031) | *0.0038 (0.0019)* | 0.0020 (0.0019) | 0.0039 (0.0021) |
| 22q | 0.0004 (0.0023) | –0.0025 (0.0014) | –0.0005 (0.0024) | 0.0007 (0.0017) | 0.0004 (0.0012) | –0.0005 (0.0011) |
| Any | 0.0009 (0.0008) | –0.0005 (0.0005) | –0.0007 (0.0006) | *–0.0013 (0.0005)* | 0.0001 (0.0004) | 0.0007 (0.0004) |

This table is the analog of Supplementary Table 13 for polygenic scores computed for six highly heritable, polygenic non-blood-cell traits [44]. These traits serve as controls, as variants influencing these traits are not typically expected to affect cell proliferation. Units are standard deviations. Mean changes in polygenic scores reaching nominal significance ($P<0.05$ before multiple hypothesis correction) are indicated in italics; no changes were significant after Bonferroni correction.

**Supplementary Table 15. Accuracy of predicting CNN-LOH directionality using genetic risk.**

| | | CNN-LOH-associated alleles only | | | Polygenic scores (for blood traits) | | | Both | | |
|---|---|---|---|---|---|---|---|---|---|---|
| Arm | Events | Pred. acc. (s.e.) | Pred. *R* (95% CI) | *P* | Pred. acc. (s.e.) | Pred. *R* (95% CI) | *P* | Pred. acc. (s.e.) | Pred. *R* (95% CI) | *P* |
| 1p | 927 | 0.639 (0.007) | 0.523 (0.474,0.568) | $2\times10^{-66}$ | 0.590 (0.016) | 0.338 (0.279,0.394) | $1.8\times10^{-26}$ | 0.636 (0.010) | 0.518 (0.470,0.564) | $3.7\times10^{-65}$ |
| 1q | 694 | 0.505 (0.002) | 0.080 (0.006,0.154) | 0.017 | 0.535 (0.018) | 0.023 (-0.051,0.097) | 0.27 | 0.550 (0.017) | 0.088 (0.013,0.161) | 0.01 |
| 2p | 169 | – | – | – | 0.556 (0.034) | 0.123 (-0.028,0.269) | 0.055 | 0.556 (0.034) | 0.123 (-0.028,0.269) | 0.055 |
| 2q | 205 | – | – | – | 0.605 (0.034) | 0.227 (0.093,0.353) | 0.00053 | 0.605 (0.034) | 0.227 (0.093,0.353) | 0.00053 |
| 3p | 164 | – | – | – | 0.497 (0.033) | 0.002 (-0.151,0.155) | 0.49 | 0.497 (0.033) | 0.002 (-0.151,0.155) | 0.49 |
| 5p | 71 | – | – | – | 0.704 (0.055) | 0.417 (0.203,0.592) | 0.00015 | 0.704 (0.055) | 0.417 (0.203,0.592) | 0.00015 |
| 5q | 162 | – | – | – | 0.574 (0.039) | 0.178 (0.025,0.324) | 0.012 | 0.574 (0.039) | 0.178 (0.025,0.324) | 0.012 |
| 8q | 134 | 0.526 (0.010) | 0.229 (0.061,0.383) | 0.0039 | – | – | – | 0.526 (0.010) | 0.229 (0.061,0.383) | 0.0039 |
| 9p | 386 | 0.680 (0.016) | 0.501 (0.422,0.572) | $3.6\times10^{-26}$ | 0.505 (0.025) | 0.009 (-0.090,0.109) | 0.43 | 0.680 (0.016) | 0.501 (0.422,0.572) | $3.6\times10^{-26}$ |
| 11q | 647 | 0.543 (0.006) | 0.294 (0.222,0.363) | $1.1\times10^{-14}$ | 0.575 (0.019) | 0.189 (0.113,0.262) | $6.6\times10^{-7}$ | 0.615 (0.019) | 0.351 (0.282,0.417) | $1.6\times10^{-20}$ |
| 12q | 302 | 0.523 (0.006) | 0.216 (0.105,0.321) | $7.9\times10^{-5}$ | 0.603 (0.028) | 0.269 (0.161,0.371) | $1\times10^{-6}$ | 0.596 (0.028) | 0.264 (0.156,0.366) | $1.7\times10^{-6}$ |
| 14q | 956 | 0.569 (0.012) | 0.194 (0.133,0.255) | $6.8\times10^{-10}$ | 0.544 (0.016) | 0.155 (0.092,0.216) | $7.7\times10^{-7}$ | 0.559 (0.014) | 0.171 (0.109,0.232) | $4.8\times10^{-8}$ |
| 15q | 638 | 0.612 (0.009) | 0.460 (0.397,0.519) | $4.3\times10^{-35}$ | 0.494 (0.011) | -0.045 (-0.123,0.032) | 0.87 | 0.601 (0.010) | 0.453 (0.389,0.513) | $6.3\times10^{-34}$ |
| 18q | 127 | – | – | – | 0.535 (0.044) | 0.170 (-0.005,0.334) | 0.028 | 0.535 (0.044) | 0.170 (-0.005,0.334) | 0.028 |

This table provides numerical data plotted in Fig. 3c. CNN-LOH directions were predicted using: (i) only CNN-LOH-associated alleles on affected chromosomal segments (for chromosome arms containing at least one association; Table 1); (ii) polygenic score differentials on affected chromosomal segments; (iii) both CNN-LOH-associated alleles and polygenic scores.

For each chromosome arm with at least one available predictor (Methods), prediction accuracy was computed as the fraction of predicted CNN-LOH directions (hard-called) that matched observed CNN-LOH directions. Prediction *R* was computed as the correlation between predicted CNN-LOH directions (continuous-valued, as output by the linear predictor) and observed CNN-LOH directions. Predictive performance was assessed using 10-fold cross-validation, and both accuracy and *R* metrics (and standard errors and 95% CIs) were computed over a merge of all held-out folds. *P*-values were computed using a one-sided test for Pearson correlation $R>0$.

#### Supplementary Table 16. Risk increase for incident cancers conferred by mCAs.

(a) Analyses restricted to individuals with normal blood counts at assessment

| | CLL | | MPN | | Any blood cancer | |
|---|---|---|---|---|---|---|
| mCA | *P* | OR (95% CI) | *P* | OR (95% CI) | *P* | OR (95% CI) |
| 1+ | 1 | 0 (0–280) | 1 | 0 (0–568) | 0.0073 | 16.8 (1.93–66.6) |
| 1p– | 1 | 0 (0–1.1e+03) | 1 | 0 (0–1.62e+03) | 0.037 | 30 (0.67–224) |
| 1q– | 1 | 0 (0–621) | 1 | 0 (0–1.28e+03) | 1 | 0 (0–70.3) |
| 1p= | 1 | 0 (0–25.7) | 0.084 | 11.6 (0.29–67.6) | 0.17 | 2.15 (0.44–6.35) |
| 1q= | 1 | 0 (0–32.3) | 1 | 0 (0–53.1) | 0.32 | 1.76 (0.21–6.44) |
| 2p– | 1 | 0 (0–164) | 1 | 0 (0–236) | 1 | 0 (0–16.1) |
| 2q– | 1 | 0 (0–478) | 1 | 0 (0–740) | 1 | 0 (0–46) |
| 2p= | 1 | 0 (0–158) | 1 | 0 (0–280) | 1 | 0 (0–16.6) |
| 2q= | 1 | 0 (0–123) | 1 | 0 (0–217) | 1 | 0 (0–12.6) |
| 3+ | 0.00029 | 85.8 (9.96–337) | 1 | 0 (0–249) | 0.00013 | 17.1 (4.47–46.9) |
| 3p– | 1 | 0 (0–681) | 1 | 0 (0–1.41e+03) | 1 | 0 (0–73.4) |
| 3p= | 1 | 0 (0–118) | 1 | 0 (0–202) | 1 | 0 (0–12.2) |
| 3q= | 1 | 0 (0–167) | 1 | 0 (0–294) | 0.21 | 4.32 (0.11–25.2) |
| 4q– | 1 | 0 (0–125) | 1 | 0 (0–224) | 0.032 | 7.42 (0.88–28.1) |
| 4p= | 1 | 0 (0–453) | 1 | 0 (0–622) | 0.088 | 11.3 (0.27–70.3) |
| 4q= | 1 | 0 (0–88.2) | 1 | 0 (0–158) | 0.0011 | 9.4 (2.5–25) |
| 5+ | 1 | 0 (0–205) | 1 | 0 (0–336) | 0.16 | 5.86 (0.14–34.7) |
| 5q– | 1 | 0 (0–134) | 1 | 0 (0–265) | 0.003 | 11 (2.21–33.6) |
| 5q= | 1 | 0 (0–155) | 1 | 0 (0–261) | 1 | 0 (0–16.1) |
| 6+ | 1 | 0 (0–1.1e+03) | 1 | 0 (0–1.41e+03) | 1 | 0 (0–111) |
| 6q– | 1 | 0 (0–367) | 1 | 0 (0–570) | 0.1 | 9.56 (0.23–58.7) |
| 6p= | 1 | 0 (0–74.8) | 1 | 0 (0–118) | 0.094 | 3.96 (0.47–14.7) |
| 6q= | 1 | 0 (0–227) | 1 | 0 (0–377) | 1 | 0 (0–23.5) |
| 7p– | 1 | 0 (0–546) | 1 | 0 (0–1.23e+03) | 1 | 0 (0–57) |
| 7q– | 1 | 0 (0–190) | 1 | 0 (0–298) | 0.019 | 9.77 (1.15–37.5) |
| 7p= | 1 | 0 (0–237) | 1 | 0 (0–369) | 1 | 0 (0–23.3) |
| 7q= | 1 | 0 (0–170) | 0.013 | 81.3 (1.99–497) | $9.8\times10^{-5}$ | 18.2 (4.76–49.9) |
| 8+ | 0.023 | 44.4 (1.09–263) | 1 | 0 (0–322) | $2.3\times10^{-6}$ | 26.7 (8.24–67) |
| 8p– | 0.0072 | 147 (3.49–979) | 1 | 0 (0–808) | 0.066 | 15.4 (0.37–98) |
| 8p= | 1 | 0 (0–566) | 1 | 0 (0–1.16e+03) | 1 | 0 (0–54.4) |
| 8q= | 1 | 0 (0–175) | 1 | 0 (0–283) | 1 | 0 (0–17.9) |
| 9+ | 0.019 | 54.3 (1.33–329) | 1 | 0 (0–402) | 0.15 | 6.3 (0.15–37.8) |
| 9q– | 1 | 0 (0–561) | 1 | 0 (0–1.08e+03) | 1 | 0 (0–71) |
| 9p= | 1 | 0 (0–57.8) | $3.6\times10^{-13}$ | 260 (89.4–631) | $8.2\times10^{-6}$ | 13.8 (4.91–31.1) |
| 9q= | 1 | 0 (0–46.8) | 1 | 0 (0–83.7) | 1 | 0 (0–4.81) |
| 10q– | 1 | 0 (0–75.2) | 1 | 0 (0–122) | 1 | 0 (0–7.28) |
| 10p= | 1 | 0 (0–393) | 1 | 0 (0–726) | 1 | 0 (0–39) |
| 10q= | 1 | 0 (0–199) | 1 | 0 (0–338) | 1 | 0 (0–20.1) |
| 11p– | 1 | 0 (0–394) | 1 | 0 (0–698) | 1 | 0 (0–48) |
| 11q– | 0.045 | 22.1 (0.55–130) | 1 | 0 (0–147) | $9.2\times10^{-5}$ | 11.9 (3.75–28.8) |
| 11p= | 1 | 0 (0–32.4) | 1 | 0 (0–53.3) | 0.63 | 0 (0–3.32) |
| 11q= | 0.092 | 10.6 (0.26–60.9) | 0.056 | 17.7 (0.44–104) | 0.0004 | 6.6 (2.39–14.7) |
| 12+ | $5.1\times10^{-22}$ | 149 (72.9–278) | 0.059 | 16.8 (0.41–98.6) | $2.9\times10^{-20}$ | 23.3 (13.9–37.3) |
| 12p– | 1 | 0 (0–788) | 1 | 0 (0–1.71e+03) | 1 | 0 (0–89.4) |
| 12q– | 1 | 0 (0–601) | 1 | 0 (0–821) | 1 | 0 (0–56.5) |
| 12p= | 1 | 0 (0–455) | 1 | 0 (0–793) | 1 | 0 (0–47.5) |
| 12q= | 1 | 0 (0–80.2) | 1 | 0 (0–141) | 1 | 0 (0–8.3) |
| 13q– | $5.1\times10^{-13}$ | 127 (48.3–280) | 1 | 0 (0–77.3) | $1.8\times10^{-7}$ | 14.4 (6.08–29.5) |
| 13q= | $8.2\times10^{-5}$ | 38.4 (7.7–117) | 1 | 0 (0–68.6) | 0.047 | 3.81 (0.78–11.4) |
| 14+ | 1 | 0 (0–73.1) | 1 | 0 (0–135) | 0.39 | 2.07 (0.05–11.9) |
| 14q– | 0.00017 | 115 (13.2–456) | 1 | 0 (0–294) | 0.00074 | 18.5 (3.63–59.2) |
| 14q= | 1 | 0 (0–23.5) | 1 | 0 (0–40) | 1 | 0.65 (0.02–3.64) |
| 15+ | 1 | 0 (0–46.8) | 1 | 0 (0–93) | 0.55 | 1.27 (0.03–7.22) |
| 15q– | 1 | 0 (0–943) | 1 | 0 (0–1.59e+03) | 1 | 0 (0–106) |
| 15q= | 1 | 0 (0–37.1) | 1 | 0 (0–66.2) | 0.62 | 1.03 (0.03–5.83) |
| 16p– | 1 | 0 (0–142) | 1 | 0 (0–259) | 1 | 0 (0–13.9) |
| 16q– | 1 | 0 (0–438) | 1 | 0 (0–850) | 1 | 0 (0–50.8) |
| 16p= | 1 | 0 (0–63.9) | 1 | 0 (0–110) | 0.11 | 3.52 (0.42–13) |
| 16q= | 1 | 0 (0–80.4) | 0.029 | 35 (0.86–207) | 0.38 | 2.13 (0.05–12.2) |
| 17p– | 1 | 0 (0–125) | 1 | 0 (0–172) | 0.27 | 3.16 (0.08–18.3) |
| 17q– | 1 | 0 (0–288) | 1 | 0 (0–544) | 0.13 | 7.51 (0.18–45.4) |
| 17p= | 1 | 0 (0–197) | 1 | 0 (0–348) | 1 | 0 (0–20.2) |
| 17q= | 1 | 0 (0–52.6) | 1 | 0 (0–90.3) | 0.51 | 1.42 (0.04–8.04) |
| 18+ | $6.7\times10^{-6}$ | 91 (18–288) | 1 | 0 (0–186) | $2.9\times10^{-5}$ | 15.3 (4.8–37.6) |
| 18p– | 1 | 0 (0–665) | 1 | 0 (0–1.41e+03) | 1 | 0 (0–77.6) |
| 18q= | 1 | 0 (0–233) | 1 | 0 (0–454) | 1 | 0 (0–25.3) |
| 19+ | 1 | 0 (0–687) | 1 | 0 (0–982) | 1 | 0 (0–72.9) |
| 19p= | 1 | 0 (0–116) | 1 | 0 (0–209) | 0.28 | 3.13 (0.08–18.1) |
| 19q= | 1 | 0 (0–93) | 0.023 | 45.1 (1.11–265) | 0.058 | 5.28 (0.63–19.7) |
| 20q– | 1 | 0 (0–35.2) | 1 | 0 (0–61.1) | 0.021 | 3.94 (1.06–10.3) |
| 20p= | 1 | 0 (0–503) | 1 | 0 (0–796) | 1 | 0 (0–48.4) |
| 20q= | 1 | 0 (0–104) | 1 | 0 (0–165) | 0.31 | 2.78 (0.07–15.9) |
| 21+ | 1 | 0 (0–100) | 1 | 0 (0–170) | 0.31 | 2.75 (0.07–15.8) |
| 21q– | 1 | 0 (0–808) | 1 | 0 (0–1.55e+03) | 0.048 | 22.3 (0.51–154) |
| 21q= | 1 | 0 (0–127) | 1 | 0 (0–210) | 0.26 | 3.32 (0.08–19.2) |
| 22+ | 1 | 0 (0–88.3) | 1 | 0 (0–142) | 0.073 | 4.61 (0.55–17.2) |
| 22q– | 1 | 0 (0–256) | 1 | 0 (0–305) | 0.13 | 7.19 (0.18–43.4) |
| 22q= | 1 | 0 (0–48.9) | 1 | 0 (0–81.5) | 0.54 | 1.29 (0.03–7.28) |

This table provides numerical data plotted in Fig. 4a. Events were grouped by chromosome and copy number, with loss and CNN-LOH events subdivided by p-arm vs. q-arm; events observed in $\geq$30 individuals were tested for association with incident blood cancers (diagnosed $>$1 year after DNA collection in individuals with no previous cancer).

#### (b) Analyses with no restrictions on blood counts at assessment

| | CLL | | MPN | | Any blood cancer | |
|---|---|---|---|---|---|---|
| mCA | *P* | OR (95% CI) | *P* | OR (95% CI) | *P* | OR (95% CI) |
| 1+ | 1 | 0 (0–151) | 1 | 0 (0–250) | 0.00081 | 18 (3.52–57.8) |
| 1p− | 1 | 0 (0–441) | 1 | 0 (0–666) | 0.0027 | 29.6 (3.21–132) |
| 1q− | 1 | 0 (0–327) | 1 | 0 (0–597) | 0.092 | 11 (0.26–71.5) |
| 1p= | 0.24 | 3.59 (0.09–20.4) | 0.00087 | 16.9 (3.41–50.8) | 0.00079 | 4.27 (1.83–8.53) |
| 1q= | 1 | 0 (0–17.1) | 1 | 0 (0–25.6) | 0.66 | 1.35 (0.16–4.91) |
| 2p− | 0.053 | 18.8 (0.47–110) | 1 | 0 (0–117) | 0.31 | 2.76 (0.07–15.9) |
| 2q− | 1 | 0 (0–220) | 1 | 0 (0–365) | 1 | 0 (0–30.5) |
| 2p= | 1 | 0 (0–80.9) | 1 | 0 (0–126) | 1 | 0 (0–11.8) |
| 2q= | 1 | 0 (0–65.8) | 1 | 0 (0–98.4) | 1 | 0 (0–9.35) |
| 3+ | $1.9\times10^{-5}$ | 63.6 (12.6–200) | 1 | 0 (0–127) | $1.1\times10^{-7}$ | 20.9 (8.02–45.9) |
| 3p− | 1 | 0 (0–377) | 1 | 0 (0–614) | 1 | 0 (0–53.6) |
| 3p= | 1 | 0 (0–63.7) | 1 | 0 (0–99.4) | 1 | 0 (0–9.61) |
| 3q= | 1 | 0 (0–89.6) | 1 | 0 (0–141) | 0.26 | 3.4 (0.08–19.8) |
| 4q− | 0.059 | 16.8 (0.42–98.2) | 1 | 0 (0–105) | 0.0086 | 7.47 (1.5–22.8) |
| 4p= | 1 | 0 (0–239) | 1 | 0 (0–295) | 0.12 | 8.01 (0.19–49.1) |
| 4q= | 1 | 0 (0–44.8) | 1 | 0 (0–77.3) | 0.00039 | 8.62 (2.73–20.9) |
| 5+ | 1 | 0 (0–93.6) | 1 | 0 (0–157) | 0.24 | 3.72 (0.09–21.8) |
| 5q− | 0.054 | 18.2 (0.45–107) | 0.034 | 29.5 (0.73–172) | $3.1\times10^{-6}$ | 16.5 (5.84–37.6) |
| 5q= | 1 | 0 (0–80.3) | 1 | 0 (0–118) | 0.29 | 2.96 (0.07–17.1) |
| 6+ | 1 | 0 (0–514) | 1 | 0 (0–670) | 1 | 0 (0–71.7) |
| 6q− | 1 | 0 (0–168) | 1 | 0 (0–294) | 0.15 | 6.45 (0.16–39) |
| 6p= | 1 | 0 (0–39.3) | 1 | 0 (0–57.8) | 0.15 | 2.97 (0.36–11) |
| 6q= | 1 | 0 (0–124) | 1 | 0 (0–193) | 1 | 0 (0–18.2) |
| 7p− | 1 | 0 (0–282) | 1 | 0 (0–499) | 1 | 0 (0–46.8) |
| 7q− | 1 | 0 (0–87.1) | 1 | 0 (0–154) | 0.00028 | 13.7 (3.6–37.4) |
| 7p= | 1 | 0 (0–124) | 1 | 0 (0–195) | 1 | 0 (0–17.9) |
| 7q= | 1 | 0 (0–94.7) | 0.027 | 37.8 (0.93–221) | 0.00027 | 13.9 (3.66–37.9) |
| 8+ | 0.046 | 21.7 (0.54–127) | 1 | 0 (0–129) | $1.3\times10^{-6}$ | 19.4 (6.84–45) |
| 8p− | 0.016 | 64.9 (1.56–406) | 1 | 0 (0–415) | 0.1 | 9.91 (0.24–61.4) |
| 8p= | 1 | 0 (0–297) | 1 | 0 (0–473) | 1 | 0 (0–42) |
| 8q= | 1 | 0 (0–93) | 1 | 0 (0–138) | 1 | 0 (0–13.6) |
| 9+ | 0.036 | 27.8 (0.68–166) | 0.00026 | 90 (10.5–354) | 0.0024 | 12.1 (2.4–37.8) |
| 9q− | 1 | 0 (0–271) | 1 | 0 (0–399) | 1 | 0 (0–37.8) |
| 9p= | 1 | 0 (0–31.3) | $6.3\times10^{-52}$ | 402 (239–671) | $1.9\times10^{-29}$ | 33.5 (21.1–51.2) |
| 9q= | 1 | 0 (0–24) | 1 | 0 (0–38.8) | 0.63 | 0 (0–3.56) |
| 10q− | 1 | 0 (0–38.4) | 1 | 0 (0–54.4) | 1 | 0 (0–5.4) |
| 10p= | 1 | 0 (0–227) | 1 | 0 (0–341) | 1 | 0 (0–32.4) |
| 10q= | 1 | 0 (0–110) | 1 | 0 (0–161) | 1 | 0 (0–16) |
| 11p− | 1 | 0 (0–199) | 1 | 0 (0–297) | 1 | 0 (0–30.3) |
| 11q− | 0.00012 | 33.8 (6.79–103) | 1 | 0 (0–70.4) | $2.3\times10^{-8}$ | 15.1 (6.71–29.9) |
| 11p= | 1 | 0 (0–17.4) | 1 | 0 (0–26.1) | 0.41 | 0 (0–2.53) |
| 11q= | 0.17 | 5.52 (0.14–31.5) | 0.11 | 8.79 (0.22–50.5) | 0.00032 | 5.71 (2.26–12) |
| 12+ | $8.9\times10^{-42}$ | 142 (86.5–225) | 0.11 | 8.46 (0.21–48.6) | $4.8\times10^{-35}$ | 26.9 (18.1–38.9) |
| 12p− | 1 | 0 (0–431) | 1 | 0 (0–769) | 1 | 0 (0–66.2) |
| 12q− | 1 | 0 (0–263) | 0.011 | 98.6 (2.37–628) | 0.11 | 9.2 (0.22–57.4) |
| 12p= | 1 | 0 (0–227) | 1 | 0 (0–339) | 1 | 0 (0–32.8) |
| 12q= | 1 | 0 (0–41.9) | 1 | 0 (0–66.9) | 1 | 0 (0–6.27) |
| 13q− | $1.2\times10^{-54}$ | 212 (133–327) | 1 | 0 (0–38.4) | $6.1\times10^{-33}$ | 28.9 (19–42.6) |
| 13q= | $1.4\times10^{-11}$ | 49.5 (20.8–102) | 1 | 0 (0–34.4) | $4.7\times10^{-6}$ | 7.71 (3.47–14.9) |
| 14+ | 1 | 0 (0–37.5) | 1 | 0 (0–65.2) | 0.49 | 1.5 (0.04–8.55) |
| 14q− | $2.3\times10^{-9}$ | 109 (33.5–276) | 1 | 0 (0–139) | $8.7\times10^{-8}$ | 21.7 (8.31–47.9) |
| 14q= | 1 | 0 (0–12.2) | 0.18 | 5.14 (0.13–29.3) | 0.062 | 2.4 (0.77–5.67) |
| 15+ | 0.14 | 6.8 (0.17–39.4) | 1 | 0 (0–47.1) | 0.086 | 2.95 (0.6–8.84) |
| 15q− | 0.011 | 94.6 (2.21–640) | 1 | 0 (0–700) | 0.074 | 13.9 (0.33–93.1) |
| 15q= | 1 | 0 (0–20.1) | 1 | 0 (0–31.5) | 1 | 0.81 (0.02–4.54) |
| 16p− | 1 | 0 (0–73.1) | 1 | 0 (0–115) | 1 | 0 (0–10.6) |
| 16q− | 0.00018 | 115 (12.9–472) | 1 | 0 (0–362) | 0.0074 | 16.9 (1.91–69.5) |
| 16p= | 1 | 0 (0–32.7) | 1 | 0 (0–53) | 0.18 | 2.6 (0.31–9.57) |
| 16q= | 1 | 0 (0–42.6) | 0.057 | 17.4 (0.43–101) | 0.46 | 1.62 (0.04–9.25) |
| 17p− | 1 | 0 (0–56.2) | 1 | 0 (0–91.5) | 0.083 | 4.27 (0.51–16) |
| 17q− | 1 | 0 (0–154) | 1 | 0 (0–258) | 0.16 | 5.88 (0.14–35.5) |
| 17p= | 1 | 0 (0–107) | 1 | 0 (0–164) | 1 | 0 (0–15.3) |
| 17q= | 1 | 0 (0–28.6) | 1 | 0 (0–43.8) | 0.6 | 1.09 (0.03–6.19) |
| 18+ | $1\times10^{-6}$ | 58.9 (15.5–159) | 1 | 0 (0–92.3) | $8.1\times10^{-7}$ | 15.2 (5.91–32.9) |
| 18p− | 1 | 0 (0–330) | 1 | 0 (0–581) | 0.085 | 12.1 (0.28–80.1) |
| 18q= | 1 | 0 (0–121) | 1 | 0 (0–188) | 1 | 0 (0–17.6) |
| 19+ | 1 | 0 (0–301) | 1 | 0 (0–463) | 1 | 0 (0–42.6) |
| 19p= | 1 | 0 (0–61.9) | 1 | 0 (0–97) | 0.35 | 2.36 (0.06–13.5) |
| 19q= | 1 | 0 (0–50.4) | 0.047 | 21.1 (0.53–123) | 0.002 | 7.88 (2.1–20.9) |
| 20q− | 1 | 0 (0–18.7) | 1 | 0 (0–31) | 0.053 | 2.9 (0.78–7.54) |
| 20p= | 1 | 0 (0–235) | 1 | 0 (0–361) | 1 | 0 (0–33.6) |
| 20q= | 0.069 | 14.2 (0.35–82) | 1 | 0 (0–81.8) | 0.085 | 4.19 (0.5–15.6) |
| 21+ | 1 | 0 (0–52.9) | 1 | 0 (0–87.5) | 0.38 | 2.09 (0.05–12) |
| 21q− | 1 | 0 (0–343) | 1 | 0 (0–540) | 0.08 | 12.8 (0.3–82.9) |
| 21q= | 1 | 0 (0–66.2) | 1 | 0 (0–101) | 0.34 | 2.46 (0.06–14.2) |
| 22+ | 0.088 | 11 (0.27–63.2) | 1 | 0 (0–71.1) | 0.00045 | 8.31 (2.64–20.1) |
| 22q− | $6.4\times10^{-16}$ | 188 (75.5–404) | 0.027 | 37.8 (0.93–224) | $2.2\times10^{-12}$ | 34.9 (15.7–69.7) |
| 22q= | 1 | 0 (0–25.1) | 1 | 0 (0–40.5) | 1 | 0.96 (0.02–5.4) |

This table provides results of analogous analyses removing the restrictions we imposed on blood counts in our primary analyses (lymphocyte count $1\text{–}3.5\times10^9$/L, red cell count $<6.1\times10^{12}$/L for males and $<5.4\times10^{12}$/L for females, platelet count $<450\times10^9$/L, RBC distribution width $<15\%$).

**Supplementary Table 17. Risk increase for cardiovascular disease (MI/stroke) during 5–10-year follow-up conferred by mCAs.**

| Event | # carriers | # matched controls | MI/stroke OR (95% CI) | *P* |
|---|---|---|---|---|
| Any loss | 2651 | 63624 | 1.15 (0.91–1.46) | 0.24 |
| Any CNN-LOH | 6977 | 118609 | 0.96 (0.82–1.14) | 0.71 |
| Any gain | 1794 | 43056 | 0.69 (0.49–0.99) | 0.038 |
| Any mCA | 14861 | 163471 | 1.00 (0.89–1.12) | 0.98 |
| *DNMT3A* loss | 107 | 3317 | 1.59 (0.49–5.16) | 0.44 |
| *TET2* loss | 66 | 3234 | 0.75 (0.10–5.49) | 1 |
| *JAK2* CNN-LOH | 291 | 7566 | **2.37 (1.40–4.01)** | **0.0031** |

This table provides numerical data plotted in Fig. 4b. The number of controls varies across mosaic events because cases and controls for each event were matched for assessment year, age, sex, smoking, hypertension, BMI, and type 2 diabetes status. The case-control ratio was chosen independently for each event to optimize statistical power.

**Supplementary Table 18. Numbers of cases and controls for association tests with CNN-LOH mutations in *cis*.**

| Arm | Locus | $N_{case}$ | $N_{control}$ |
|---|---|---|---|
| 1p | *MPL* | 633 | 377674 |
| 1q | *FH* | 666 | 377674 |
| 8q | *NBN* | 76 | 379049 |
| 9p | *JAK2* | 394 | 378410 |
| 11q | *MRE11* | 520 | 378073 |
| 11q | *ATM* | 581 | 378073 |
| 12q | *SH2B3* | 250 | 378874 |
| 14q | *TCL1A* | 1021 | 378180 |
| 14q | *DLK1* | 1052 | 378180 |
| 15q | *TM2D3* | 605 | 378617 |

Sample sets were determined by first filtering on ancestry and relatedness, and then at each locus, defining cases to be individuals with a mosaic event spanning the locus (or within 4Mb of the locus) likely to be a CNN-LOH event (Methods). Individuals with likely CNN-LOH events on the same chromosome but not within 4Mb of the locus were excluded from association analyses.

**Supplementary Table 19. Associations of telomere length SNPs with mosaic chromosomal alterations on any autosome.**

| Locus | SNP | Chr | Position | Alleles | EAF | $\beta_{telo}$ (s.e.) | $P_{telo}$ | $\beta_{mCA}$ (s.e.) | $P_{mCA}$ |
|---|---|---|---|---|---|---|---|---|---|
| *TERC* | rs10936599 | 3 | 169492101 | T/C | 0.252 | –0.097 (0.008) | $2.5\times10^{-31}$ | –0.0024 (0.0005) | $1.1\times10^{-7}$ |
| *TERT* | rs2736100 | 5 | 1286516 | A/C | 0.514 | –0.078 (0.009) | $4.4\times10^{-19}$ | –0.0028 (0.0004) | $1.2\times10^{-12}$ |
| *NAF1* | rs7675998 | 4 | 164007820 | A/G | 0.217 | –0.074 (0.009) | $4.4\times10^{-16}$ | –0.0006 (0.0005) | $2.0\times10^{-1}$ |
| *OBFC1* | rs9420907 | 10 | 105676465 | A/C | 0.865 | –0.069 (0.010) | $6.9\times10^{-11}$ | –0.0019 (0.0006) | $1.1\times10^{-3}$ |
| *ZNF208* | rs8105767 | 19 | 22215441 | A/G | 0.709 | –0.048 (0.008) | $1.1\times10^{-9}$ | –0.0005 (0.0004) | $2.8\times10^{-1}$ |
| *RTEL1* | rs755017 | 20 | 62421622 | A/G | 0.869 | –0.062 (0.011) | $6.7\times10^{-9}$ | –0.0014 (0.0006) | $1.5\times10^{-2}$ |
| *ACYP2* | rs11125529 | 2 | 54475866 | C/A | 0.858 | –0.056 (0.010) | $4.5\times10^{-8}$ | 0.0002 (0.0006) | $7.9\times10^{-1}$ |

Results from BOLT-LMM [26, 44] analysis of the "any autosomal mCA" phenotype are reported for variants previously associated with telomere length [59]. Alleles, effect allele / other allele. EAF, effect allele frequency as reported by ref. [59]. $\beta_{telo}$ (s.e.) and $P_{telo}$, effect size and association $P$-value for telomere length reported by ref. [59]. $\beta_{mCA}$ (s.e.) and $P_{mCA}$, effect size and association $P$-value for presence of an mCA on any autosome.

#### Supplementary Table 20. Effects of known CLL GWAS variants on mosaic +12 and 13q LOH risk.

| | | | CLL, Law et al. [60] | | CLL, UK Biobank | | Mosaic +12 | | Mosaic 13q LOH | |
|---|---|---|---|---|---|---|---|---|---|---|
| Locus | hg19 bp | Risk allele | OR (95% CI) | *P* | OR (95% CI) | *P* | OR (95% CI) | *P* | OR (95% CI) | *P* |
| 2p22.2 | 37603801 | rs888096:A | 1.15 (1.09–1.21) | 5.2e-08 | 1.02 (0.91–1.14) | 0.74 | 1.04 (0.93–1.16) | 0.54 | 1.08 (0.98–1.18) | 0.12 |
| 2q13 | 111616619 | rs1002015:C | 1.30 (1.23–1.37) | 2.2e-23 | 1.18 (1.06–1.32) | 0.003 | 1.15 (1.03–1.29) | 0.016 | 1.15 (1.05–1.27) | 0.003 |
| 2q13 | 111831793 | rs58055674:C | 1.41 (1.32–1.50) | 2e-27 | 1.17 (1.02–1.33) | 0.027 | 1.17 (1.02–1.34) | 0.025 | 1.16 (1.03–1.30) | 0.011 |
| 2q13 | 111927379 | rs6708784:G | 1.30 (1.24–1.37) | 2.7e-25 | 1.40 (1.25–1.56) | 2.8e-09 | 1.22 (1.10–1.37) | 0.00033 | 1.26 (1.15–1.38) | 1.3e-06 |
| 2q33.1 | 202023949 | rs7558911:A | 1.18 (1.12–1.24) | 5.1e-11 | 1.26 (1.13–1.41) | 3.1e-05 | 1.09 (0.98–1.22) | 0.12 | 1.06 (0.97–1.16) | 0.2 |
| 2q37.1 | 231154012 | rs34004493:G | 1.39 (1.31–1.47) | 3.7e-32 | 1.40 (1.25–1.57) | 9.9e-09 | 1.30 (1.16–1.46) | 1.1e-05 | 1.33 (1.21–1.47) | 1.2e-08 |
| 2q37.3 | 242294913 | rs3755397:G | 1.32 (1.22–1.43) | 9.5e-12 | 1.32 (1.12–1.56) | 0.00081 | 1.14 (0.96–1.36) | 0.14 | 1.12 (0.96–1.29) | 0.14 |
| 3p24.1 | 27777779 | rs9880772:A | 1.16 (1.11–1.22) | 1.9e-09 | 1.16 (1.04–1.30) | 0.006 | 1.10 (0.98–1.23) | 0.096 | 1.12 (1.02–1.23) | 0.014 |
| 3q26.2 | 169497585 | rs1317082:A | 1.19 (1.12–1.26) | 5.8e-09 | 1.21 (1.06–1.38) | 0.0049 | 1.12 (0.98–1.28) | 0.09 | 1.40 (1.25–1.58) | 1.8e-08 |
| 3q28 | 188128794 | rs73192661:C | 1.13 (1.07–1.19) | 1.7e-06 | 1.03 (0.93–1.15) | 0.56 | 1.04 (0.93–1.16) | 0.54 | 1.07 (0.97–1.17) | 0.18 |
| 4q25 | 109025865 | rs7690934:C | 1.16 (1.11–1.22) | 6.1e-09 | 1.19 (1.06–1.33) | 0.0024 | 1.19 (1.06–1.33) | 0.0027 | 1.00 (0.91–1.10) | 1 |
| 4q26 | 114698696 | rs1476569:G | 1.14 (1.08–1.20) | 4.5e-06 | 1.04 (0.93–1.18) | 0.48 | 1.11 (0.99–1.26) | 0.083 | 1.08 (0.98–1.20) | 0.14 |
| 5p15.33 | 1285974 | rs7705526:A | 1.18 (1.12–1.25) | 5.9e-10 | 1.19 (1.06–1.33) | 0.0028 | 1.11 (0.99–1.24) | 0.081 | 1.20 (1.09–1.31) | 0.00025 |
| 5p15.33 | 1321873 | rs10073340:T | 1.13 (1.06–1.20) | 0.00028 | 1.21 (1.06–1.39) | 0.006 | 1.15 (1.00–1.33) | 0.045 | 1.13 (1.00–1.27) | 0.044 |
| 6p25.3 | 412802 | rs9392504:A | 1.33 (1.26–1.40) | 9.8e-29 | 1.34 (1.20–1.50) | 1.8e-07 | 1.07 (0.96–1.19) | 0.25 | 1.19 (1.08–1.30) | 0.0003 |
| 6p25.2 | 2969278 | rs73718779:T | 1.14 (1.06–1.23) | 0.0007 | 1.18 (1.00–1.39) | 0.054 | 1.08 (0.91–1.28) | 0.4 | 1.00 (0.86–1.16) | 0.97 |
| 6p21.32 | 32578127 | rs9271176:G | 1.29 (1.22–1.36) | 3.2e-20 | 1.33 (1.18–1.50) | 4.4e-06 | 1.10 (0.98–1.24) | 0.11 | 1.18 (1.07–1.31) | 0.0012 |
| 6p21.31 | 33546930 | rs210143:C | 1.26 (1.19–1.33) | 5.8e-16 | 1.01 (0.90–1.14) | 0.81 | 1.22 (1.07–1.38) | 0.0021 | 1.00 (0.91–1.11) | 0.99 |
| 6q25.2 | 154471225 | rs4869818:G | 1.15 (1.09–1.21) | 4.1e-08 | 1.16 (1.04–1.29) | 0.0093 | 1.13 (1.01–1.26) | 0.029 | 0.95 (0.87–1.04) | 0.3 |
| 7q31.33 | 124392512 | rs2267708:T | 1.16 (1.10–1.22) | 8.6e-09 | 1.12 (1.00–1.24) | 0.047 | 0.98 (0.88–1.10) | 0.74 | 1.12 (1.02–1.23) | 0.015 |
| 8q22.3 | 103577865 | rs2511713:G | 1.17 (1.10–1.23) | 6e-08 | 1.14 (1.01–1.28) | 0.032 | 1.16 (1.03–1.31) | 0.016 | 1.11 (1.00–1.23) | 0.046 |
| 8q24.21 | 128200971 | rs2466029:G | 1.23 (1.17–1.30) | 7.5e-16 | 1.16 (1.03–1.29) | 0.0098 | 1.10 (0.98–1.23) | 0.091 | 1.09 (0.99–1.19) | 0.081 |
| 9p21.3 | 22206987 | rs1679013:C | 1.16 (1.10–1.22) | 2.2e-08 | 1.18 (1.05–1.31) | 0.0038 | 1.12 (1.00–1.25) | 0.05 | 1.21 (1.10–1.33) | 7.4e-05 |
| 10q23.31 | 90752018 | rs6586163:A | 1.23 (1.17–1.29) | 1.1e-15 | 1.31 (1.18–1.47) | 9.2e-07 | 1.30 (1.17–1.46) | 3.2e-06 | 1.20 (1.10–1.32) | 8.5e-05 |
| 11p15.5 | 2321650 | rs2651823:A | 1.18 (1.13–1.25) | 5.2e-11 | 1.20 (1.07–1.33) | 0.0012 | 0.95 (0.85–1.06) | 0.38 | 1.26 (1.15–1.38) | 6.4e-07 |
| 11q24.1 | 123355391 | rs35923643:G | 1.63 (1.53–1.72) | 4.3e-58 | 1.58 (1.40–1.78) | 1.3e-13 | 1.17 (1.02–1.33) | 0.024 | 1.44 (1.30–1.60) | 8e-12 |
| 12q24.13 | 113381376 | rs6489882:G | 1.16 (1.10–1.22) | 4.8e-08 | 1.10 (0.98–1.23) | 0.099 | 1.13 (1.01–1.27) | 0.031 | 1.06 (0.97–1.17) | 0.2 |
| 15q15.1 | 40403657 | rs8024033:C | 1.26 (1.20–1.32) | 7.1e-19 | 1.32 (1.18–1.47) | 6.9e-07 | 1.19 (1.07–1.33) | 0.0016 | 1.12 (1.02–1.23) | 0.013 |
| 15q21.3 | 56777691 | rs142215530:G | 1.39 (1.29–1.50) | 2.5e-18 | 1.35 (1.16–1.57) | 0.0001 | 1.36 (1.17–1.59) | 7.9e-05 | 1.26 (1.10–1.43) | 0.00075 |
| 15q23 | 70020525 | rs11637565:G | 1.35 (1.28–1.42) | 2e-31 | 1.34 (1.20–1.49) | 1.4e-07 | 1.03 (0.92–1.15) | 0.64 | 1.16 (1.06–1.27) | 0.0019 |
| 15q25.2 | 83237899 | rs17356118:A | 1.12 (1.05–1.19) | 0.00025 | 1.06 (0.93–1.21) | 0.38 | 1.10 (0.96–1.26) | 0.16 | 1.08 (0.97–1.21) | 0.18 |
| 16q24.1 | 85973866 | rs305065:C | 1.16 (1.10–1.22) | 7.6e-08 | 0.98 (0.88–1.10) | 0.76 | 1.00 (0.89–1.12) | 0.96 | 1.17 (1.06–1.29) | 0.0017 |
| 16q24.1 | 85928621 | rs391855:A | 1.34 (1.27–1.41) | 1.3e-28 | 1.28 (1.14–1.43) | 1.7e-05 | 1.17 (1.05–1.31) | 0.0058 | 1.13 (1.03–1.24) | 0.012 |
| 18q21.32 | 57622287 | rs4368253:C | 1.17 (1.11–1.24) | 1.3e-08 | 1.11 (0.98–1.25) | 0.091 | 1.21 (1.07–1.37) | 0.002 | 1.00 (0.91–1.10) | 1 |
| 18q21.33 | 60788745 | rs77551289:A | 1.37 (1.25–1.50) | 1.8e-11 | 1.26 (1.03–1.53) | 0.026 | 1.15 (0.94–1.40) | 0.17 | 1.18 (1.00–1.39) | 0.049 |
| 18q21.33 | 60793921 | rs4987852:C | 1.32 (1.20–1.44) | 4.7e-09 | 1.20 (0.98–1.46) | 0.076 | 1.29 (1.06–1.57) | 0.0099 | 1.13 (0.95–1.34) | 0.17 |
| 19q13.3 | 47176752 | rs874460:C | 1.24 (1.15–1.34) | 3.4e-08 | 1.29 (1.08–1.55) | 0.0042 | 0.89 (0.76–1.04) | 0.14 | 1.16 (1.00–1.34) | 0.047 |
| 1p36.11 | 23943735 | rs34676223:C | 1.19 (1.14–1.25) | 5e-13 | 1.09 (0.97–1.24) | 0.14 | 0.96 (0.85–1.08) | 0.47 | 1.14 (1.03–1.26) | 0.013 |
| 1q42.13 | 228880296 | rs41271473:G | 1.19 (1.13–1.26) | 1.1e-10 | 1.04 (0.91–1.19) | 0.6 | 1.19 (1.03–1.38) | 0.018 | 1.02 (0.91–1.15) | 0.71 |
| 4q24 | 102741002 | rs71597109:C | 1.17 (1.11–1.22) | 1.4e-10 | 1.26 (1.11–1.42) | 0.00023 | 1.00 (0.89–1.13) | 0.98 | 1.20 (1.08–1.32) | 0.00065 |
| 4q35.1 | 185254772 | rs57214277:T | 1.13 (1.08–1.18) | 3.7e-08 | 1.06 (0.95–1.18) | 0.33 | 1.04 (0.93–1.16) | 0.51 | 1.10 (1.00–1.21) | 0.038 |
| 6p21.31 | 34616322 | rs3800461:C | 1.20 (1.13–1.28) | 2e-08 | 1.29 (1.10–1.51) | 0.0013 | 1.27 (1.08–1.49) | 0.0033 | 1.21 (1.06–1.38) | 0.0053 |
| 11q23.2 | 113517203 | rs61904987:T | 1.24 (1.16–1.32) | 2.5e-11 | 1.19 (1.02–1.38) | 0.025 | 1.03 (0.88–1.21) | 0.73 | 1.22 (1.07–1.38) | 0.0026 |
| 18q21.1 | 47843534 | rs1036935:A | 1.15 (1.10–1.21) | 3.3e-08 | 0.98 (0.86–1.12) | 0.77 | 1.14 (1.00–1.30) | 0.046 | 0.99 (0.89–1.11) | 0.91 |
| 19p13.3 | 4069119 | rs7254272:A | 1.17 (1.10–1.23) | 4.7e-08 | 0.96 (0.83–1.11) | 0.58 | 1.00 (0.86–1.15) | 0.97 | 1.10 (0.98–1.24) | 0.099 |
| 22q13.33 | 50971266 | rs140522:T | 1.15 (1.10–1.20) | 2.7e-09 | 1.21 (1.08–1.35) | 0.001 | 0.95 (0.84–1.06) | 0.35 | 1.18 (1.08–1.30) | 0.0005 |

This table provides numerical data plotted in Supplementary Fig. 37 for the 46 lead SNPs reported in Supplementary Table 3 and Table 1 of Law et al. [60]. Details of analyses are provided in the Supplementary Note.
